# Comparative transcriptomics of seed nourishing tissues: uncovering conserved and divergent pathways in seed plants

**DOI:** 10.1101/2023.11.30.569347

**Authors:** Ana Marcela Florez-Rueda, Célia M. Miguel, Duarte D. Figueiredo

## Abstract

The evolutionary and ecological success of spermatophytes is intrinsically linked to the seed habit, which provides a protective environment for the initial development of the new generation. This environment includes an ephemeral nourishing tissue that supports embryo growth. In gymnosperms this tissue originates from the asexual proliferation of the maternal megagametophyte, while in angiosperms it is a product of fertilization, and is called the endosperm. The emergence of these nourishing tissues is of profound evolutionary value, and they are also food staples for most of the world’s population. Here, using Orthofinder to infer orthologue genes among novel and previously published datasets, we provide a comparative transcriptomic analysis of seed nourishing tissues from representative species of all main angiosperm clades, including those of early diverging basal angiosperms, and a gymnosperm representative. Our results show that, although the structure and composition of seed nourishing tissues has seen significant divergence along evolution, there are signatures that are conserved throughout the phylogeny. Conversely, we identified processes that are specific to species within the clades studied, and thus illustrate their functional divergence. With this, we aimed to provide a foundation for future studies on the evolutionary history of seed nourishing structures, as well as a resource for gene discovery in new functional studies.

**Significance Statement:** Within seeds a specialized structure is responsible for nourishing the embryo during its development. These nourishing tissues are also important sources of staple foods and feed. Here, we provide novel gene expression datasets of nourishing tissues of early diverging angiosperms, and use this information for a meta-analysis to identify pathways conserved, or divergent, throughout evolution. Thus, we aim to provide a resource for gene discovery for seed biology studies.

## Introduction

The seed nourishing tissues are specialized structures that provide nutrients and support for the developing embryo. There is an outstanding diversity of plant seeds and strategies for nutrient transfer and storage (Baroux *et al*., 2002; Linkies *et al*., 2010; Chen *et al*., 2017a). In most angiosperms, these functions are carried out by the endosperm, the development of which, like that of the embryo, is derived from a fertilization event (Nawaschin, 1898; Guignard, 1899; Friedman, 1998). However, this coupling to fertilization is an innovation of the angiosperms: gymnosperm seeds do not produce an endosperm, but their embryos are surrounded by a nutritive tissue that results from the proliferation of the megagametophyte (Singh & Johri, 1972; Hardev, 1978; Norstog, 1982; Haig & Westoby, 1989; Haig, 1992; Linkies *et al*., 2010; Sakai, 2013; Rodrigues *et al*., 2018).

These seed nourishing structures can be persistent or be consumed by the developing embryo. Seedlings of species with persistent endosperms, like cereals, rely on it for nourishment (Liu *et al*., 2022). Morphologically, those cereal endosperms are often larger and more prominent than those of other angiosperms, occupying a larger proportion of the mature seed (Sreenivasulu *et al*., 2010; Sreenivasulu & Wobus, 2013). Because the endosperm is typically the primary source of nutrients for the developing embryo, the monocot cotyledon is often very small or absent (Sabelli & Larkins, 2009). Contrastingly, in most eudicots, although serving a critical role during early seed development, the endosperm is much smaller, and is consumed as seeds mature (Le *et al*., 2010; Sreenivasulu & Wobus, 2013). This leaves the cotyledons as the main storage tissue for the germinating seedling (Weber *et al*., 2005). Like in most eudicots, in early divergent angiosperms such as in the Amborellaceae, Nymphaeales and Austrobaileyales, the endosperm is not involved in nutrient storage but rather its main role is transfer (Floyd & Friedman, 2000; Friedman & Bachelier, 2013; Povilus & Friedman, 2022). In fact, in *Nymphaea*, the storage function is handled by the perisperm, which derives from the ovule nucellus (Povilus *et al*., 2015). This is also the case in other species with perispermic seed habits, like those of the Amaranthaceae and Malpighiaceae families (Burrieza *et al*., 2014; Souto & Oliveira, 2014; Vandelook *et al*., 2021).

The transcriptional landscape of seed nourishing tissues is expected to vary between various plant clades and to reflect their diverse evolutionary histories and functional adaptations. In fact, seed nourishing tissues have substantially diverged in morphology and developmental patterns throughout the more than 300 million years since angiosperms diverged from their common gymnosperm ancestor (Zimmer *et al*., 2007; Ran *et al*., 2018; Lubna *et al*., 2021). For instance, endosperms of different species can have varying ploidies and maternal to paternal genome ratios, depending on the type of megagametogenesis that occurs in each given species, and on the type of reproduction, sexual or asexual (Baroux *et al*., 2002; Geeta, 2003; Rangan, 2020). Moreover, endosperms can follow different modes of development (Floyd & Friedman, 2000): nuclear endosperms, like those of *Arabidopsis thaliana* and of cereals, are coenocytic in the first days of development, and only cellularize at a later time point (Olsen, 2004); while in cellular endosperms, like those of the Solanaceae, Lamiales and of early diverging angiosperms, karyokinesis is always coupled to cytokinesis (Povilus *et al*., 2015; Oneal *et al*., 2016; Roth *et al*., 2018); finally, in the more uncommon helobial endosperms, two chambers are formed, each of which undergoes a different developmental program (Swamy & Parameswaran, 1963). Even given this vast diversity of nourishing tissues, we hypothesized that there should be common transcriptomic signatures that remained unchanged throughout evolution.

Evolutionary convergence and divergence at the transcriptional level refer to the similarities and differences in gene expression patterns between different species (Ran *et al*., 2018; García de la Torre *et al*., 2021). Recently, evolutionary transcriptomics has greatly benefited from advances in orthologous inference pipelines such as OrthoFinder (Emms & Kelly, 2019). One major advantage is to accurately identify orthologous genes across large numbers of species, allowing for a more comprehensive comparative gene expression analysis, and thus the potential to uncover kingdom-wide mechanisms (Julca *et al*., 2021, 2023). Such studies have led to substantial insights into the evolution of gene regulation and the role of gene expression in phenotypic divergence between species (Ferrari *et al*., 2019; Yu *et al*., 2020; Gao *et al*., 2021; García de la Torre *et al*., 2021). The availability of large-scale transcriptomic datasets, combined with advances in computational and analytical tools, has enabled researchers to identify co-expression networks and functional modules across different species (Hansen *et al*., 2014; Qiao *et al*., 2016; Vercruysse *et al*., 2020). The integration of the large amount of publicly available datasets with novel analytical methods is opening the way for the community to generate and test powerful hypotheses about gene function and evolution (Leebens-Mack *et al*., 2019; Julca *et al*., 2023).

In this study we leverage the diversity of seed-nourishing tissues and perform a multispecies comparative transcriptomic study. Our main question is what the transcriptional signatures are, that are conserved in different nourishing tissues across the plant phylogeny. Additionally, we aimed to identify unique transcriptional features of the nourishing tissues of species in divergent plant clades. We chose to examine distinct clades with different endosperm conformations to maximize the power of our inferences. Specifically, we used early divergent angiosperms represented by *Amborella trichopoda* (Amborella) and *Nymphaea caerulea* (water lily). We included maize (*Zea mays*) and rice (*Oryza sativa*) which possess a large and persistent starchy endosperm (Zheng & Wang, 2015). As core eudicots we included the well-studied endosperm of *Arabidopsis thaliana* (Arabidopsis), and those of *Solanum peruvianum* (wild tomato) and *Mimulus guttatus* (monkeyflower). This selection also covers species with distinct patterns of endosperm development: nuclear, for Arabidopsis, rice and maize (Olsen, 2004); and *ab initio* cellular, for the remaining species (Tobe *et al*., 2000; Povilus *et al*., 2015; Oneal *et al*., 2016; Roth *et al*., 2018). As representatives of the gymnosperms, we studied the megagametophyte transcriptome of *Pinus pinaster* (maritime pine) and *Ginkgo biloba* (ginkgo), although only the former was used for the clade-specific comparisons (Rodrigues *et al*., 2018; Zumajo-Cardona *et al*., 2021). Our analyses provide a comparative framework to identify potential orthologs of genes of interest that are expressed in the seed nourishing structures of land plants. With this, we aim to provide a resource for future functional studies in seed biology.

## Experimental Procedures

Extended materials and methods are provided as Supporting Information (Methods S1). We sampled transcriptomes of the earliest available stages of nourishing tissues in species of all main angiosperm clades. We generated transcriptomes of laser microdissected endosperms and of leaves of *Nymphaea caerulea* and *Amborella trichopoda* (Figure S1), and of manually dissected megagametophytes of *Pinus pinaster*. We obtained transcriptomes of nourishing tissues and leaves from public repositories for *Gingko biloba*, *Arabidopsis thaliana*, *Solanum peruvianum*, *Mimulus guttatus, Oryza sativa* and *Zea mays*. Details of data provenance and availability are in Table S1. We inferred Orthogroups for all species using Orthofinder (Emms & Kelly, 2019). Mapping to reference genomes was performed with Hisat2 (Kim *et al*., 2015), and the genome versions are described in Table S1. We performed differential gene expression analyses on each species comparing nourishing tissues vs. leaves using DESEQ (Love *et al*., 2014). DEGs per species were translated to OG assignments, resulting in differentially expressed orthogroups (DEOGs). Using DEOGs we performed set operations to identify conserved and divergent OG sets among the species studied. Enrichment analyses and protein network inferences were performed in STRING (Szklarczyk *et al*., 2017) using the best Arabidopsis ortholog Blast hit for the OGs. All additional data plotting and statistical tests were performed in R (version 4.2.0).

## Results

### Broad gene expression patterns discriminate the larger angiosperm clades: Monocots set the difference

To test the hypothesis that there should be some degree of conservation between seed nourishing tissues of different species, we compared the gene expression profiles of seed nourishing tissues to those of somatic ones (in this case leaves). To include representatives of all main angiosperm clades, we generated novel transcriptomes of endosperms of the early diverging angiosperms *Amborella trichopoda* and *Nymphaea caerulea*. We also hypothesized that some of those transcriptomic signatures could already be present in the gymnosperm megagametophyte, which fulfills the nourishing function in seeds of those species. Therefore, together with analyzing previously published transcriptomes of ginkgo megagametophytes (Zumajo-Cardona et al., 2021), we also generated novel transcriptomes of megagametophytes of *Pinus pinaster* at early stages of seed development. We also included in the analysis the perisperm of *Nymphaea*, because in this species it is the perisperm, and not the endosperm, that functions as a storage tissue. For the remaining species we used publicly available datasets (described in Methods S1). For all species we used, or produced, datasets from nourishing tissues at the earliest available stages of seed development (Figure 1 and S1, Table S1).

**Figure 1.**
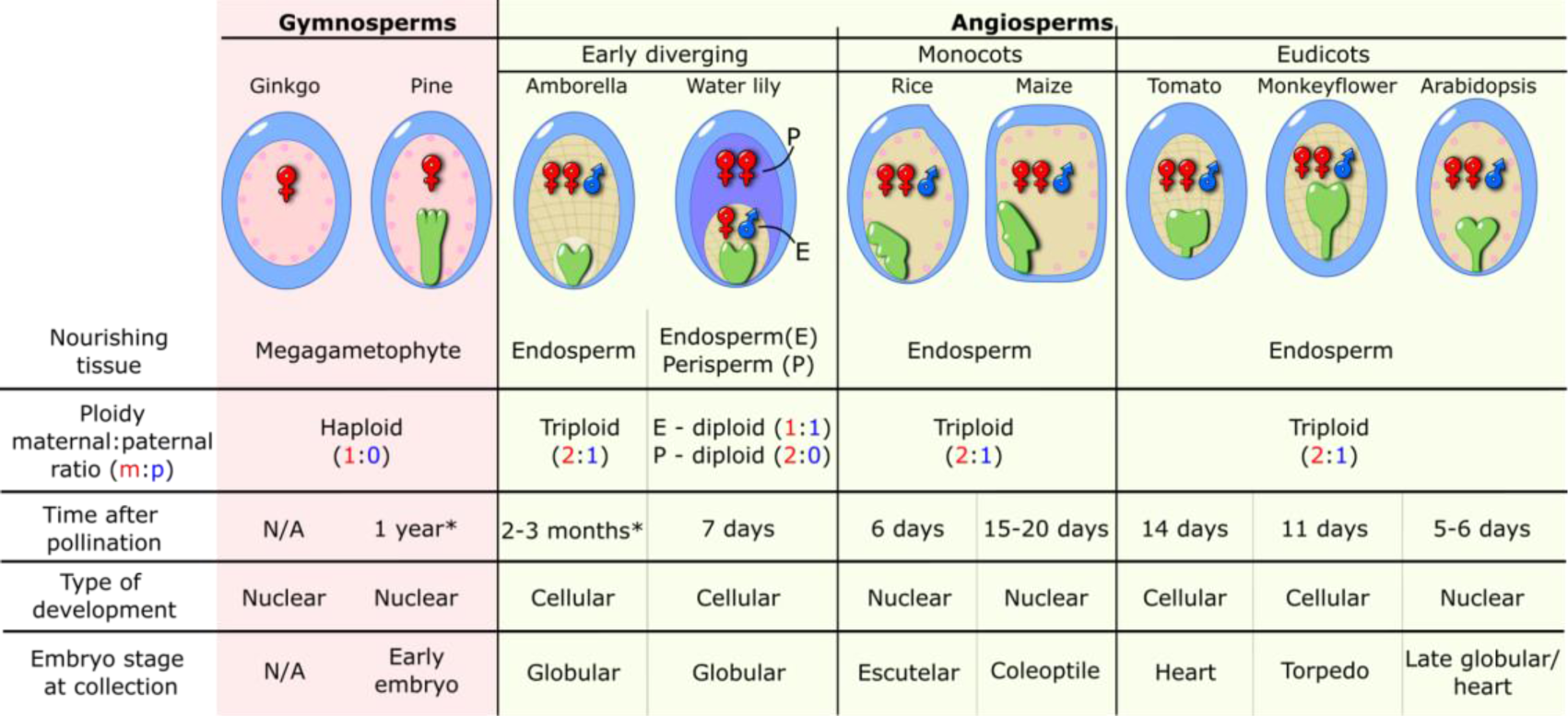
Seed structure and sample information. The diagrams at the top indicate the seed structure for all species studied. The type of structure, ploidy, chronological age, type of development and corresponding embryo stages are indicated in each row. Male and female symbols indicate the ploidy and parental contribution to each nourishing structure. The sporophytic tissues are indicated in blue: integument, seed coat and/or pericarp. The water lily perisperm (P), is indicated in purple. Embryos are indicated in green. The Ginkgo samples were collected from mature ovules and not developing seeds, thus lacking an embryo. * indicates an estimate, based on the pollination season. Further information and references to previous studies are found in Table S1.

To compare the transcriptional landscapes among tissues of divergent clades, we used Orthofinder (Emms & Kelly, 2019), and assigned orthologous genes to orthogroups (OGs). These OGs represent the set of genes from the species used in this study, which descended from a single gene in the last common ancestor of this set of species (Emms & Kelly, 2019). Eighty nine percent of the input genes were assigned to 34,041 OGs. Rice had the largest percentage of unassigned genes (16.7%) and the largest number of duplications were detected on the monocot node clade in the phylogenetic tree produced (3,405, Figure 2A-B). The species tree of Figure 2B reflects the individual OGs gene trees and the gene duplication events that were identified using our datasets. On the other hand, *Ginkgo* and *Pinus* revealed the smallest OG overlap with the other species included in the study while also displaying numerous gene duplication events (Figure 2B-C). The largest OG overlap is between the two monocot species analyzed, sharing 13,291 OGs, followed by overlaps within the eudicot species (Figure 2C). Rice and maize displayed striking similarities and overlapping OG sets (total and differentially expressed), despite maize having almost twice the number of genes annotated and assigned to OGs, compared to rice (Figure 2A). Details of the OGs identified and the species gene correspondence can be found in Table S2.

**Figure 2.**
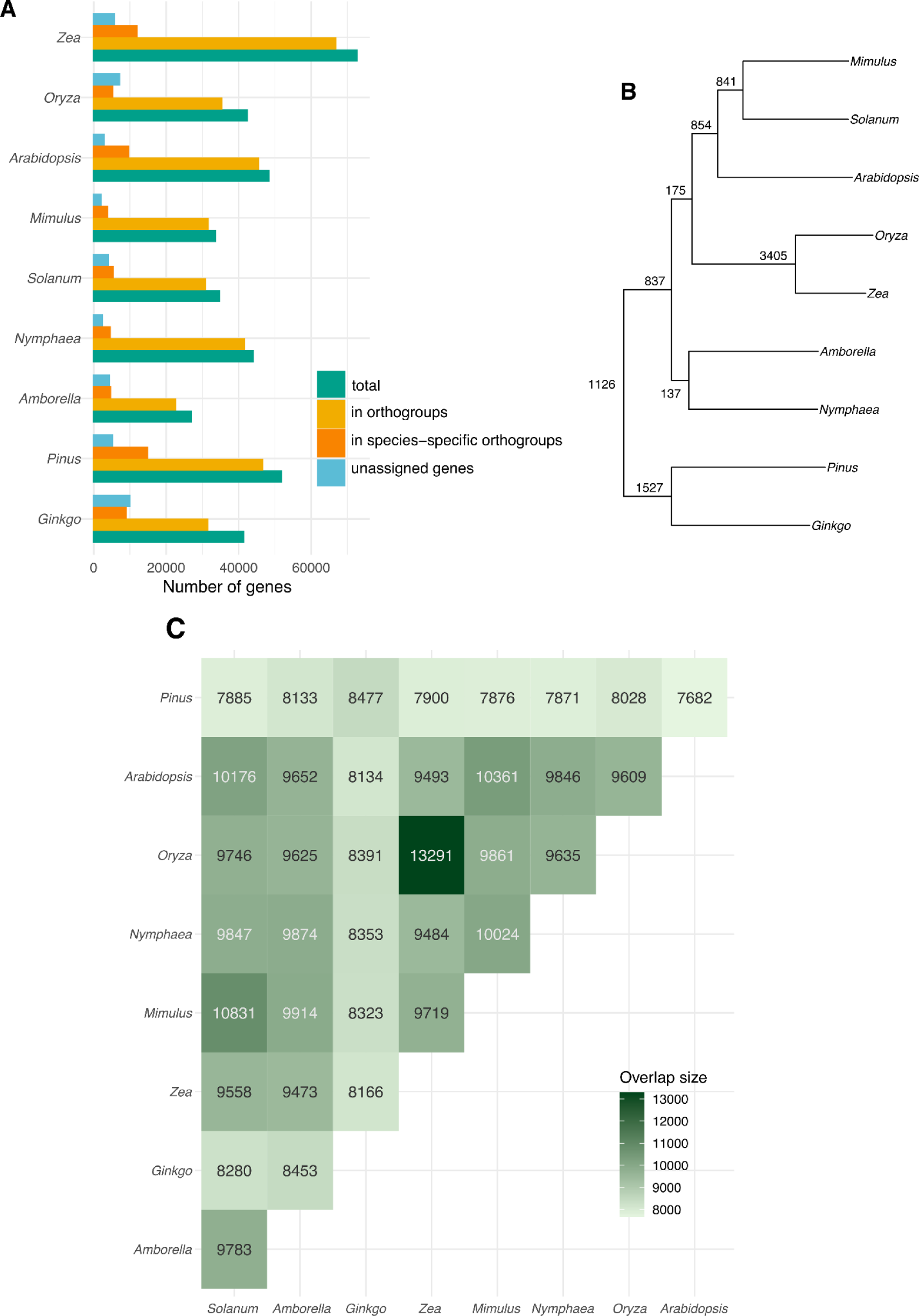
Orthology inference pipeline descriptive results. A. Overall statistics per species. B. Species trees with nodes displaying gene duplication events C. Overlap in the number of OGs in each pair of species.

Having assigned genes into OGs, we used expression estimates for leaves and nourishing tissues and ran a PCA analysis to assess the variation among our datasets. Most of the variation in overall gene expression is captured by PC1, accounting for 34.83% of the total variance and exhibiting a significant correlation (0.75) with the corresponding clade (Figure 3A and Figure S2): the transcriptomic diversity of a tissue is determined by whether it belongs to a gymnosperm, early divergent angiosperm, eudicot, or monocot. Monocots are prominently located on the left side of the PCA (Figure 3A) and are distinct from other angiosperms, including from the more divergent plant clades. This pattern may be a reflection of the specialization of the large, persistent and starchy endosperm of cereals (Poaceae) (Zheng & Wang, 2015), which are the only representatives of the monocot clade in this study. Interestingly, however, this divergence also extends to the somatic tissues. The PC6, which accounts for 3.49% of the variance, exhibits the second strongest and significant correlation (0.71) with the tissue variable (Figure S2). The top loadings of PC6 therefore define the transcriptional network of angiosperm nourishing tissues, and correspond to OGs annotated with functions related to nutrient storage: CRUCIFERIN 2 (CRU2) and RMLC-LIKE CUPIN (Pang *et al*., 1988), CYSTEINE PEROXIREDOXIN 1 (PER1) (Haslekås *et al*., 1998), and VICILIN-LIKE SEED STORAGE PROTEIN (PAP85)(Shutov *et al*., 1995) (Figure S2 and Table S3).

**Figure 3.**
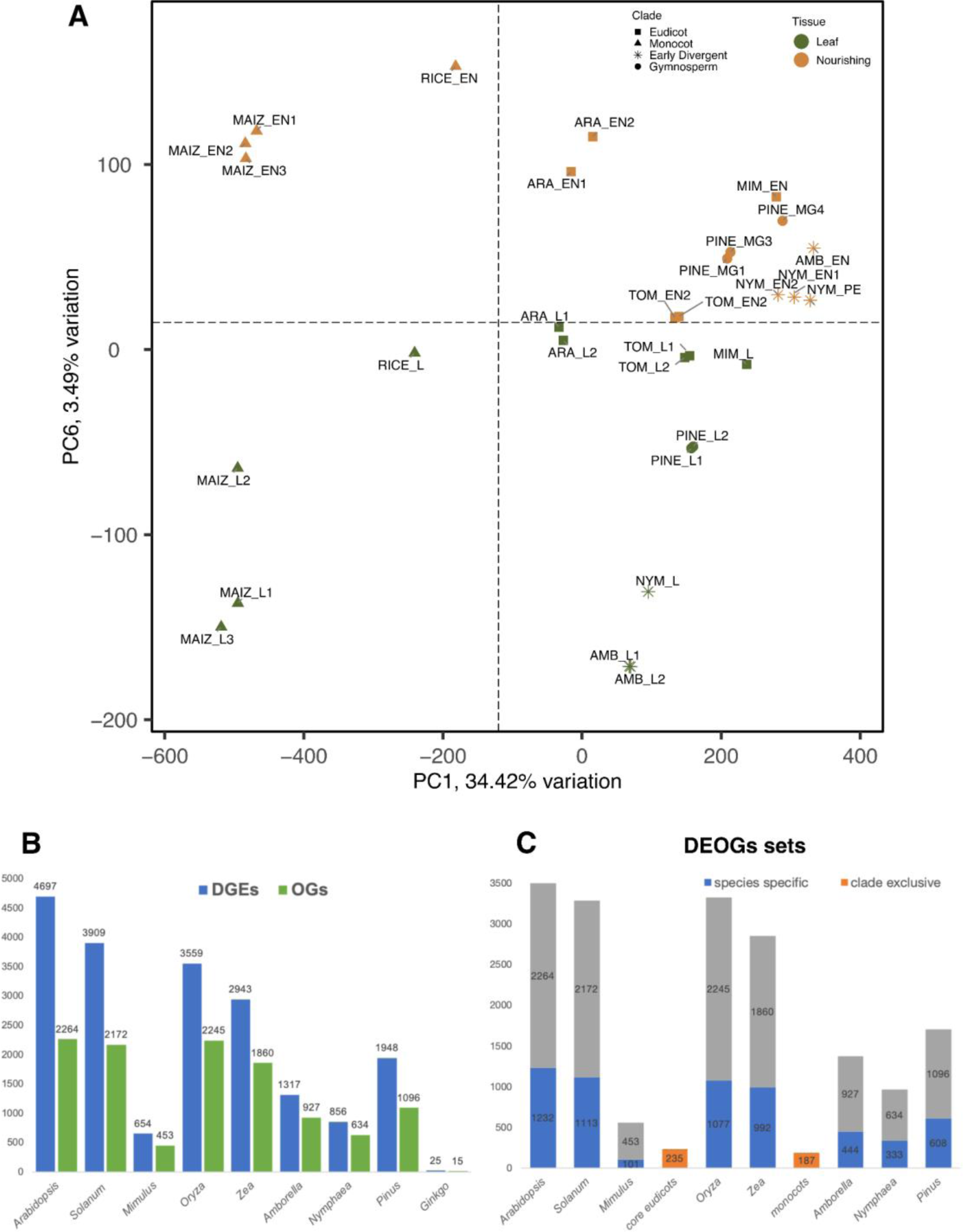
A. PCA describes broad gene expression patterns of photosynthetic vs. nourishing tissues in a broad range of land plants. B. Differentially expressed genes (DEGs, in blue) and corresponding differentially expressed orthogroups (DEOG in green) per species. C. DEOGs set description. Fractions of species specific OGs are depicted in blue, shared OGs in grey. mim=*Mimulus*, nym=*Nymphaea*, amb =*Amborella*, gin=*Ginkgo*, pin=*Pine*, maiz=*Zea*, tom=*Solanum*, rice=*Oryza*, ara=*Arabidopsis*, L= leaf, MG=megagametophyte, EN=endosperm, PE=perisperm.

Interestingly, the *Pinus* nourishing tissue clusters together with those of the angiosperms, regardless of their intrinsic biological differences, such as ploidy and parental origin (Figure 1) (Linkies *et al*., 2010). However, for the other gymnosperm representative, Ginkgo, we only detected 25 DEGs in its nourishing tissue, which was orders of magnitude lower than for all other species (Figure 3B). This may be a consequence of the stage from which the transcriptomes were obtained: the Ginkgo megagametophytes originated from developing ovules, and not fertilized seeds (Stage S9 in (Zumajo-Cardona *et al*., 2021)). Therefore, although we included the Ginkgo datasets for calling OGs, we did not include this species in the follow-up analyses. Although all eudicot nourishing tissues cluster together in the upper right quadrant of the PCA, there appears to be a close relationship in the overall patterns of gene expression of nourishing structures of the early diverging angiosperms, *Amborella* and *Nymphaea*. This is in contrast to the more divergent samples of *Solanum*, *Arabidopsis*, and *Mimulus*, which probably reflects specialization of the nourishing tissues in the later diversified clades of angiosperms.

### Cross-species DGE analyses using OGs

To find transcriptional signatures that are characteristic of seed nourishing tissues, for each species we performed an analysis of differential gene expression (DGE) between the transcriptomes of the leaves and of the nourishing tissues (Table S3). To enable cross-species comparisons, an OG identity was assigned to each DEG, delimiting Differentially Expressed OrthoGroups (DEOGs; Table S3). Some OGs consisted of multiple genes and some genes did not have OG assignment (Figure 2), resulting in less DEOGs than actual DEGs (Figure 3B). It is important to note that the number of identified DEOGs can be determined not exclusively by the biological properties of the taxa and tissue but also by technical limitations, such as genome annotation or library complexity, resulting in low numbers of identifiable DEOGs in the tissue. A significant proportion of the DEOGs were species-specific, meaning they did not overlap between species (Figure 3C, Figure S2). DEOGs in the endosperms of rice and maize show the highest degree of overlap. This can be attributed to their similar transcriptional strategies, which also explains why they are grouped together on PC1 (Figure 3A). The second largest set of overlapping DEOGs is found in *Solanum* and *Arabidopsis* (Figure 3C, Figure S2), which also entail the best annotated eudicot genomes. In contrast, overlaps of *Mimulus* DEOGs with the other eudicot taxa were circa one order of magnitude lower (Figure S2). Due to this low number of shared features, the DEOGs of *Mimulus* were not included in the overlapping eudicot set (described below). Among all species tested, *Arabidopsis*, *Solanum*, *Oryza*, and *Zea* had the highest number of DEOGs in their endosperms (Figure 3B-C, Figure S2, Table S1, Table S3). We therefore focused our eudicot and monocot clade analyses on the shared sets of these taxa. Next, we investigate the identity of shared, non-shared, and species-specific (or unique) DEOGs between and within the main plant clades to describe the transcriptional networks of their nourishing tissues. For the sake of simplicity, the full transcriptional networks are provided as Supplementary Figures, and in each Figure we focus on a pair of relevant DEOG networks per comparative analysis.

### Common transcriptional features of seed nourishing tissues across all tested angiosperms

First, we identified 175 OGs that were enriched in angiosperm seed nourishing tissues in at least one monocot, one dicot, and one early diverging taxon (Hypergeometric test p-value: 2,70E-02). We constructed functional clusters out of these OGs, to assess what biological functions are specifically enriched in seed nourishing tissues. Sixteen functional clusters were identified, two of which were prominent and we describe here in more detail (Figure S3, Table S4). First, one cluster involved in cell cycle and nucleosome architecture drives the enrichment of the KEGG term “Nucleosome Core’’ (KW-0544; Figure S3, Table S4). At the core of this cluster are five histone-annotated OGs (Figure 4, Figure S3 and Table S3), which are functionally linked to two OGs representing subunits of the RNA Polymerase IV (RNAP IV). Also interacting with the main histone cluster are three OGs that represent A2 and type B cyclins, as well as UV-B-INSENSITIVE 4 (UVI4), a regulator of cyclin turnover (Heyman *et al*., 2011). Additionally, there are four OGs representing components of the microtubule machinery. These OGs and the species-specific genes they encompass represent a gene network that is involved in cell cycle transitions and cell division in the angiosperm nourishing tissues. Interestingly, an OG annotated as *SYP111*/*KNOLLE* is also enriched. SYP111/KNOLLE is necessary for endosperm cellularization in Arabidopsis (Tiwari *et al*., 2010; Park *et al*., 2018), and our results thus indicate that its role in nourishing tissues is likely evolutionarily conserved.

**Figure 4.**
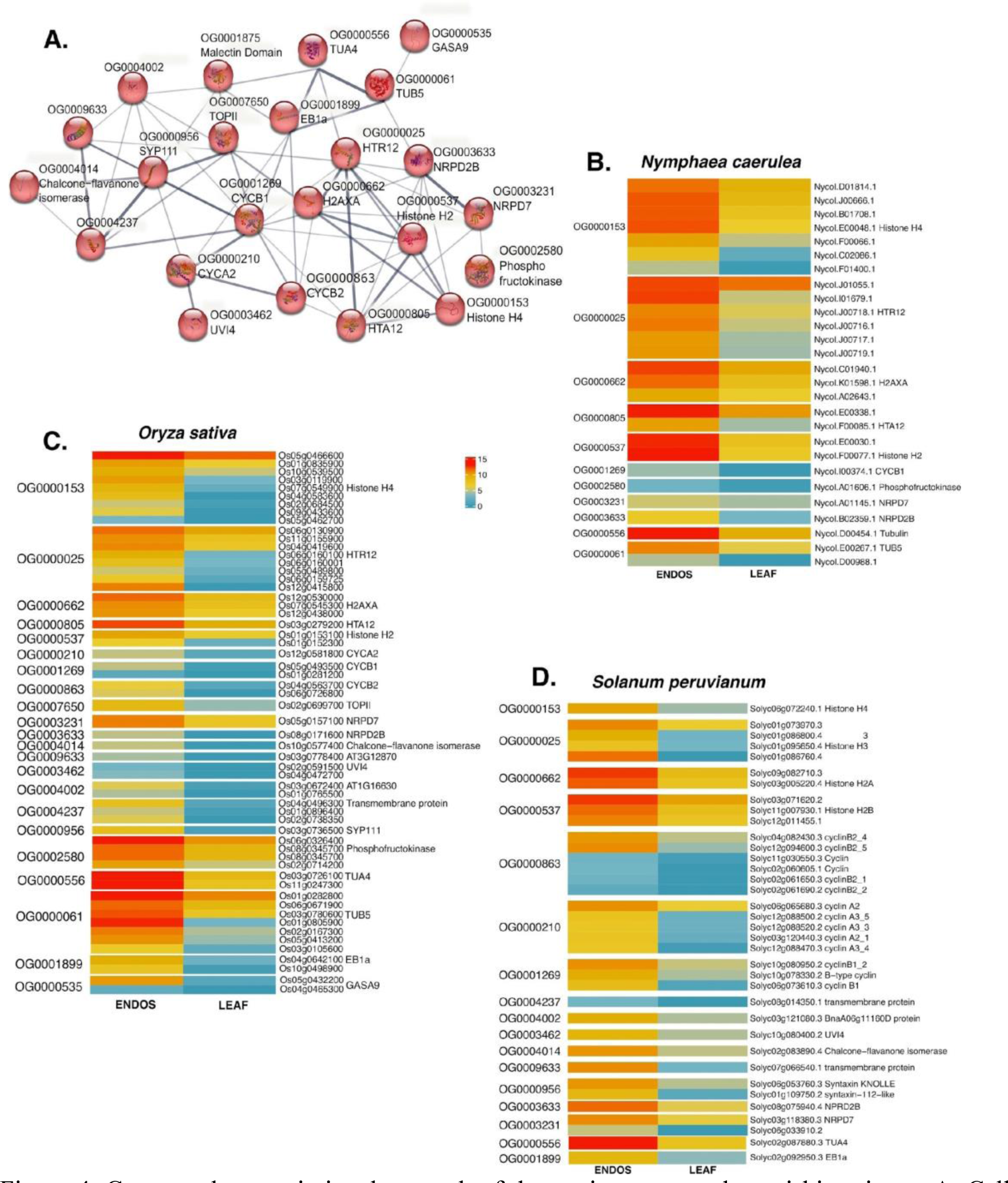
Conserved transcriptional network of the angiosperm seed nourishing tissue. A. Cell cycle and nucleosome architecture interacting network. B-D. Orthogroup to DEG correspondence and expression values in CPM for B. *Nymphaea caerulea.* C. *Oryza sativa* and D. *Solanum peruvianum*.

To illustrate the structure of OGs in each of our target species, we created heatmaps displaying the expression values for all DEGs belonging to the OGs in the cluster (Figure 4 and Figure S4). OGs annotated as histones and cyclins, in particular, are composed of multiple genes per species. This pattern indicates their diversification in the endosperm. Other OGs in this cluster do not include a significant number of DEGs. For instance, those representing RNAP IV subunits or components of the microtubule assembly machinery typically have only one DEG in most species examined (Figure 4 and Figure S4). Importantly, histone variants are emerging as modulators of chromatin architecture (Borg *et al*., 2021; Bhagyshree *et al*., 2023). The histone proteins that we identified in this conserved cluster of enriched OGs are therefore strong candidates for determining the specific chromatin state of the endosperm.

A second important cluster conserved in all clades relates to nutrient storage (Figure 5A). The term “nutrient reservoir activity, and lipid droplet” was found enriched in the conserved sets of 175 OGs (STRING cluster CL:39149). This cluster comprises the OGs that drive most of the variation of the PC6 (Figure S2). These OGs are mostly annotated as CUPINS, which are non-enzymatic proteins involved in seed storage (Dunwell, 1998). One example being CRU2 (Pang *et al*., 1988; Lin *et al*., 2013). To display the relative expression of these genes in the endosperm we plotted heatmaps of expression of DGEs belonging to these OGs in all species studied (Figure 5B-D, Figure S5). Although most OGs in this cluster are single gene OGs, OG0000230, annotated as *CRU2*, shows multiple members in all species, evidencing diversification of cruciferin storage proteins in angiosperms. Indeed, the CUPIN superfamily is often described as an hallmark example of functional diversity (Dunwell, 1998; Dunwell *et al*., 2001, 2004). Interestingly, a well-defined set of glutelins also belongs to this OG, which is expected as glutelins are the main storage proteins in rice grains (Mitsuda *et al*., 1967). The OG0007276 is best annotated as a Late Embryogenesis Abundant protein (LEA), *LEA4-5* (Battaglia *et al*., 2008), and is similarly present in the nutrient transfer cluster (Figure 5). Furthermore, three more OGs annotated as LEA proteins are present in the larger set of 175 OGs conserved expressed in endosperms (Table S4).

**Figure 5.**
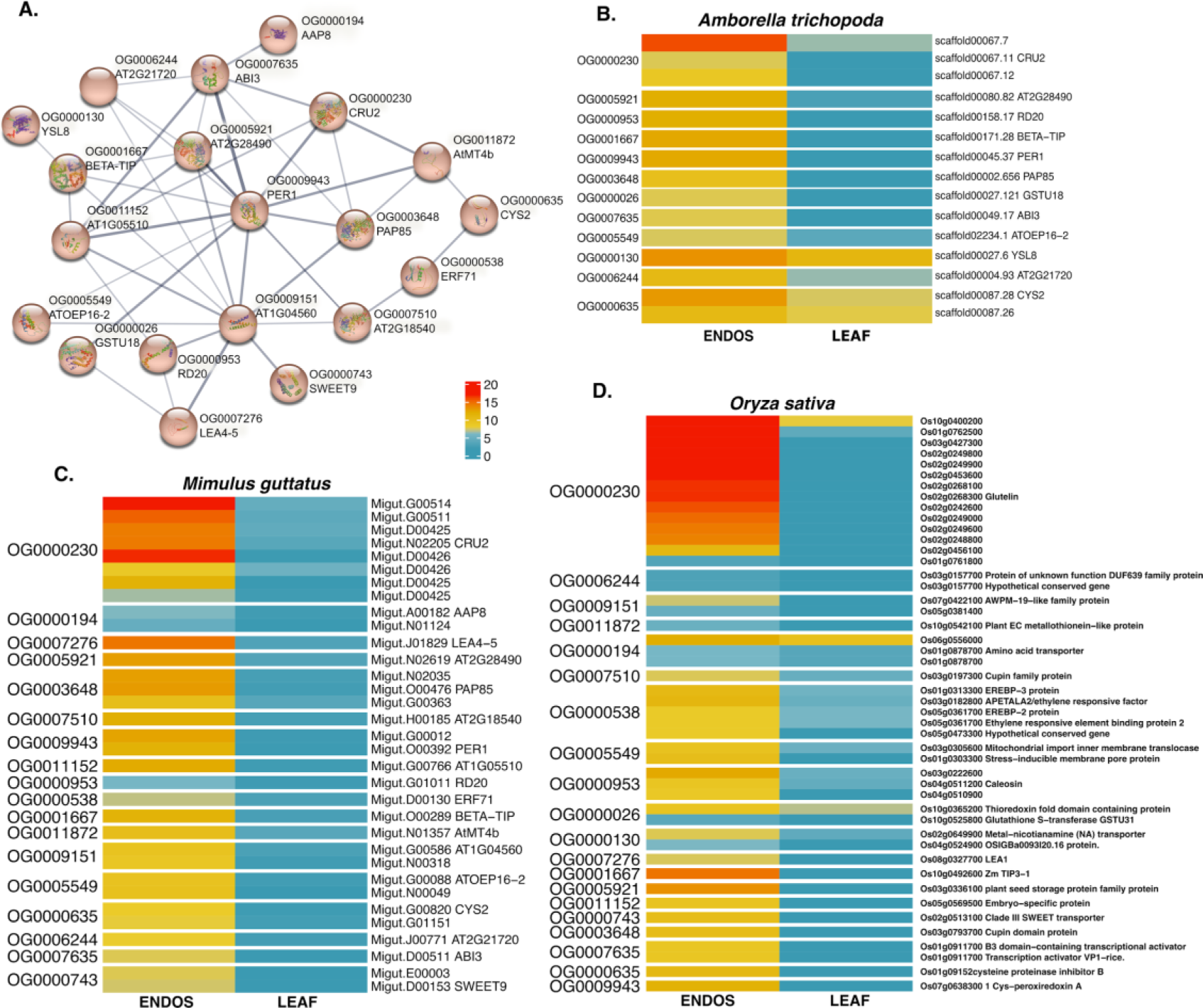
Conserved transcriptional network of the angiosperm seed nourishing tissue. A. Nourishing cluster protein interacting network. Orthogroup to DEG correspondence and expression values in CPM for B. *Amborella trichopoda*. C. *Mimulus guttatus* and D. *Oryza sativa*.

Interestingly, OGs related to Abscisic acid (ABA) signaling were also found on this interacting cluster. Specifically, *ABSCISIC ACID INSENSITIVE 3* (*ABI3*), encoding a transcription factor that participates in ABA-regulated gene expression during seed development (Mönke et al., 2012), and regulates seed storage genes (Lara *et al*., 2003). The same was true for *GLUTATHIONE S-TRANSFERASE 18* (*GSTU18*), as glutathione metabolism plays roles in seed dormancy and germination (Koramutla *et al*., 2021). Other OGs of relevance in this cluster encode seed transporter proteins, such as AMINO ACID PERMEASE 8 (AAP8) and the sugar transporter SWEET9 (Table S4) (Schmidt *et al*., 2007; Lin *et al*., 2014).

The shared transcriptional network of angiosperm nourishing tissues also includes four OGs that form a chaperone/heat shock cluster (Table S4, Figure S3). Moreover, there are a number of OGs encoding transcription factors (TFs) that are shared among angiosperms. These include the MYB-type TFs MYB65 and MYB100 (Stracke *et al*., 2001; Millar & Gubler, 2005). Interestingly, only one OG annotated as encoding a MADS-box transcription factor was present in the angiosperm common set, which was composed of a large set of SEPALLATA-like proteins across the angiosperms (Zahn *et al*., 2005).

Likewise present was *RGE1*, also known as *ZHOUPI*, which has known roles in endosperm development (Kondou *et al*., 2008; Yang *et al*., 2008). Moreover, two OGs annotated as encoding subtilisin proteases (SBT1.1, SBT1.7) are consistently present in angiosperm endosperms, which fits with their functions during seed formation (Rautengarten *et al*., 2008; D’Erfurth *et al*., 2012). These subtilases may also fulfill functions similar to that of ALE1, which together with ZHOUPI acts in an intercompartment signaling pathway between endosperms and embryos (Xing *et al*., 2013; Moussu *et al*., 2017). Other conserved OGs in the angiosperm nourishing tissues but not forming clusters in the STRING protein network encode proteins involved in hormone biosynthesis and signaling. These proteins include YUCCA11, involved in auxin biosynthesis during early seed development (Cheng *et al*., 2007; Figueiredo & Köhler, 2018), and the brassinosteroid response regulator BRASSINOSTEROID RELATED HOMEOBOX 2 (Hasegawa *et al*., 2022) (Table S4).

### The transcriptional network of the pine megagametophyte

Although also bearing seeds, the gymnosperms are expected to exhibit some degree of evolutionary divergence when it comes to the seed nourishing tissues. Their seeds contain a nourishing tissue resulting from the proliferation of the haploid female megagametophyte, which is not the product of fertilization (Hardev, 1978; Norstog, 1982). We aimed to uncover similarities and differences of the nourishing tissues of angiosperms and gymnosperms, using *Pinus pinaster* as a reference gymnosperm species. For this, we intersected the set of exclusive DEOGs of the conserved angiosperm nourishing tissue (see above and in Table S4) and compared it to the full set of DEOGs of the pine megagametophyte. Sixty-four OGs that were conserved in angiosperms were also found as DEOGs in pine (Table S5). We found that members of the two main groups of conserved OGs in the angiosperm private network were partly present in the pine DEOG set. This included OGs annotated as histones, RNAP IV subunits and microtubule machinery (Table S5). The only OG encoding a MADS box TF that was found conserved in all angiosperms, annotated as *SEP3* (Immink *et al*., 2009), was not present in the DEOGs uncovered in pine. This was expected, given that *SEP* genes are specific to the angiosperms (Zahn *et al*., 2005). Other sets of DEOGs conserved in angiosperms and absent in pine include those annotated as encoding gibberellin regulated proteins, GASAs (Herzog *et al*., 1995), and the subtilisin-like proteases SBT1.1 and SBT1.7 (Rautengarten *et al*., 2008; D’Erfurth *et al*., 2012).

Next, we identified 608 OGs that are unique to the nourishing tissue of pine (Hypergeometric test p-value: 1,49E-21, Figure 3C, Figure S2; Figure S6). Enrichment of several terms shows the specialization of pine megagametophyte in the biosynthesis of specific secondary metabolites (ATH01110 KEGG term “Biosynthesis of secondary metabolites’’). Moreover, we found unique pine OGs involved in pathways related to jasmonic acid (JA), alpha linoleic acid, ascorbic acid, and phenylpropanoid and flavonoid biosynthesis. Regarding JA, OGs related to both its biosynthesis and signaling were enriched (Fig. 5A). For example, this includes OGs annotated as encoding LIPOXYGENASE 3 (LOX3), which is involved in JA biosynthesis (Vick & Zimmerman, 1983), as well as the JA effector JASMONATE-ASSOCIATED 10 (JAZ10) (Thines *et al*., 2007). Importantly, JA has been shown to be involved in the regulation of seed size and germination (Linkies & Leubner-Metzger, 2012; Pan *et al*., 2020a; Hu *et al*., 2021). Moreover, we identified a cluster involved in phenylpropanoid biosynthesis (Fig. 5B), a process tightly linked to lignin biosynthesis (Douglas, 1996). Although lignin has been found to play a role in angiosperm seed coats (Tobimatsu *et al*., 2013; O’Leary, 2020), the private occurrence of this cluster in the pine megagametophyte may be related to the exposed nature of the gymnosperm seed, and this particular cluster of genes may be involved in mediating its protection (Emonet & Hay, 2022). Flavonoids are also a product of the phenylpropanoid pathway (Vogt, 2010), and we identified a cluster involved in their biosynthesis (Figure 6B). Finally, still regarding metabolite biosynthesis, the term “L-ascorbic acid biosynthesis process” was enriched in our analyses. Key OGs in this pathway include those annotated as *GLUTATHIONE S-TRANSFERASE 20* (*GSTU20*) and *DEHYDROASCORBATE REDUCTASE 2* (*DHAR2*; Figure 6B) (Wagner *et al*., 2002; Zhang *et al*., 2015). In addition to their roles in ascorbic acid biosynthesis, these OGs may be involved in gymnosperm seed dormancy and germination, as part of the glutathione metabolism machinery (Koramutla *et al*., 2021).

**Figure 6.**
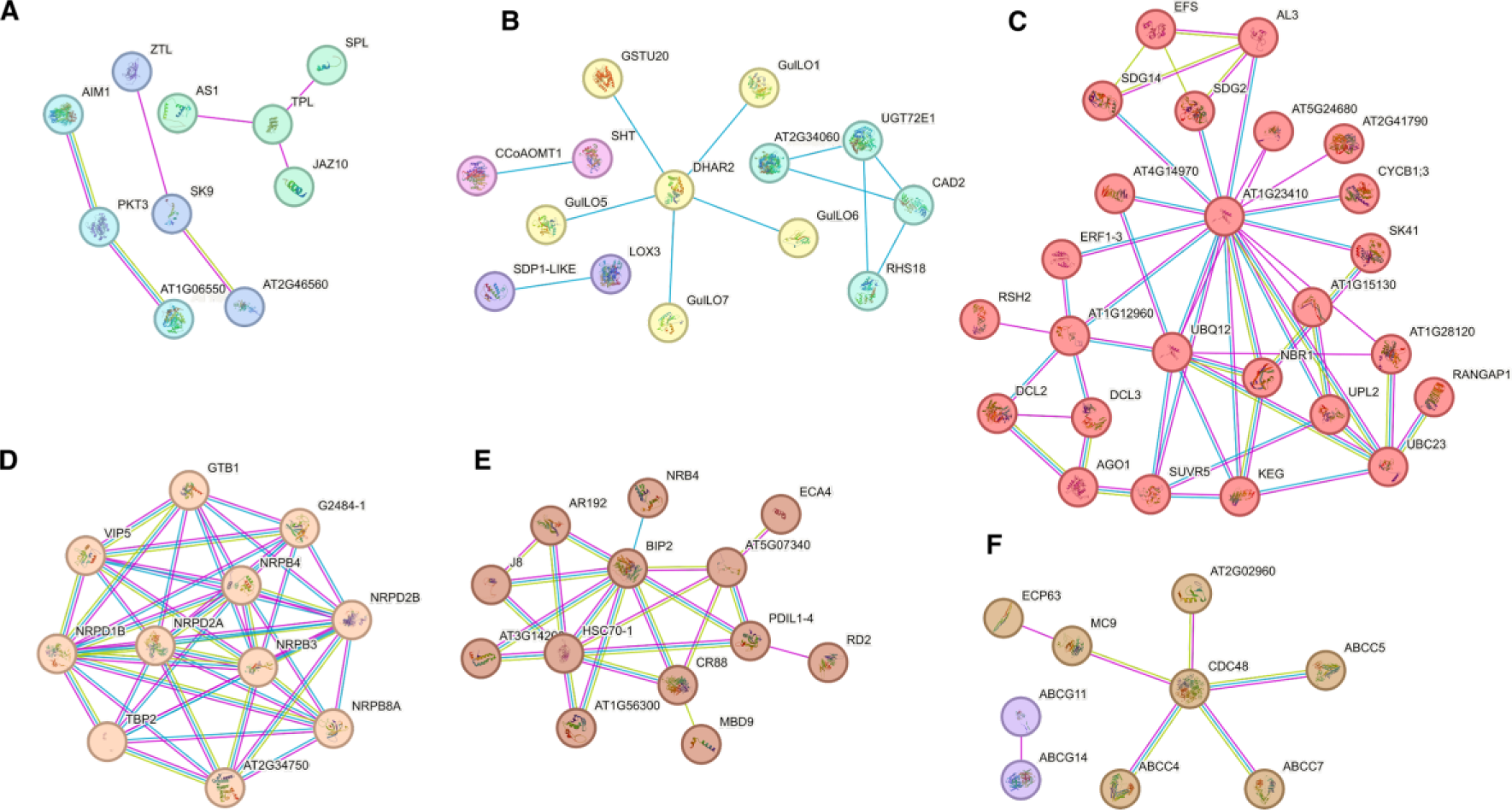
Orthogroup-based putative protein clusters unique to the pine megagametophyte transcriptional network. A. Jasmonate related. B. Metabolite biosynthesis C. Epigenetic machinery D. RNA polymerases. E. Chaperones F. ABC-type transporters.

The pine nourishing tissues also entail a specific set of epigenetic machinery, as illustrated by the largest cluster identified in our analyses (Figure 6C). This includes, for example, two OGs annotated as encoding DICER-LIKE (DCL) proteins, and as ARGONAUTE 1 (AGO1) (Henderson *et al*., 2006; Fang & Qi, 2016). Moreover, this large interacting cluster of proteins includes four SET domain-containing OGs annotated as histone methyltransferases, namely, SET DOMAIN GROUP 2 (SDG2) (Guo *et al*., 2010), EARLY FLOWERING IN SHORT DAYS (EFS) (Kim *et al*., 2005), TRITHORAX 3 (SDG14) (Chen *et al*., 2017b), and SU(VAR)3-9-RELATED PROTEIN 5 (SUVR5) (Caro *et al*., 2012) (Figure 6C). This cluster also contains several OGs annotated as encoding proteins belonging to the ubiquitin conjugation pathway, such as UBIQUITIN 12 (UBQ12) (Callis *et al*., 1995) and UBIQUITIN-CONJUGATING ENZYME 23 (UBC23) (Kraft *et al*., 2005), among others (Figure 6C). These, together with a smaller cluster containing OGs annotated as *SKP1-LIKE 9* (*SK9*) and *ZEITLUPE* (*ZTL*) (Somers *et al*., 2000; Zhao *et al*., 2003), represent the ubiquitination pathway in the pine megagametophyte. This pathway may be particularly important for the programmed cell death events that occur in *Pinus*, as in other conifers, whereby all the embryos resulting from cleavage polyembryony except the dominant one are eliminated (Filonova *et al*., 2002), in a process in which the megagametophyte may play a role (Williams, 2009).

Further related to epigenetic processes, we detected a separate large cluster in the protein interaction network which is composed of eight OGs encoding subunits of the RNAPs II, IV, and V (Figure 6D). The identification of OGs highlighting crucial components associated with 24-nt siRNA production and the silencing machinery, such as DCL3, AGO1 and RNAPs IV and V (Zhou & Law, 2015), along with previous findings showing the presence of these small RNAs in these tissues (Rodrigues *et al*., 2019), supports a role of the gymnosperm megagametophyte in RdDM. In angiosperms, 24-nt siRNAs in the endosperm were first proposed to originate solely maternally (Mosher *et al*., 2009), but are now thought to originate from both parental genomes in the endosperm, although with a strong maternal bias (Rodrigues *et al*., 2013; Xin *et al*., 2014; Martinez *et al*., 2018; Dziasek *et al*., 2023). Despite the differences between both nourishing tissue types, it appears that active silencing pathways associated with siRNAs are key, possibly to ensure genome stability and reproductive success.

One of the largest sets of interacting proteins uncovered in our analyses comprises eight OGs annotated as chaperones, as well as five OGs annotated as chaperones with DnaJ domains (Figure 6E). Interestingly, eight ATPase coupled ABC transporters are present in the OGs that are enriched in the pine megagametophyte (Figure 6F). They drive the enrichment of the GO term “ATPase-coupled transmembrane transporter activity”, and include OGs annotated as ABC transporters of Type 1 and Type 2, such as ABCG11 and ABCG14 (Bird *et al*., 2007; Zhang *et al*., 2014) (Figure 6F). Among other functions, ABC transporters have been implicated in the mobilization of surface lipids, accumulation of phytate in seeds, and in transporting several phytohormones, like cytokinin, auxin and abscisic acid (Kang *et al*., 2011; Zhang *et al*., 2014; Do *et al*., 2018).

### The transcriptional signatures of the nourishing tissues of the early divergent angiosperm lineages

The basal angiosperms investigated in this study are *Amborella* and *Nymphaea*. Although both are considered early divergent angiosperms, *Amborella* forms a sister clade to all remaining extant flowering plants (Amborella Genome Project, 2013; The Angiosperm Phylogeny Group, 2016). The endosperms of both species, however, exhibit striking differences: water lilies exhibit a diploid endosperm with a reduced size (Williams & Friedman, 2002), suggesting a more derived state compared to the larger triploid endosperm of *Amborella* (Tobe *et al*., 2000). We examined the private transcriptional networks of their nourishing tissues separately due to these distinctive characteristics.

For *Amborella*, we detected 437 OGs that were expressed in its nourishing tissue but not in any of the other angiosperms studied here (Hypergeometric test p-value: 3,62E-14, Figure 3C, Figure S2). The most prominent cluster in the private transcriptional network of the *Amborella* endosperm involves proteins related to cell cycle processes and chromosome organization (Figure 7A, Figure S7, Table S6). Particularly relevant OGs are three annotated as encoding cyclins, and two as components of RNAPs, namely, INCURVATA2 (ICU2) and REVERSIONLESS 1 (REV1) (Barrero *et al*., 2007; Takahashi *et al*., 2007). Additionally, there are OGs annotated as encoding proteins involved in chromosome organization, such as MINICHROMOSOME MAINTENANCE 5 (MCM5) (Shultz *et al*., 2009), MUTS HOMOLOG 7 (MSH7) (Culligan & Hays, 2000), and CURLY LEAF (CLF), a Polycomb Group protein (PcG) (Goodrich *et al*., 1997). Another set of OGs present in the protein interaction network are involved in transcriptional regulation and encode plastid DNA-dependent RNA Polymerase subunits (Figure 7B) (Pfannschmidt *et al*., 2015). Interacting with those, there are two OGs annotated as encoding TRANSCRIPTION INITIATION FACTORS, TFIIB2 and TFIIE (Table S6) (Knutson, 2013).

**Figure 7.**
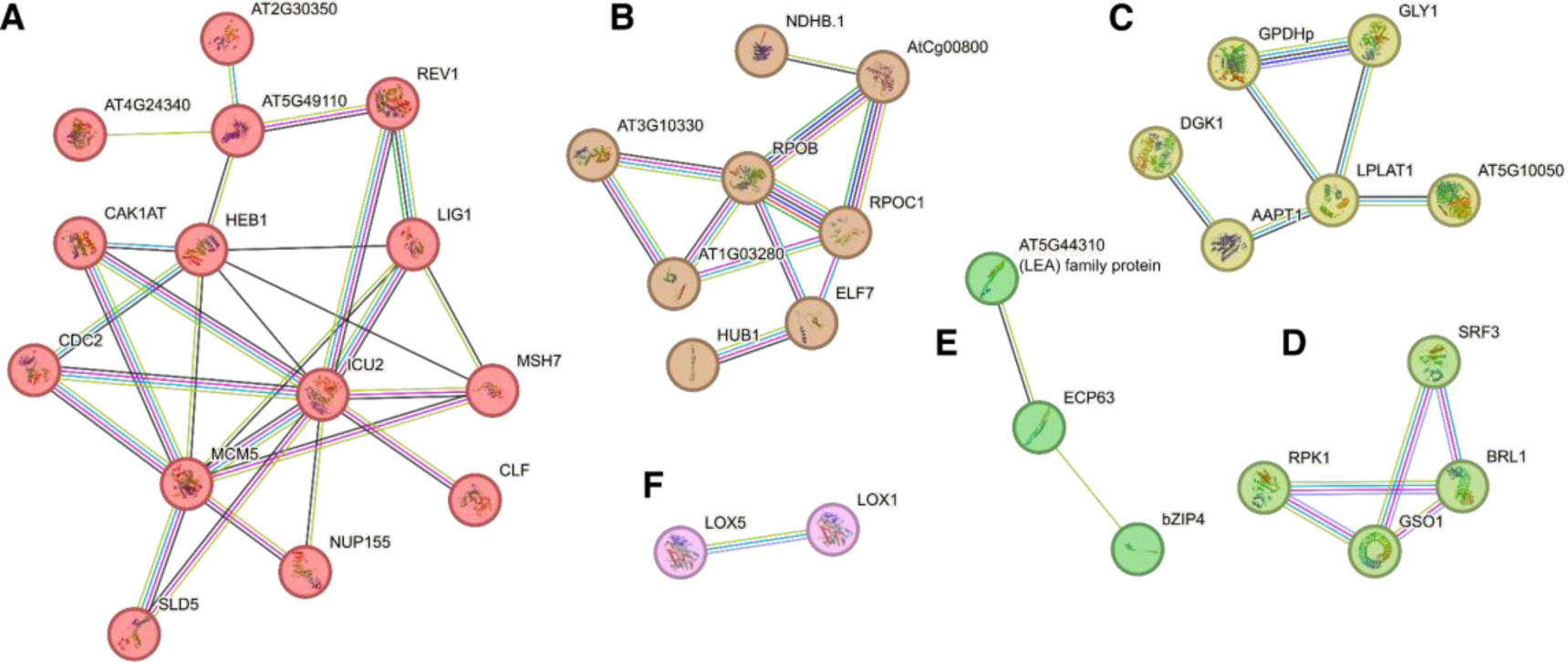
Orthogroup-based putative protein clusters unique to the *Amborella* endosperm transcriptional network. A. Cell cycle and chromosome organization. B. DNA-directed RNA polymerases and associated proteins. C. Glycerolipid biosynthesis. D. Receptor Kinases. E. LEA proteins. F. Lipoxygenases.

Metabolism-related OGs were also found enriched in the *Amborella* endosperm. For instance, there is a large cluster of genes related to lipid biosynthesis (Figure 7C). Moreover, three OGs were annotated as encoding lipoxygenases (LOX; Table S6, Figure 7G), which are typical of seeds and catalyze the oxidation of polyunsaturated fatty acids into functionally diverse oxylipins (Casey, 1999; Porta & Rocha-Sosa, 2002; Wasternack & Feussner, 2018).

Importantly, a set of four OGs annotated as encoding receptors harboring protein kinase domains were found exclusive in *Amborella*. These are STRUBBELIG-RECEPTOR FAMILY 3 (SRF3) (Eyüboglu *et al*., 2007), RECEPTOR-LIKE PROTEIN KINASE 1 (RPK1) (Hong *et al*., 1997), GASSHO1 (GSO1) (Tsuwamoto *et al*., 2008), and the putative brassinosteroid receptor BRI1-LIKE 1 (BRL1) (Zhou *et al*., 2004; Caño-Delgado *et al*., 2004) (Figure 7D). Other hormone-related OGs were also found enriched in the *Amborella* endosperm, namely those related to auxin and ethylene signaling. Some examples are *AUXIN RESPONSE FACTOR 2* (*ARF2*) (Okushima *et al*., 2005), *ETHYLENE-INSENSITIVE3-LIKE 1* (*EIL1*) and *ETHYLENE RESPONSE FACTOR 71* (*ERF71*) (Chao *et al*., 1997; Hess *et al*., 2011) (Table S6).

A specialization of a subset of LEA proteins seems to be shared among all the clades studied. In the case of *Amborella,* OGs encoding LEA proteins were found in the exclusive set of OGs in the endosperm, such as *LEA3* (Battaglia *et al*., 2008) (Figure 7F).

Next, we identified the DEGs that are specific to the nourishing tissues of the *Nymphaea* seeds, the endosperm and the perisperm. At the time of collection, around a week after pollination, the endosperm was significantly smaller than the perisperm, which was filled with starch and oil deposits (Fig. S1) (Povilus *et al*., 2015). Our analyses of differential gene expression between these two tissues showed that the endosperm had more upregulated DEGs (2214) than the perisperm (826), likely reflecting a more active transcriptional state. There were commonalities in the sets of enriched terms in both the perisperm and endosperm, when compared to somatic tissues (Table S7). Shared functions include hormone metabolism and signaling, as illustrated by the enriched terms “ABA-activated signaling pathway”, “Response to auxin” and “Hormone biosynthetic process” (Table S7). Also, development related terms were found enriched in both DEG sets, like “Meristem development” and “Plant organ development” (Table S7). The genes that are privately enriched in each tissue separately give us an overview of their specific functions in the *Nymphaea* seed. The perisperm was enriched in several secondary metabolism related processes, including “Flavonoid biosynthetic process”, “Phenylpropanoid metabolic process” and “Ethylene biosynthetic process”, pointing to a pivotal role for the perisperm in the biosynthesis of hormones and secondary metabolites. Likewise, peroxisome related terms were specific to the perisperm. Roles for the peroxisome in the perisperm may be related to hormone biosynthesis and fatty acid-oil body interactions (Pan et al., 2020). The endosperm had however a larger set of enriched terms correlating with the larger number of DEGs (Table S7). Among them we can highlight those with transcriptional regulation and epigenetic functions, like “Chromatin assembly”, “RNA methylation”, as well as those highlighting specific cell components, such as “Plasmodesmata”, “Plant-type cell wall”, “Plant-type vacuole” and “Microtubule”. These DEGs between endosperm and perisperm clearly indicate a sub-functionalization of these two structures, with the endosperm being enriched in processes related to gene regulation and cellular dynamics, while the perisperm is mostly enriched in metabolism-related processes.

In addition to the DEOGs shared with other Angiosperms (Table S4), we identified transcriptional networks that were specific to the water lily nourishing tissues. We detected 334 OGs that were exclusively DE in the endosperm and perisperm (Hypergeometric Test p-value: 3,12E-09, Figure 3C, Figure S2). The protein interaction network of the *Nymphaea* nourishing tissues displayed proteins involved in chromatin organization (Figure S8, Table S7). This was evidenced by a large cluster that includes OGs encoding proteins with epigenetic roles, like HISTONE DEACETYLASE 9 (HDA9) (Kim *et al*., 2016), PHOTOPERIOD-INDEPENDENT EARLY FLOWERING 1 (PIE1), a chromatin remodeler of the SWI2/SNF2 family (Noh & Amasino, 2003), and the Flowering Locus VE (FVE), an MSI1 family protein and putative subunit of Polycomb Repressive Complexes (PRC) (Pazhouhandeh *et al*., 2011). Members of the Cul4-RING E3 ubiquitin ligase complex are also present in this interaction cluster (Figure 8A), which may reflect interactions of endosperm-specific epigenetic machinery that involves PRC2 and Cul4-RING E3 interactions, as reported in Arabidopsis (Pazhouhandeh *et al*., 2011). Still related to the ubiquitination machinery, we identified a protein cluster with several components of the proteasome (Figure 8D).

**Figure 8.**
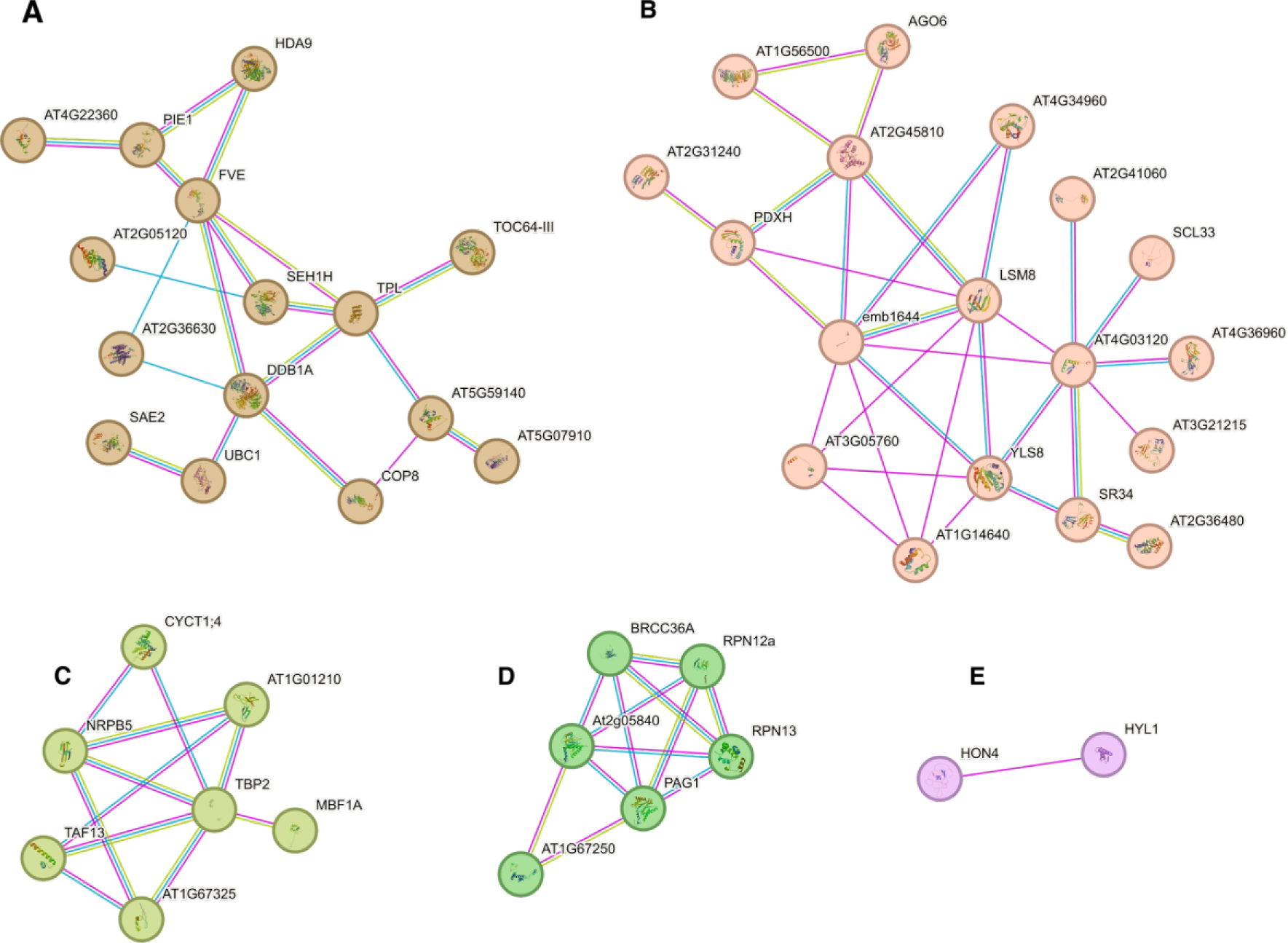
Orthogroup-based putative protein clusters unique to the *Nymphaea* nourishing tissues transcriptional network. A. Chromatin organization. B. Spliceosome. C. RNA Pol II. D. Proteasome. E. RdDM.

Another prominent functional cluster includes members of the spliceosome, driving the enrichment of the term “Interchromatin granule”, which in animal cells is linked to spliceosome activity (Mintz *et al*., 1999). And a further interesting cluster is composed of two OGs representing RNAPs, namely, *DNA-DIRECTED RNA POLYMERASE*, *SUBUNIT M*, and *RNA POLYMERASE II SUBUNIT 5* (*NRPB5*) (Haag *et al*., 2014) (Figure 8C). Interacting with them are three OGs that encode accompanying proteins of the Polymerase II holoenzyme (Figure 8C). Additionally, two OGs putatively involved in RdDM were present in the private *Nymphaea* set, including the linker histone HON4/HYPONASTIC LEAVES 1 (HYL1) and ARGONAUTE 6 (AGO6; Figure 8E) (Vazquez *et al*., 2004; Duan *et al*., 2015).

### Transcriptional Landscapes in Monocots and Eudicots: Shared Features and Unique Patterns

We then proceeded to the realm of core angiosperms, monocots and eudicots. Here, we focus on describing the transcriptional signatures inherent to the endosperms of the species we studied in both clades, elucidating distinctive and common gene expression patterns.

The transcriptional signatures of the rice and maize endosperms showed a strong differentiation to other seed plants, as illustrated by the PCA analysis (Figure 3A). It is worth highlighting that our monocot dataset is restricted to the Poaceae, which displays the large persistent endosperm of cereals (Zheng & Wang, 2015). With the aim to describe its transcriptional network we identified DEOGs exclusively expressed in the monocot endosperms and, secondly, looked for OGs that were absent in rice and maize but present in the eudicots in our study. We thus identified 235 DEOGs unique to the rice and maize endosperms (Hypergeometric test p-value: 6,45E-06, Figure 3C, Figure S2, Table S8). Interestingly, we identified a monocot-exclusive nutrient storage cluster (Figure 9A). At least seven of the OGs encode LEA proteins and dehydrins, which are LEA-type proteins (Dure *et al*., 1989; Artur *et al*., 2019), evidencing a definitive role for these in monocot endosperm nutrient storage. Another important cluster includes many genes involved in processes related to the Krebs cycle and starch biosynthesis, processes that take place in the amyloplast of cereal endosperms (Figure 9B, Figure S9 and Table S8) (Zheng & Wang, 2015). These OGs drive the enrichment of the GO process terms “Starch biosynthetic process”, “Glycogen metabolic process”, and the GO Component term “Amyloplast” (Table S8). Another relevant cluster is related to flavonoid biosynthesis and other secondary metabolites (Figure 9C, Figure S9). The accumulation of flavonoids has been proposed to inhibit auxin transport and modulate seed development (Peer & Murphy, 2007). Most of these OGs had their best Orthofinder assigned target in the *Oryza* genome, supporting their clade specificity (Table S8).

**Figure 9.**
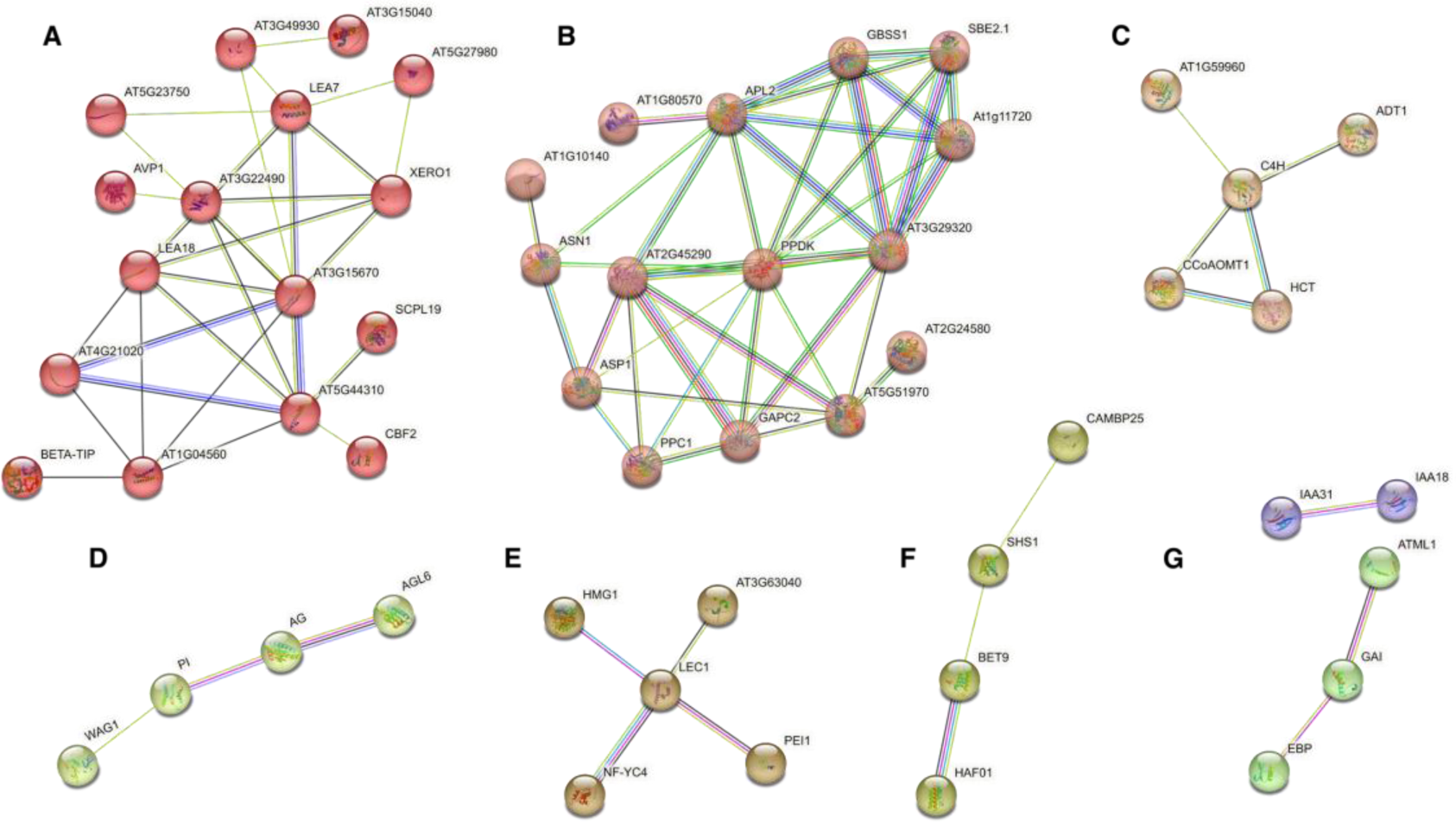
Orthogroup-based putative protein clusters unique to the monocot endosperm transcriptional network. A.Storage. B. Starch biosynthesis / Amyloplast. C. Secondary metabolite biosynthesis. D. MADS-box proteins. E. Other TFs. F. Bromodomain-containing proteins. G. Hormone signaling.

Monocots appear to have a transcriptional regulatory machinery specialized in the endosperm. We found enriched the terms: “Regulation of transcription, DNA-templated” and “DNA-binding transcription factor activity” (Table S8). Moreover, we identified clusters of OGs that encode putative transcription factors and other transcriptional regulators (Figure 9D-F, Figure S9, Table S8). Among those are three MADS domain encoding OGs, and two encoding putative subunits of Nuclear Factor Y, the latter being a best hit for LEAFY COTYLEDON 1 (LEC1) (Kwong *et al*., 2003; Song *et al*., 2021). Among other sets of transcriptional regulators found in this OG set, are bromodomain containing proteins which likely function as epigenetic regulators (Jarończyk *et al*., 2021): two OGs encoding bromodomain-containing proteins, the histone acetyltransferase TBP-ASSOCIATED FACTOR 1 (HAF01) and BROMODOMAIN AND EXTRATERMINAL DOMAIN PROTEIN 9 (BET9) (Pandey, 2002), were found exclusively in the monocot endosperm (Figure 9F).

Additionally, we found OGs related to hormone signaling in the monocot nourishing tissue (Figure 9G): two OGs corresponding to the INDOLE-3-ACETIC ACID INDUCIBLE (IAA) family of proteins, implicated in auxin signaling (Nemhauser, 2018), and OGs related to gibberellin and ethylene signaling, like GIBBERELLIC ACID INSENSITIVE (GAI) (Peng *et al*., 1997), and ETHYLENE-RESPONSIVE ELEMENT BINDING PROTEIN (EBP) (Büttner & Singh, 1997).

Likewise, we detected 187 DEOGs that were exclusive to the eudicot endosperms (Hypergeometric test p-value: 8,10E-03, Figure 3C, Figure S2; Table S9). The term “Microtubule cytoskeleton” was enriched in this DEOG set. Genes that drive its enrichment and that appear to form a cluster of interacting proteins include those encoding three mitosis specific kinesins (KIN), KIN5A, KIN7A and KIN14D (Vanstraelen *et al*., 2006), as well as other important components of the cytoskeleton (Figure 10A, Figure S20). Specialization of the replication and transcription machinery in the eudicot endosperm is also evidenced by the presence of two related protein clusters (Figure 10B-C). One of them encompasses DEOGs involved in DNA replication (Figure 10B), encoding subunits of origin-of-replication complexes, ORC1 and ORC2 (Collinge *et al*., 2004; Vergara *et al*., 2023). Likewise, POLA2, an alpha subunit of the DNA polymerase (Yang *et al*., 2009), and EXO1, an exonuclease (Kazda *et al*., 2012), were present in this replication cluster (Figure 10B). The other cluster is composed of several DEOGs that are annotated as subunits of DNA-directed RNAPs and their accompanying proteins (Figure 10C). This includes for example: NRPD3B, NRPD5B, RPA12-like and RPA3A (Larkin *et al*., 1999; Ream *et al*., 2009; Aklilu *et al*., 2020). Additionally, the term “Condensed chromosome” was found enriched. Its corresponding DEOGs encode a cluster of interacting proteins, which include ASYNAPTIC1 (ASY1) (Armstrong *et al*., 2002), and the SWI/SNF member RAD54 (Osakabe *et al*., 2006) (Figure 10D). Other DEOGs of importance are annotated as encoding transcription factors, including AGAMOUS-like 18 (AGL18) (Adamczyk *et al*., 2007), PHAVOLUTA (ATHB-9/PHV) (Prigge *et al*., 2005), GLABRA 2 (GL2) (Masucci *et al*., 1996), and REGULATOR OF AXILLARY MERISTEMS 2 (RAX2) (Müller *et al*., 2006) (Table S9).

**Figure 10.**
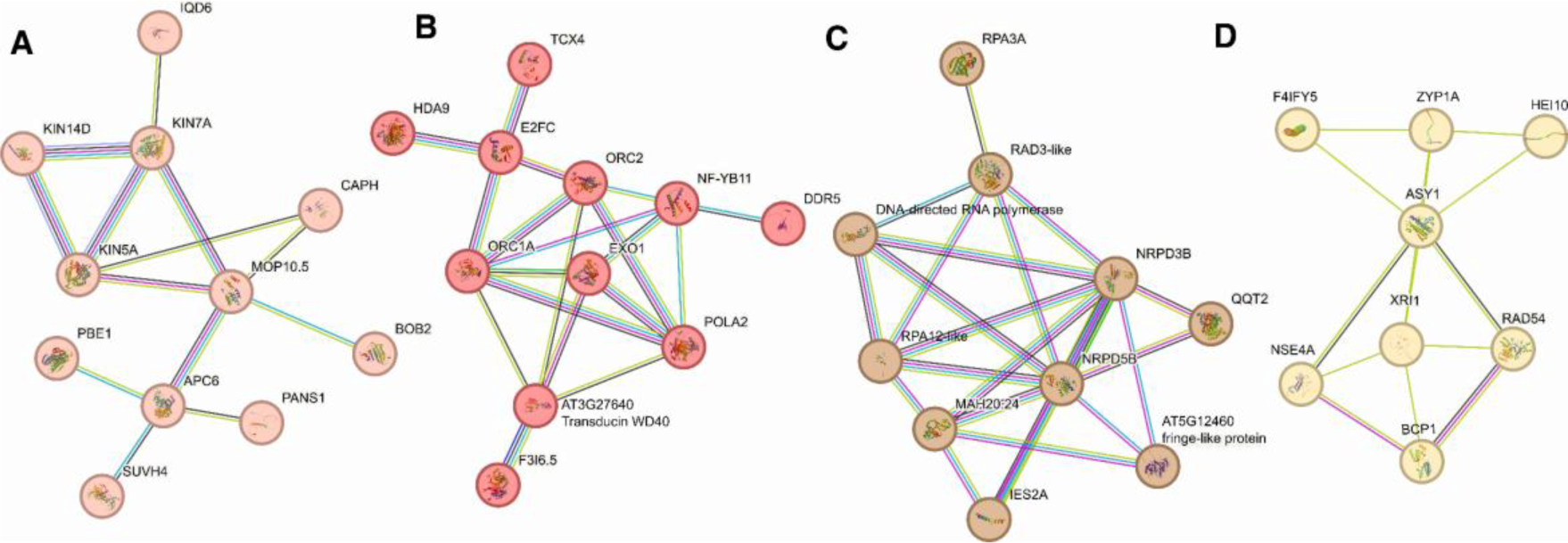
Orthogroup-based putative protein clusters unique to the eudicot endosperm transcriptional network. A.Cytoskeleton. B. Replication. C. Transcription. D. Chromosome condensation.

With the aim to detect differences between monocots and eudicots we identified non-exclusive DEOGs sets of the monocot and eudicot taxa. We found 568 DEOGs shared between maize and rice and 574 DEOGs shared between the eudicots. We performed set operations to disentangle the differences in the transcriptomic landscape of those taxa. This analysis clarifies angiosperm differences, disregarding patterns predating the core angiosperms that might obscure diversification within clades, if gymnosperms and early divergent angiosperm DEOG sets were included. With the aim to determine which DEOGs were specific to each of the two main angiosperm groups, monocots and eudicots, we determined the complement DEOG sets of both clades. A total of 464 DEOGs were found in eudicots and not in monocots (Table S9). Replicating the trend of the exclusive set of DEOGs of core eudicots, we found five DEOGs annotated as subunits of RNAPs, as well as nine members of the kinesin complex. Although microtubule cytoskeleton proteins were part of the conserved transcriptional network of the angiosperm seed nourishing tissue (Table S4), the large diversification of kinesins was observed only in the core eudicots and not in monocots. The larger number of DEOGs belonging to these families, in this broader comparison (non-exclusive to other taxa), indicates that these OGs likely share ancestry with non-core angiosperm taxa such as gymnosperms and early divergent groups, rather than the more closely related monocots. This suggests an ancestral evolutionary role in the nourishing tissues for these OGs and their diversification within the eudicot lineage. Other DEOGs show the same trend of increased diversification in the eudicots in comparison to monocots. For example, DEOGs encoding five ABC-type transporters (Do *et al*., 2018), two WUSCHEL RELATED HOMEOBOX (WOX) TFs (Van Der Graaff *et al*., 2009), and three DNA repair RAD helicases (Hernandez Sanchez-Rebato *et al*., 2021), were found only in eudicots and not in either monocot species that we studied (Table S9).

Comparing only the DEOGs of core angiosperms, we then identified 459 monocot-specific DEOGs, including encoding families such as ATL RING-H2 finger proteins (Serrano *et al*., 2006), ethylene-responsive factors (ERFs) (Müller & Munné-Bosch, 2015), Subtilisin-like proteases (SBTs) (Rautengarten *et al*., 2008), UDP-glycosyltransferases (UGTs), previously implicated in grain size in rice (Dong *et al*., 2020) , and ZAT family TFs (ZATs) (Englbrecht *et al*., 2004), showcasing examples of diversifying evolution within monocots. These families were notably absent in the core eudicots complement set of DEOGs (Table S8).

## Discussion

The evolutionary advent of seeds is one of the factors that allowed spermatophytes to reproduce over long distances and independently of the presence of water and, therefore, to occupy new ecological niches (Pettitt, 1970; Brenner & Stevenson, 2006). In part, this is due to the protection of the embryo against environmental conditions by the sporophytic tissues of the maternal plant (Pettitt, 1970; Linkies *et al*., 2010; Huss & Gierlinger, 2021). Moreover, seeds allow for a state of dormancy, only germinating when the conditions are favorable (Debeaujon *et al*., 2000; Willis *et al*., 2014). But a major advance of the seed habit is the initial development of the new sporophytic generation within the maternal tissues, where it is nourished by a specialized structure (Hardev, 1978; Baroux *et al*., 2002; Brenner & Stevenson, 2006; Linkies *et al*., 2010). Interestingly, already in seed ferns there were structures with a nourishing function, probably derived from the megagametophyte (Doyle, 2006; Spencer *et al*., 2013). In the gymnosperms the nourishing function is carried out by the proliferating megagametophyte, while in the angiosperms, this is done in most cases by the endosperm (Baroux *et al*., 2002; Linkies *et al*., 2010). The main difference being that the nourishing structure of the gymnosperms develops autonomously, i.e., its proliferation is not coupled to a fertilization event, while the development of the angiosperm endosperm is coupled to fertilization (Linkies *et al*., 2010). This coupling has obvious advantages, such as ensuring that nutrients are not allocated to “empty” seeds (without embryos), but it also allows for a coordinated development between the embryo and the endosperm (Lafon-Placette & Köhler, 2014; Doll & Ingram, 2022). Nevertheless, we hypothesized that some transcriptomic signatures should be conserved between the nourishing tissues of gymnosperms and those of the angiosperms, as both fulfill similar functions. Although transcriptomes of gymnosperm megagametophytes have been recently described in the literature, those samples originated from either mature ovules of *Ginkgo biloba* (Zumajo-Cardona *et al*., 2021), or germinating seeds of *Pinus sylvestris* (Cervantes *et al*., 2021). As our goal was to assess transcriptomic signatures at early stages of seed development, we generated novel RNAseq datasets of megagametophytes of *Pinus pinaster* obtained from young developing seeds. While indeed there were shared signatures between those megagametophytes and the nourishing tissues of the angiosperms, we did observe several pathways enriched specifically in the gymnosperm samples. This fits with previous observations made during ovule development in *Ginkgo biloba* and in *Gnetum gnemon* (Zumajo-Cardona & Ambrose, 2021; Zumajo-Cardona *et al*., 2021). While some genes, like *WUSCHEL* (*WUS*), likely have a conserved function during ovule development in *Arabidopsis* and *Gnetum* (Groß-Hardt *et al*., 2002; Zumajo-Cardona & Ambrose, 2021), there is no conservation of the tissue-specific expression of several other regulators of ovule development (Zumajo-Cardona & Ambrose, 2021; Zumajo-Cardona *et al*., 2021). Interestingly, although the *WUS* homologue is expressed specifically in the nucellus in *Arabidopsis* and in *Gnetum* (Groß-Hardt *et al*., 2002; Zumajo-Cardona & Ambrose, 2021), it is also expressed in the integuments in *Ginkgo* (Zumajo-Cardona *et al*., 2021). Thus, while our analyses in *Pinus* can provide some foundation to understand the development of the megagametophyte during seed development in gymnosperms, there is certainly a great deal of variation within that clade, reflecting its rich evolutionary history. The same applies to our analyses of the unique features of the angiosperm species studied here. While some transcriptomic signatures are surely characteristic of the clades to which each species belongs, others are likely species-specific.

In most angiosperms the nourishing function in the seed is fulfilled by the endosperm, however, in some species it can also be fulfilled by the perisperm, which originates from the sporophytic nucellus of the ovule. This is the case in *Nymphaea*, which we studied here, and in species of Amaranthaceae, like amaranth and quinoa (Burrieza *et al*., 2014; Povilus *et al*., 2015), and of the Malpighiaceae (Souto & Oliveira, 2014). Interestingly, it seems that in endospermic seeds, like those of Arabidopsis, the endosperm develops antagonistically with the nucellus, instructing its degeneration (Xu *et al*., 2016). Thus, in seeds where the two structures co-exist, it is reasonable to hypothesize that they carry out complementary functions. This is supported by our comparative analysis of DEOGs between the endosperm and the perisperm of *Nymphaea*. Indeed, while the perisperm is mostly enriched in metabolism-related processes, the endosperm is enriched in gene regulation and cell-related processes, likely linked to its proliferative nature and it being a site for parental conflict (Povilus *et al*., 2018; Köhler *et al*., 2021; Montgomery & Berger, 2021). This suggests that in perispermic seeds, there is a subfunctionalization of these two structures, whereas in endospermic species these functions are fully fulfilled by the endosperm.

The fact that the seed nourishing structures of different clades share common functionalities, suggests that they probably also share some transcriptomic signatures. However, it is also expected that there are signatures that are specific to certain clades, since there is significant divergence in the form and function of some of those structures. This hypothesis is supported by our data, which shows that certain gene expression signatures are conserved throughout evolution, while others are unique to the nourishing structures of species within each clade.

### Conserved transcriptional features of the angiosperm nourishing tissues

Seed storage proteins are a vital component of plant seeds and play a crucial role in ensuring the successful germination and growth of new plants (Shewry *et al*., 1995; Shewry & Halford, 2002; Liu *et al*., 2022). OGs encompassing genes that encode storage proteins were some of the main drivers of the differentiation of the nourishing tissue transcriptomes. Proteins assembled in these OGs and consistently present in all angiosperms studied are annotated as cruciferins, cupins and glutenins (Payne *et al*., 1982; Dunwell, 1998; Dunwell *et al*., 2004; Nietzel *et al*., 2013), but also as caleosins and peroxygenases, which are involved in the biogenesis and maintenance of lipid bodies (Poxleitner *et al*., 2006; Meesapyodsuk & Qiu, 2011). LEA proteins, although not fulfilling a specific storage function, were also part of the conserved angiosperm nutrient storage cluster. These proteins primarily function to safeguard cellular components and maintain seed viability under desiccation and dehydration stress (Cuming, 1999; Battaglia *et al*., 2008). Interestingly, diversification of LEA proteins in the monocot endosperm was evidenced in our analyses. It is important to note that some overlap or interactions between LEA proteins and nutrient storage components may exist (Dirk *et al*., 2020). For instance, during seed desiccation and maturation, LEA proteins may interact with and stabilize storage proteins and lipids, helping to maintain their integrity and functionality (Cuming, 1999; Dirk *et al*., 2020). Another group of seed proteins enriched in our analyses were lipoxygenases (LOX), which were first discovered in legumes and are prominent seed proteins (Casey, 1999; Porta & Rocha-Sosa, 2002). LOX-generated oxylipins, particularly JA and its derivatives, can modulate seed dormancy by influencing hormone signaling pathways and by interacting with other dormancy-related regulators (Casey, 1999; Chauvin *et al*., 2013). LOX specialization in the nourishing tissue as revealed by our analyses was not a conserved feature of angiosperms but rather appeared to be an ancestral trait present only in gymnosperms and in *Amborella*. Instead of playing a critical role in core angiosperm endosperms, LOX proteins seem to play roles in plant defense, development and stress responses in vegetative tissues (Chauvin *et al*., 2013).

A protein cluster involved in cell cycle and nucleosome assembly was also a highlight of the clusters shared by angiosperms. Strikingly, OGs annotated as histone variants were enriched in all angiosperm clades. Histone variants contribute to the definition of chromatin states and gene expression patterns (Borg *et al*., 2021; Foroozani *et al*., 2022; Bhagyshree *et al*., 2023). By modifying chromatin structure, they regulate the accessibility of DNA, influence gene expression, contribute to epigenetic inheritance, and participate in plant responses to environmental cues and developmental processes (Borg *et al*., 2021; Bhagyshree *et al*., 2023). Interestingly, histone variants were exclusively present in the shared protein network and not in any of the uniquesets of proteins specific to particular plant clades. This result may signify that the identified histone variant OGs represent the entire range of variants that play crucial, and therefore evolutionarily conserved, roles in angiosperm endosperms. Interestingly, genes encoding histone variants have been shown to be determinant for parental effects during embryogenesis in mammals (Molaro *et al*., 2020). Moreover, histone variants have been shown to correlate with different chromatin states (Borg *et al*., 2021), and, for instance, the paternal expression of an H3 variant which is insensitive to PcG function is required for reprogramming of the paternal germline (Borg *et al*., 2020). This is particularly relevant because the angiosperm endosperm is a site for genomic conflict between the mother and the father, and the chromatin landscape of the endosperm underlies reproductive barriers in angiosperms (Jiang *et al*., 2017). Thus, it is tempting to hypothesize that the differential expression of histone variants in the endosperm correlates with the arisal of a parental conflict for nutrient allocation.

### Unique transcriptional networks shed light into specializations within clades

The sets of unique transcriptional networks in species of each major clade of seed plants potentially represent genes that contributed to specialization within those clades. Genes that are expressed in only a subset of plants within a restricted phylogenetic group should be considered as potential candidates for functional studies. The unique network of the *Amborella* endosperm revealed OGs involved in epigenetic processes. Similarly, the private network of *Nymphaea* consists of OGs that are annotated as RNA polymerases and putative PcG components. These results highlight the evolutionary divergence of these taxa from core angiosperms and precisely pinpoint the differences in the epigenetic machinery that is active in their endosperms. In Arabidopsis, components of the FERTILIZATION-INDEPENDENT SEED PRC2 (FIS-PRC2) are specific to the gametophyte and the endosperm (Luo *et al*., 2000; Chaudhury *et al*., 2001; Ungru *et al*., 2008). However, their expression progressively declines during seed development: after a few mitotic rounds, FIS-PRC2 components like FIS2, MEA and FIE become undetectable in the endosperm or limited to the chalazal cyst (Luo *et al*., 2000). Given that we do not detect differential expression of PRC2-encoding genes in the endosperms of other eudicots and monocots, it is likely that those genes are also downregulated after fertilization, like in Arabidopsis. However, the same may not be true for early diverging lineages, where we do detect PcG-encoding genes expressed in the endosperm. This suggests that constitutive PRC2 activity in the endosperm may be the ancestral state, and its downregulation during endosperm formation arose later during angiosperm speciation. An evident hallmark of the nourishing tissue specialization was the biosynthesis of secondary metabolites. For example, the term “Biosynthetic process” was enriched in the exclusive sets of pine and in monocots. Regarding the transcriptional network of pine, it demonstrated a wide range of biosynthetic pathways related to hormone signaling, such as JA, and to structural characteristics of the seed. JA is derived from linolenic acid (Wasternack & Strnad, 2018) and, together with other phytohormones, like auxin and ABA, it has been shown to affect seed germination in Arabidopsis (Pan *et al*., 2020a; Mei *et al*., 2023). Furthermore, flavonoids have been suggested to play a regulatory role in phytohormone signaling (Appelhagen *et al*., 2011; Li & Zachgo, 2013; Brunetti *et al*., 2018). It is tempting to hypothesize that the specific proteins assembled in the OGs identified with secondary metabolite biosynthesis in monocots and pine may act as regulators of phytohormone signaling in their nourishing tissues.

The persistent nature of the Poaceae cereal endosperm leaves a distinct mark on their exclusive transcriptional network. A large protein cluster involved in processes that take place in the amyloplast of cereal endosperms, such as starch biosynthesis, was conspicuous in our analysis. We also identified two OG clusters annotated as encoding TFs in the private transcriptional network of monocots. This highlights their functional role and possibly signals the diversification of these TFs in the monocot endosperm. The MADS-box and AP2-B3 protein families are well-studied transcription factor families that play important roles in regulating endosperm development (Lu *et al*., 2012; Batista *et al*., 2019; Yang *et al*., 2020; Song *et al*., 2021). Although we identified a single DEOG grouping a large set of AGAMOUS-LIKE proteins, AGL9, in the shared set of expressed OGs among all angiosperms, and AGL18 in the unique transcriptional network of the eudicot endosperm, we did not observe any strong trend of diversification of OGs containing these TFs in the various endosperm of angiosperms. The pattern may be restricted to monocots, in which we identified three MADS domain containing DEOGs. This could suggest that remaining OGs encompassing additional MADS-box TFs have functional roles throughout the entire plant body and indeed play roles in the endosperms, but not specifically in it. Alternatively, it is very likely that genes encoding endosperm-specific MADS-box TFs were no longer expressed in the samples that we analyzed. Type I MADS-box TFs have been shown to play prominent roles in endosperm development, and their downregulation at the time of endosperm cellularization is crucial for this developmental transition (Erilova *et al*., 2009; Hehenberger *et al*., 2012; Batista *et al*., 2019). Because the datasets that we analyzed originated in endosperms that were in early developmental stages, either cellular or close to undergoing cellularization, this likely explains why this family of genes is not prominent in our analysis.

Our work provides the first comparative transcriptomic analysis of early seed nourishing tissues, which includes representatives of all main angiosperm clades. We also provide the first transcriptomic datasets of isolated endosperms of basal angiosperms. In addition to the newly generated datasets, we present an orthogroup database encompassing the main phylogenetic plant clades. These catalogs (unique and shared sets) will streamline the selection of candidate genes for functional genetic studies in nourishing tissues across the plant kingdom, enabling targeted investigations and enhancing our understanding of gene function and evolution.

## Acknowledgements

We thank Bernhard Reiken at the Bonn botanical Garden for providing the *Amborella* materials, and Ingo Kallemeier and Michael Burkart at the Potsdam Botanical Garden for the *Nymphaea* materials. We are grateful to Barbara Schmidt and Prof. Julien Bachelier at the Berlin Botanical Garden for guidance with *Nymphaea* material collection. We also thank Arun Sampathkumar and Anja Froehlich for help with the LCM, and Andreia Rodrigues for help with *Pinus* megagametophyte dissection.

This work was funded by a Humboldt Postdoctoral Fellowship to AMFR, by the Max Planck Society to DDF, and by Fundação para a Ciência e a Tecnologia to CMM (IF/01168/2013).

## Author contributions

AMFR and DDF designed the research. AMFR, CMM and DDF collected the plant materials. AMFR performed the experiments and analyzed the data. AMFR wrote the first draft of the manuscript, and all authors contributed to and approved the final version.

## Data statement

Newly generated data for the endosperms and leaves of *Nymphaea caerulea* and *Amborella trichopoda* as well as the megagametophyte transcriptomes of *Pinus pinaster* have been posted in the NCBI SRA database under submission SUB13828753. All other data used in this manuscript is publicly available and their sources are described in Table S1.

## Supplementary Information

### Methods S1

**Table S1. Data provenance, tissue details, number of total mapped reads, total number of genes detected, DEGs and DEOGs.**

**Table S2. Orthogroup description.**

**Table S3. Results of differential gene expression between nourishing tissues and leaves of all species studied.**

**Table S4. Conserved transcriptional network of the angiosperm seed nourishing tissue.**

**Table S5. The transcriptional network of the *Pinus pinaster* megagametophyte.**

**Table S6. The private transcriptional network of the *Amborella trichopoda* endosperm.**

**Table S7 – The transcriptional network of the *Nymphaea caerulea* seed.**

**Table S8. The monocot endosperm transcriptional network.**

**Table S9. The transcriptional network of the eudicot endosperm.**

**Figure S1. Examples of seeds of *Amborella trichopoda* and *Nymphaea caerulea* used for microdissection and transcriptome generation.**

**Figure S2. Details of PCA analyses and Upset plot of DEOGs showing intersections**

**Figure S3. The conserved transcriptional network of the angiosperm endosperm.**

**Figure S4. The conserved transcriptional network of the angiosperm endosperm, the cell cycle and nucleosome architecture interacting network**

**Figure S5. Conserved transcriptional network of the angiosperm seed nourishing tissue, the Nourishing cluster protein interacting network**

**Figure S6. The *Pinus pinaster* megagametophyte private protein network.**

**Figure S7. The *Amborella* endosperm private protein network.**

**Figure S8. The *Nymphaea caerulea* endosperm private protein network.**

**Figure S9. The monocot endosperm private protein network.**

**Figure S10. The eudicot endosperm private protein network.**

## References

1. Adamczyk BJ, Lehti-Shiu MD, Fernandez DE. 2007. The MADS domain factors AGL15 and AGL18 act redundantly as repressors of the floral transition in Arabidopsis. The Plant Journal 50: 1007–1019.

2. Aklilu BB, Peurois F, Saintomé C, Culligan KM, Kobbe D, Leasure C, Chung M, Cattoor M, Lynch R, Sampson L, et al. 2020. Functional Diversification of Replication Protein A Paralogs and Telomere Length Maintenance in Arabidopsis. Genetics 215: 989–1002.

3. Amborella Genome Project. 2013. The Amborella Genome and the Evolution of Flowering Plants. Science 342: 1241089.

4. Appelhagen I, Lu G-H, Huep G, Schmelzer E, Weisshaar B, Sagasser M. 2011. TRANSPARENT TESTA1 interacts with R2R3-MYB factors and affects early and late steps of flavonoid biosynthesis in the endothelium of Arabidopsis thaliana seeds. The Plant Journal 67: 406–419.

5. Armstrong SJ, Caryl AP, Jones GH, Franklin FCH. 2002. Asy1, a protein required for meiotic chromosome synapsis, localizes to axis-associated chromatin in *Arabidopsis* and *Brassica*. Journal of Cell Science 115: 3645–3655.

6. Artur MAS, Zhao T, Ligterink W, Schranz E, Hilhorst HWM. 2019. Dissecting the Genomic Diversification of Late Embryogenesis Abundant (LEA) Protein Gene Families in Plants (Y Van De Peer, Ed.). Genome Biology and Evolution 11: 459–471.

7. Baroux C, Spillane C, Grossniklaus U. 2002. Evolutionary origins of the endosperm in flowering plants. Genome Biology 3: reviews1026.1.

8. Barrero JM, González-Bayón R, Del Pozo JC, Ponce MR, Micol JL. 2007. *INCURVATA2* Encodes the Catalytic Subunit of DNA Polymerase α and Interacts with Genes Involved in Chromatin-Mediated Cellular Memory in *Arabidopsis thaliana*. The Plant Cell 19: 2822–2838.

9. Batista RA, Moreno-Romero J, Qiu Y, van Boven J, Santos-González J, Figueiredo DD, Köhler C. 2019. The mads-box transcription factor pheres1 controls imprinting in the endosperm by binding to domesticated transposons. eLife 8.

10. Battaglia M, Olvera-Carrillo Y, Garciarrubio A, Campos F, Covarrubias AA. 2008. The Enigmatic LEA Proteins and Other Hydrophilins. Plant Physiology 148: 6– 24.

11. Bhagyshree J J LZ, Elin A, Akihisa O, Vikas S, Ramesh Y, Svetlana A, Luisa KA, Frédéric B. 2023. Histone variants shape the chromatin states in Arabidopsis. eLife 12.

12. Bird D, Beisson F, Brigham A, Shin J, Greer S, Jetter R, Kunst L, Wu X, Yephremov A, Samuels L. 2007. Characterization of Arabidopsis ABCG11/WBC11, an ATP binding cassette (ABC) transporter that is required for cuticular lipid secretion ^†^. The Plant Journal 52: 485–498.

13. Borg M, Jacob Y, Susaki D, LeBlanc C, Buendía D, Axelsson E, Kawashima T, Voigt P, Boavida L, Becker J, et al.2020. Targeted reprogramming of H3K27me3 resets epigenetic memory in plant paternal chromatin. Nature Cell Biology 22: 621–629.

14. Borg M, Jiang D, Berger F. 2021. Histone variants take center stage in shaping the epigenome. Current Opinion in Plant Biology 61: 101991.

15. Brenner ED, Stevenson D. 2006. Using Genomics to Study Evolutionary Origins of Seeds. In: Williams Claire G, ed. Managing Forest Ecosystems. Landscapes, Genomics and Transgenic Conifers. Dordrecht: Springer Netherlands, 85–106.

16. Brunetti C, Fini A, Sebastiani F, Gori A, Tattini M. 2018. Modulation of Phytohormone Signaling: A Primary Function of Flavonoids in Plant–Environment Interactions. Frontiers in Plant Science 9.

17. Burrieza HP, López-Fernández MP, Maldonado S. 2014. Analogous reserve distribution and tissue characteristics in quinoa and grass seeds suggest convergent evolution. Frontiers in Plant Science 5.

18. Büttner M, Singh KB. 1997. *Arabidopsis thaliana* ethylene-responsive element binding protein (AtEBP), an ethylene-inducible, GCC box DNA-binding protein interacts with an ocs element binding protein. Proceedings of the National Academy of Sciences 94: 5961–5966.

19. Callis J, Carpenter T, Sun CW, Vierstra RD. 1995. Structure and evolution of genes encoding polyubiquitin and ubiquitin-like proteins in Arabidopsis thaliana ecotype Columbia. Genetics 139: 921–939.

20. Caño-Delgado A, Yin Y, Yu C, Vafeados D, Mora-García S, Cheng J-C, Nam KH, Li J, Chory J. 2004. BRL1 and BRL3 are novel brassinosteroid receptors that function in vascular differentiation in *Arabidopsis*. Development 131: 5341–5351.

21. Caro E, Stroud H, Greenberg MVC, Bernatavichute YV, Feng S, Groth M, Vashisht AA, Wohlschlegel J, Jacobsen SE. 2012. The SET-Domain Protein SUVR5 Mediates H3K9me2 Deposition and Silencing at Stimulus Response Genes in a DNA Methylation–Independent Manner (S Grewal, Ed.). PLoS Genetics 8: e1002995.

22. Casey R. 1999. Lipoxygenases. In: Shewry PR, Casey R, eds. Seed Proteins. Dordrecht: Springer Netherlands, 685–708.

23. Cervantes S, Vuosku J, Pyhäjärvi T. 2021. Atlas of tissue-specific and tissue-preferential gene expression in ecologically and economically significant conifer *Pinus sylvestris*. PeerJ 9: e11781.

24. Chao Q, Rothenberg M, Solano R, Roman G, Terzaghi W, Ecker† JR. 1997. Activation of the Ethylene Gas Response Pathway in Arabidopsis by the Nuclear Protein ETHYLENE-INSENSITIVE3 and Related Proteins. Cell 89: 1133–1144.

25. Chaudhury AM, Koltunow A, Payne T, Luo M, Tucker MR, Dennis ES, Peacock WJ. 2001. Control of Early Seed Development. Annual Review of Cell and Developmental Biology 17: 677–699.

26. Chauvin A, Caldelari D, Wolfender J-L, Farmer EE. 2013. Four 13-lipoxygenases contribute to rapid jasmonate synthesis in wounded Arabidopsis thaliana leaves: a role for lipoxygenase 6 in responses to long-distance wound signals. New Phytologist 197: 566–575.

27. Chen F, Liu X, Yu C, Chen Y, Tang H, Zhang L. 2017a. Water lilies as emerging models for Darwin’s abominable mystery. Horticulture Research 4.

28. Chen L-Q, Luo J-H, Cui Z-H, Xue M, Wang L, Zhang X-Y, Pawlowski WP, He Y. 2017b. *ATX3* , *ATX4* , and *ATX5* Encode Putative H3K4 Methyltransferases and Are Critical for Plant Development. Plant Physiology 174: 1795–1806.

29. Cheng Y, Dai X, Zhao Y. 2007. Auxin synthesized by the YUCCA flavin monooxygenases is essential for embryogenesis and leaf formation in Arabidopsis. Plant Cell 19: 2430–2439.

30. Collinge MA, Spillane C, Köhler C, Gheyselinck J, Grossniklaus U. 2004. Genetic Interaction of an Origin Recognition Complex Subunit and the *Polycomb* Group Gene *MEDEA* during Seed Development[W]. The Plant Cell 16: 1035–1046.

31. Culligan KM, Hays JB. 2000. Arabidopsis MutS Homologs—AtMSH2, AtMSH3, AtMSH6, and a Novel AtMSH7—Form Three Distinct Protein Heterodimers with Different Specificities for Mismatched DNA. The Plant Cell 12: 991–1002.

32. Cuming AC. 1999. LEA Proteins. In: Shewry PR, Casey R, eds. Seed Proteins. Dordrecht: Springer Netherlands, 753–780.

33. Debeaujon I, Leon-Kloosterziel KM, Koornneef M. 2000. Influence of the testa on seed dormancy, germination, and longevity in Arabidopsis. Plant Physiology 122: 403– 413.

34. D’Erfurth I, Le Signor C, Aubert G, Sanchez M, Vernoud V, Darchy B, Lherminier J, Bourion V, Bouteiller N, Bendahmane A, et al.2012. A role for an endosperm-localized subtilase in the control of seed size in legumes. New Phytologist 196: 738–751.

35. Dirk LMA, Abdel CG, Ahmad I, Neta ICS, Pereira CC, Pereira FECB, Unêda-Trevisoli SH, Pinheiro DG, Downie AB. 2020. Late Embryogenesis Abundant Protein–Client Protein Interactions. Plants 9: 814.

36. Do THT, Martinoia E, Lee Y. 2018. Functions of ABC transporters in plant growth and development. Current Opinion in Plant Biology 41: 32–38.

37. Doll NM, Ingram GC. 2022. Embryo–Endosperm Interactions. Annual Review of Plant Biology 73: 293–321.

38. Dong N-Q, Sun Y, Guo T, Shi C-L, Zhang Y-M, Kan Y, Xiang Y-H, Zhang H, Yang Y-B, Li Y-C, et al. 2020. UDP-glucosyltransferase regulates grain size and abiotic stress tolerance associated with metabolic flux redirection in rice. Nature Communications 11: 2629.

39. Douglas CJ. 1996. Phenylpropanoid metabolism and lignin biosynthesis: from weeds to trees. Trends in Plant Science 1: 171–178.

40. Doyle JA. 2006. Seed ferns and the origin of angiosperms. The Journal of the Torrey Botanical Society 133: 169–209.

41. Duan C, Zhang H, Tang K, Zhu X, Qian W, Hou Y, Wang B, Lang Z, Zhao Y, Wang X, et al. 2015. Specific but interdependent functions for *A rabidopsis* AGO 4 and AGO 6 in RNA -directed DNA methylation. The EMBO Journal 34: 581–592.

42. Dunwell JM. 1998. Cupins: A New Superfamily of Functionally Diverse Proteins that Include Germins and Plant Storage Proteins. Biotechnology and Genetic Engineering Reviews 15: 1–32.

43. Dunwell JM, Culham A, Carter CE, Sosa-Aguirre CR, Goodenough PW. 2001. Evolution of functional diversity in the cupin superfamily. Trends in Biochemical Sciences 26: 740–746.

44. Dunwell JM, Purvis A, Khuri S. 2004. Cupins: the most functionally diverse protein superfamily? Phytochemistry 65: 7–17.

45. Dure L, Crouch M, Harada J, Ho T-HD, Mundy J, Quatrano R, Thomas T, Sung ZR. 1989. Common amino acid sequence domains among the LEA proteins of higher plants. Plant Molecular Biology 12: 475–486.

46. Dziasek K, Santos-González J, Qiu Y, Rigola D, Nijbroek K, Köhler C. 2023. Dosage sensitive maternal siRNAs determine hybridization success in Capsella. In Review.

47. Emms DM, Kelly S. 2019. OrthoFinder: Phylogenetic orthology inference for comparative genomics. Genome Biology 20.

48. Emonet A, Hay A. 2022. Development and diversity of lignin patterns. Plant Physiology 190: 31–43.

49. Englbrecht CC, Schoof H, Böhm S. 2004. Conservation, diversification and expansion of C2H2 zinc finger proteins in the Arabidopsis thaliana genome. BMC Genomics 5: 39.

50. Erilova A, Brownfield L, Exner V, Rosa M, Twell D, Scheid OM, Hennig L, Kohler C. 2009. Imprinting of the polycomb group gene MEDEA serves as a ploidy sensor in Arabidopsis. PLoS Genetics 5: e1000663.

51. Eyüboglu B, Pfister K, Haberer G, Chevalier D, Fuchs A, Mayer KF, Schneitz K. 2007. Molecular characterisation of the STRUBBELIG-RECEPTOR FAMILY of genes encoding putative leucine-rich repeat receptor-like kinases in Arabidopsis thaliana. BMC Plant Biology 7: 16.

52. Fang X, Qi Y. 2016. RNAi in Plants: An Argonaute-Centered View. The Plant Cell 28: 272–285.

53. Ferrari C, Proost S, Janowski M, Becker J, Nikoloski Z, Bhattacharya D, Price D, Tohge T, Bar-Even A, Fernie A, et al.2019. Kingdom-wide comparison reveals the evolution of diurnal gene expression in Archaeplastida. Nature Communications 10: 737.

54. Figueiredo DD, Köhler C. 2018. Auxin: a molecular trigger of seed development. Genes & Development 32: 479–490.

55. Filonova LH, Von Arnold S, Daniel G, Bozhkov PV. 2002. Programmed cell death eliminates all but one embryo in a polyembryonic plant seed. Cell Death & Differentiation 9: 1057–1062.

56. Floyd SK, Friedman WE. 2000. Evolution of Endosperm Developmental Patterns among Basal Flowering Plants. International Journal of Plant Sciences 161: S57–S81.

57. Foroozani M, Holder DH, Deal RB. 2022. Histone Variants in the Specialization of Plant Chromatin. Annual Review of Plant Biology 73: 149–172.

58. Friedman WE. 1998. The evolution of double fertilization and endosperm: an “historical” perspective. Sexual Plant Reproduction 11: 6–16.

59. Friedman WE, Bachelier JB. 2013. Seed development in *Trimenia* (Trimeniaceae) and its bearing on the evolution of embryo-nourishing strategies in early flowering plant lineages. American Journal of Botany 100: 906–915.

60. Gao P, Quilichini TD, Yang H, Li Q, Nilsen KT, Qin L, Babic V, Liu L, Cram D, Pasha A, et al. 2021. Evolutionary divergence in embryo and seed coat development of U’s Triangle Brassica species illustrated by a spatiotemporal transcriptome atlas. New Phytologist.

61. García de la Torre VS, Majorel-Loulergue C, Rigaill GJ, Alfonso-González D, Soubigou-Taconnat L, Pillon Y, Barreau L, Thomine S, Fogliani B, Burtet-Sarramegna V, et al.2021. Wide cross-species RNA-Seq comparison reveals convergent molecular mechanisms involved in nickel hyperaccumulation across dicotyledons. New Phytologist 229: 994–1006.

62. Geeta R. 2003. The origin and maintenance of nuclear endosperms: viewing development through a phylogenetic lens. Proceedings of the Royal Society of London. Series B: Biological Sciences 270: 29–35.

63. Goodrich J, Puangsomlee P, Martin M, Long D, Meyerowitz EM, Coupland G. 1997. A Polycomb-group gene regulates homeotic gene expression in Arabidopsis. Nature 386: 44–51.

64. Groß-Hardt R, Lenhard M, Laux T. 2002. *WUSCHEL* signaling functions in interregional communication during *Arabidopsis* ovule development. Genes & Development 16: 1129–1138.

65. Guignard L. 1899. Sur les anthérozoïdes et la double copulation sexuelle chez les végétaux angiospermes. Gauthier-Villars.

66. Guo L, Yu Y, Law JA, Zhang X. 2010. SET DOMAIN GROUP2 is the major histone H3 lysine 4 trimethyltransferase in *Arabidopsis*. Proceedings of the National Academy of Sciences 107: 18557–18562.

67. Haag JR, Brower-Toland B, Krieger EK, Sidorenko L, Nicora CD, Norbeck AD, Irsigler A, LaRue H, Brzeski J, McGinnis K, et al.2014. Functional Diversification of Maize RNA Polymerase IV and V Subtypes via Alternative Catalytic Subunits. Cell Reports 9: 378–390.

68. Haig D. 1992. Brood Reduction In Gymnosperms. In: Elgar MA, Crespi BJ, eds. Cannibalism. Oxford University PressOxford, 63–84.

69. Haig D, Westoby M. 1989. Selective forces in the emergence of the seed habit. Biological Journal of the Linnean Society 38: 215–238.

70. Hansen BO, Vaid N, Musialak-Lange M, Janowski M, Mutwil M. 2014. Elucidating gene function and function evolution through comparison of co-expression networks of plants. Frontiers in Plant Science 5.

71. Hardev S. 1978. Embryology of gymnosperms. *Berlin*: Gerbrüder Borntraeger.

72. Hasegawa R, Fujita K, Tanaka Y, Takasaki H, Ikeda M, Yamagami A, Mitsuda N, Nakano T, Ohme-Takagi M. 2022. Arabidopsis zinc finger homeodomain transcription factor BRASSINOSTEROID-RELATED HOMEOBOX 2 acts as a positive regulator of brassinosteroid response. Plant Biotechnology 39: 185–189.

73. Haslekås C, Stacy RAP, Nygaard V, Culiáñez-Macià FA, Aalen RB. 1998. The expression of a peroxiredoxin antioxidant gene, AtPer1, in Arabidopsis thaliana is seed-specific and related to dormancy. Plant Molecular Biology 36: 833–845.

74. Hehenberger E, Kradolfer D, Köhler C. 2012. Endosperm cellularization defines an important developmental transition for embryo development. Development 139: 2031– 2039.

75. Henderson IR, Zhang X, Lu C, Johnson L, Meyers BC, Green PJ, Jacobsen SE. 2006. Dissecting Arabidopsis thaliana DICER function in small RNA processing, gene silencing and DNA methylation patterning. Nature Genetics 38: 721–725.

76. Hernandez Sanchez-Rebato M, Bouatta AM, Gallego ME, White CI, Da Ines O. 2021. RAD54 is essential for RAD51-mediated repair of meiotic DSB in Arabidopsis (M Grelon, Ed.). PLOS Genetics 17: e1008919.

77. Herzog M, Dorne A-M, Grellet F. 1995. GASA, a gibberellin-regulated gene family from Arabidopsis thaliana related to the tomato GAST1 gene. Plant Molecular Biology 27: 743–752.

78. Hess N, Klode M, Anders M, Sauter M. 2011. The hypoxia responsive transcription factor genes *ERF71/HRE2* and *ERF73/HRE1* of *Arabidopsis* are differentially regulated by ethylene. Physiologia Plantarum 143: 41–49.

79. Heyman J, Daele HV den, Wit KD, Boudolf V, Berckmans B, Verkest A, Kamei CLA, Jaeger GD, Koncz C, Veylder LD. 2011. Arabidopsis ULTRAVIOLET-B-INSENSITIVE4 Maintains Cell Division Activity by Temporal Inhibition of the Anaphase-Promoting Complex/Cyclosome. Plant Cell 23: 4394–4410.

80. Hong SW, Jon JH, Kwak JM, Nam HG. 1997. Identification of a Receptor-Like Protein Kinase Gene Rapidly Induced by Abscisic Acid, Dehydration, High Salt, and Cold Treatments in Arabidopsis thaliana. Plant Physiology 113: 1203–1212.

81. Hu S, Yang H, Gao H, Yan J, Xie D. 2021. Control of seed size by jasmonate. Science China Life Sciences 64: 1215–1226.

82. Huss JC, Gierlinger N. 2021. Functional packaging of seeds. New Phytologist 230: 2154–2163.

83. Immink RG, Tonaco IA, De Folter S, Shchennikova A, Van Dijk AD, Busscher-Lange J, Borst JW, Angenent GC. 2009. SEPALLATA3: the ‘glue’ for MADS box transcription factor complex formation. Genome Biology 10: R24.

84. Jarończyk K, Sosnowska K, Zaborowski A, Pupel P, Bucholc M, Małecka E, Siwirykow N, Stachula P, Iwanicka-Nowicka R, Koblowska M, et al.2021. Bromodomain-containing subunits BRD1, BRD2, and BRD13 are required for proper functioning of SWI/SNF complexes in Arabidopsis. Plant Communications 2: 100174.

85. Jiang H, Moreno-Romero J, Santos-González J, De Jaeger G, Gevaert K, Van De Slijke E, Köhler C. 2017. Ectopic application of the repressive histone modification H3K9me2 establishes post-zygotic reproductive isolation in Arabidopsis thaliana. Genes & Development 31: 1272–1287.

86. Julca I, Ferrari C, Flores-Tornero M, Proost S, Lindner A-C, Hackenberg D, Steinbachová L, Michaelidis C, Gomes Pereira S, Misra CS, et al.2021. Comparative transcriptomic analysis reveals conserved programmes underpinning organogenesis and reproduction in land plants. Nature Plants 7: 1143–1159.

87. Julca I, Tan QW, Mutwil M. 2023. Toward kingdom-wide analyses of gene expression. Trends in Plant Science 28: 235–249.

88. Kang J, Park J, Choi H, Burla B, Kretzschmar T, Lee Y, Martinoia E. 2011. Plant ABC Transporters. The Arabidopsis Book / American Society of Plant Biologists 9: e0153.

89. Kazda A, Zellinger B, Rössler M, Derboven E, Kusenda B, Riha K. 2012. Chromosome end protection by blunt-ended telomeres. Genes & Development 26: 1703–1713.

90. Kim SY, He Y, Jacob Y, Noh Y-S, Michaels S, Amasino R. 2005. Establishment of the Vernalization-Responsive, Winter-Annual Habit in *Arabidopsis* Requires a Putative Histone H3 Methyl Transferase. The Plant Cell 17: 3301–3310.

91. Kim D, Langmead B, Salzberg SL. 2015. HISAT: A fast spliced aligner with low memory requirements. Nature Methods 12: 357–360.

92. Kim YJ, Wang R, Gao L, Li D, Xu C, Mang H, Jeon J, Chen X, Zhong X, Kwak JM, et al. 2016. POWERDRESS and HDA9 interact and promote histone H3 deacetylation at specific genomic sites in *Arabidopsis*. Proceedings of the National Academy of Sciences 113: 14858–14863.

93. Knutson BA. 2013. Emergence and expansion of TFIIB-like factors in the plant kingdom. Gene 526: 30–38.

94. Köhler C, Dziasek K, Del Toro-De León G. 2021. Postzygotic reproductive isolation established in the endosperm: mechanisms, drivers and relevance. Philosophical Transactions of the Royal Society B: Biological Sciences 376: 20200118.

95. Kondou Y, Nakazawa M, Kawashima M, Ichikawa T, Yoshizumi T, Suzuki K, Ishikawa A, Koshi T, Matsui R, Muto S, et al. 2008. RETARDED GROWTH OF EMBRYO1, a New Basic Helix-Loop-Helix Protein, Expresses in Endosperm to Control Embryo Growth. Plant Physiology 147: 1924–1935.

96. Koramutla MK, Negi M, Ayele BT. 2021. Roles of Glutathione in Mediating Abscisic Acid Signaling and Its Regulation of Seed Dormancy and Drought Tolerance. Genes 12: 1620.

97. Kraft E, Stone SL, Ma L, Su N, Gao Y, Lau O-S, Deng X-W, Callis J. 2005. Genome Analysis and Functional Characterization of the E2 and RING-Type E3 Ligase Ubiquitination Enzymes of Arabidopsis. Plant Physiology 139: 1597–1611.

98. Kwong RW, Bui AQ, Lee H, Kwong LW, Fischer RL, Goldberg RB, Harada JJ. 2003. LEAFY COTYLEDON1-LIKE Defines a Class of Regulators Essential for Embryo Development. The Plant Cell 15: 5–18.

99. Lafon-Placette C, Köhler C. 2014. Embryo and endosperm, partners in seed development. Current Opinion in Plant Biology 17: 64–69.

100. Lara P, Oñate-Sánchez L, Abraham Z, Ferrándiz C, Díaz I, Carbonero P, Vicente-Carbajosa J. 2003. Synergistic activation of seed storage protein gene expression in Arabidopsis by ABI3 and two bZIPs related to OPAQUE2. The Journal of Biological Chemistry 278: 21003–21011.

101. Larkin RM, Hagen G, Guilfoyle TJ. 1999. Arabidopsis thaliana RNA polymerase II subunits related to yeast and human RPB5. Gene 231: 41–47.

102. Le BH, Cheng C, Bui AQ, Wagmaister JA, Henry KF, Pelletier J, Kwong L, Belmonte M, Kirkbride R, Horvath S, et al. 2010. Global analysis of gene activity during *Arabidopsis* seed development and identification of seed-specific transcription factors. Proceedings of the National Academy of Sciences 107: 8063–8070.

103. Leebens-Mack JH, Barker MS, Carpenter EJ, Deyholos MK, Gitzendanner MA, Graham SW, Grosse I, Li Z, Melkonian M, Mirarab S, et al.2019. One thousand plant transcriptomes and the phylogenomics of green plants. Nature 574: 679–685.

104. Li S, Zachgo S. 2013. TCP3 interacts with R2R3-MYB proteins, promotes flavonoid biosynthesis and negatively regulates the auxin response in Arabidopsis thaliana. The Plant Journal 76: 901–913.

105. Lin Y, Pajak A, Marsolais F, McCourt P, Riggs CD. 2013. Characterization of a Cruciferin Deficient Mutant of Arabidopsis and Its Utility for Overexpression of Foreign Proteins in Plants. PLoS ONE 8: e64980.

106. Lin IW, Sosso D, Chen L-Q, Gase K, Kim S-G, Kessler D, Klinkenberg PM, Gorder MK, Hou B-H, Qu X-Q, et al.2014. Nectar secretion requires sucrose phosphate synthases and the sugar transporter SWEET9. Nature 508: 546–549.

107. Linkies A, Graeber K, Knight C, Leubner-Metzger G. 2010. The evolution of seeds. New Phytologist 186: 817–831.

108. Linkies A, Leubner-Metzger G. 2012. Beyond gibberellins and abscisic acid: how ethylene and jasmonates control seed germination. Plant Cell Reports 31: 253–270.

109. Liu J, Wu M-W, Liu C-M. 2022. Cereal Endosperms: Development and Storage Product Accumulation. Annual Review of Plant Biology 73: 255–291.

110. Love MI, Huber W, Anders S. 2014. Moderated estimation of fold change and dispersion for RNA-seq data with DESeq2. Genome Biology 15: 550.

111. Lu J, Zhang C, Baulcombe D, Chen Z. 2012. Maternal siRNAs as regulators of parental genome imbalance and gene expression in endosperm of *Arabidopsis* seeds. Proceedings of the National Academy of Sciences of the United States of America 109: 5529–5534.

112. Lubna, Asaf S, Khan AL, Jan R, Khan A, Khan A, Kim K, Lee I. 2021. The dynamic history of gymnosperm plastomes: Insights from structural characterization, comparative analysis, phylogenomics, and time divergence. The Plant Genome 14: e20130.

113. Luo M, Bilodeau P, Dennis ES, Peacock WJ, Chaudhury A. 2000. Expression and parent-of-origin effects for FIS2, MEA, and FIE in the endosperm and embryo of developing Arabidopsis seeds. Proceedings of the National Academy of Sciences 97: 10637–10642.

114. Martinez G, Wolff P, Wang Z, Moreno-Romero J, Santos-González J, Conze LL, DeFraia C, Slotkin RK, Köhler C. 2018. Paternal easiRNAs regulate parental genome dosage in Arabidopsis. Nat Genet 50: 193–198.

115. Masucci JD, Rerie WG, Foreman DR, Zhang M, Galway ME, Marks MD, Schiefelbein JW. 1996. The homeobox gene *GLABRA 2* is required for position-dependent cell differentiation in the root epidermis of *Arabidopsis thaliana*. Development 122: 1253–1260.

116. Meesapyodsuk D, Qiu X. 2011. A Peroxygenase Pathway Involved in the Biosynthesis of Epoxy Fatty Acids in Oat. Plant Physiology 157: 454–463.

117. Mei S, Zhang M, Ye J, Du J, Jiang Y, Hu Y. 2023. Auxin contributes to jasmonate-mediated regulation of abscisic acid signaling during seed germination in Arabidopsis. The Plant Cell 35: 1110–1133.

118. Millar AA, Gubler F. 2005. The Arabidopsis *GAMYB-Like* Genes, *MYB33* and *MYB65*, Are MicroRNA-Regulated Genes That Redundantly Facilitate Anther Development. The Plant Cell 17: 705–721.

119. Mintz PJ, Patterson SD, Neuwald AF, Spahr CS, Spector DL. 1999. Purification and biochemical characterization of interchromatin granule clusters. The EMBO Journal 18: 4308–4320.

120. Mitsuda H, Yasumoto K, Murakami K, Kusano T, Kishida H. 1967. Studies on the Proteinaceous Subcellular Particles in Rice Endosperm: Electron-Microscopy and Isolation. Agricultural and Biological Chemistry 31: 293–300.

121. Molaro A, Wood AJ, Janssens D, Kindelay SM, Eickbush MT, Wu S, Singh P, Muller CH, Henikoff S, Malik HS. 2020. Biparental contributions of the H2A.B histone variant control embryonic development in mice. PLoS Biology 18: e3001001.

122. Mönke G, Seifert M, Keilwagen J, Mohr M, Grosse I, Hähnel U, Junker A, Weisshaar B, Conrad U, Bäumlein H, et al. 2012. Toward the identification and regulation of the Arabidopsis thaliana ABI3 regulon. Nucleic Acids Research 40: 8240– 8254.

123. Montgomery SA, Berger F. 2021. The evolution of imprinting in plants: beyond the seed. Plant Reproduction 34: 373–383.

124. Mosher RA, Melnyk CW, Kelly KA, Dunn RM, Studholme DJ, Baulcombe DC. 2009. Uniparental expression of PolIV-dependent siRNAs in developing endosperm of Arabidopsis. Nature 460: 283–286.

125. Moussu S, Doll NM, Chamot S, Brocard L, Creff A, Fourquin C, Widiez T, Nimchuk ZL, Ingram G. 2017. ZHOUPI and KERBEROS Mediate Embryo/Endosperm Separation by Promoting the Formation of an Extracuticular Sheath at the Embryo Surface. The Plant Cell 29: 1642–1656.

126. Müller M, Munné-Bosch S. 2015. Ethylene Response Factors: A Key Regulatory Hub in Hormone and Stress Signaling. Plant Physiology 169: 32–41.

127. Müller D, Schmitz G, Theres K. 2006. *Blind* Homologous *R2R3 Myb* Genes Control the Pattern of Lateral Meristem Initiation in *Arabidopsis*. The Plant Cell 18: 586–597.

128. Nawaschin S. 1898. Resultate einer Revision der Befruchtungsvorgange bei Lilium martagon und Fritillaria tenella. Известия Российской академии наук. Серия математическая 9: 377–382.

129. Nemhauser JL. 2018. Back to basics: what is the function of an Aux/ IAA in auxin response? New Phytologist 218: 1295–1297.

130. Nietzel T, Dudkina NV, Haase C, Denolf P, Semchonok DA, Boekema EJ, Braun H-P, Sunderhaus S. 2013. The Native Structure and Composition of the Cruciferin Complex in Brassica napus. Journal of Biological Chemistry 288: 2238–2245.

131. Noh Y-S, Amasino RM. 2003. *PIE1* , an ISWI Family Gene, Is Required for *FLC* Activation and Floral Repression in Arabidopsis. The Plant Cell 15: 1671–1682.

132. Norstog K. 1982. Experimental Embryology of Gymnosperms. In: Johri BM, ed. Experimental Embryology of Vascular Plants. Berlin, Heidelberg: Springer Berlin Heidelberg, 25–51.

133. Okushima Y, Mitina I, Quach HL, Theologis A. 2005. AUXIN RESPONSE FACTOR 2 (ARF2): a pleiotropic developmental regulator. The Plant Journal 43: 29– 46.

134. O’Leary BM. 2020. The Lure of Lignin: Deciphering High-value Lignin Formation in Seed Coats. The Plant Cell 32: 3652–3653.

135. Olsen O-A. 2004. Nuclear Endosperm Development in Cereals and Arabidopsis thaliana. THE PLANT CELL ONLINE 16: S214–S227.

136. Oneal E, Willis JH, Franks RG. 2016. Disruption of endosperm development is a major cause of hybrid seed inviability between *Mimulus guttatus* and *Mimulus nudatus*. New Phytologist 210: 1107–1120.

137. Osakabe K, Abe K, Yoshioka T, Osakabe Y, Todoriki S, Ichikawa H, Hohn B, Toki S. 2006. Isolation and characterization of the *RAD54* gene from *Arabidopsis thaliana*. The Plant Journal 48: 827–842.

138. Pan J, Hu Y, Wang H, Guo Q, Chen Y, Howe GA, Yu D. 2020a. Molecular Mechanism Underlying the Synergetic Effect of Jasmonate on Abscisic Acid Signaling during Seed Germination in Arabidopsis. The Plant Cell 32: 3846–3865.

139. Pan R, Liu J, Wang S, Hu J. 2020b. Peroxisomes: versatile organelles with diverse roles in plants. New Phytologist 225: 1410–1427.

140. Pandey R. 2002. Analysis of histone acetyltransferase and histone deacetylase families of Arabidopsis thaliana suggests functional diversification of chromatin modification among multicellular eukaryotes. Nucleic Acids Research 30: 5036–5055.

141. Pang PP, Pruitt RE, Meyerowitz EM. 1988. Molecular cloning, genomic organization, expression and evolution of 12S seed storage protein genes of Arabidopsis thaliana. Plant Molecular Biology 11: 805–820.

142. Park M, Krause C, Karnahl M, Reichardt I, El Kasmi F, Mayer U, Stierhof Y-D, Hiller U, Strompen G, Bayer M, et al. 2018. Concerted Action of Evolutionarily Ancient and Novel SNARE Complexes in Flowering-Plant Cytokinesis. Developmental Cell 44: 500–511.e4.

143. Payne PI, Holt LM, Lawrence GJ, Law CN. 1982. The genetics of gliadin and glutenin, the major storage proteins of the wheat endosperm. Qualitas Plantarum Plant Foods for Human Nutrition 31: 229–241.

144. Pazhouhandeh M, Molinier J, Berr A, Genschik P. 2011. MSI4/FVE interacts with CUL4–DDB1 and a PRC2-like complex to control epigenetic regulation of flowering time in Arabidopsis. Proceedings of the National Academy of Sciences 108: 3430–3435.

145. Peer WA, Murphy AS. 2007. Flavonoids and auxin transport: modulators or regulators? Trends in Plant Science 12: 556–563.

146. Peng J, Carol P, Richards DE, King KE, Cowling RJ, Murphy GP, Harberd NP. 1997. The *Arabidopsis GAI* gene defines a signaling pathway that negatively regulates gibberellin responses. Genes & Development 11: 3194–3205.

147. Pettitt J. 1970. HETEROSPORY AND THE ORIGIN OF THE SEED HABIT. Biological Reviews 45: 401–415.

148. Pfannschmidt T, Blanvillain R, Merendino L, Courtois F, Chevalier F, Liebers M, Grübler B, Hommel E, Lerbs-Mache S. 2015. Plastid RNA polymerases: orchestration of enzymes with different evolutionary origins controls chloroplast biogenesis during the plant life cycle. Journal of Experimental Botany 66: 6957–6973.

149. Porta H, Rocha-Sosa M. 2002. Plant Lipoxygenases. Physiological and Molecular Features. Plant Physiology 130: 15–21.

150. Povilus RA, Diggle PK, Friedman WE. 2018. Evidence for parent-of-origin effects and interparental conflict in seeds of an ancient flowering plant lineage. Proceedings of the Royal Society B: Biological Sciences 285: 20172491.

151. Povilus RA, Friedman WE. 2022. Transcriptomes across fertilization and seed development in the water lily Nymphaea thermarum (Nymphaeales): evidence for epigenetic patterning during reproduction. Plant Reproduction.

152. Povilus RA, Losada JM, Friedman WE. 2015. Floral biology and ovule and seed ontogeny of Nymphaea thermarum, a water lily at the brink of extinction with potential as a model system for basal angiosperms. Annals of Botany 115: 211–226.

153. Poxleitner M, Rogers SW, Lacey Samuels A, Browse J, Rogers JC. 2006. A role for caleosin in degradation of oil-body storage lipid during seed germination. The Plant Journal 47: 917–933.

154. Prigge MJ, Otsuga D, Alonso JM, Ecker JR, Drews GN, Clark SE. 2005. Class III Homeodomain-Leucine Zipper Gene Family Members Have Overlapping, Antagonistic, and Distinct Roles in Arabidopsis Development. The Plant Cell 17: 61–76.

155. Qiao Z, Pingault L, Nourbakhsh-Rey M, Libault M. 2016. Comprehensive Comparative Genomic and Transcriptomic Analyses of the Legume Genes Controlling the Nodulation Process. Frontiers in Plant Science 7: 34.

156. Ran J-H, Shen T-T, Wang M-M, Wang X-Q. 2018. Phylogenomics resolves the deep phylogeny of seed plants and indicates partial convergent or homoplastic evolution between Gnetales and angiosperms. Proceedings of the Royal Society B: Biological Sciences 285: 20181012.

157. Rangan P. 2020. Endosperm variability: from endoreduplication within a seed to higher ploidy across species, and its competence. Seed Science Research 30: 173–185.

158. Rautengarten C, Usadel B, Neumetzler L, Hartmann J, Büssis D, Altmann T. 2008. A subtilisin-like serine protease essential for mucilage release from Arabidopsis seed coats. The Plant Journal 54: 466–480.

159. Ream TS, Haag JR, Wierzbicki AT, Nicora CD, Norbeck AD, Zhu J-K, Hagen G, Guilfoyle TJ, Paša-Tolić L, Pikaard CS. 2009. Subunit Compositions of the RNA-Silencing Enzymes Pol IV and Pol V Reveal Their Origins as Specialized Forms of RNA Polymerase II. Molecular Cell 33: 192–203.

160. Rodrigues AS, Chaves I, Costa BV, Lin Y-C, Lopes S, Milhinhos A, Van De Peer Y, Miguel CM. 2019. Small RNA profiling in Pinus pinaster reveals the transcriptome of developing seeds and highlights differences between zygotic and somatic embryos. Scientific Reports 9: 11327.

161. Rodrigues JA, Ruan R, Nishimura T, Sharma MK, Sharma R, Ronald PC, Fischer RL, Zilberman D. 2013. Imprinted expression of genes and small RNA is associated with localized hypomethylation of the maternal genome in rice endosperm. Proceedings of the National Academy of Sciences 110: 7934–7939.

162. Rodrigues AS, Vega JJ, Miguel CM. 2018. Comprehensive assembly and analysis of the transcriptome of maritime pine developing embryos. BMC Plant Biology 18: 1–20.

163. Roth M, Florez-Rueda AM, Griesser S, Paris M, Städler T. 2018. Incidence and developmental timing of endosperm failure in post-zygotic isolation between wild tomato lineages. Annals of Botany 121: 107–118.

164. Sabelli PA, Larkins BA. 2009. The development of endosperm in grasses. Plant Physiology 149: 14–26.

165. Sakai S. 2013. EVOLUTIONARILY STABLE SIZE OF A MEGAGAMETOPHYTE: EVOLUTION OF TINY MEGAGAMETOPHYTES OF ANGIOSPERMS FROM LARGE ONES OF GYMNOSPERMS: SIZE OF MEGAGAMETOPHYTES. Evolution 67: 539–547.

166. Schmidt R, Stransky H, Koch W. 2007. The amino acid permease AAP8 is important for early seed development in Arabidopsis thaliana. Planta 226: 805–813.

167. Serrano M, Parra S, Alcaraz LD, Guzmán P. 2006. The ATL Gene Family from Arabidopsis thaliana and Oryza sativa Comprises a Large Number of Putative Ubiquitin Ligases of the RING-H2 Type. Journal of Molecular Evolution 62: 434–445.

168. Shewry PR, Halford NG. 2002. Cereal seed storage proteins: structures, properties and role in grain utilization. Journal of Experimental Botany 53: 947–958.

169. Shewry PR, Napier JA, Tatham AS. 1995. Seed storage proteins: structures and biosynthesis. The Plant Cell 7: 945–956.

170. Shultz RW, Lee T-J, Allen GC, Thompson WF, Hanley-Bowdoin L. 2009. Dynamic Localization of the DNA Replication Proteins MCM5 and MCM7 in Plants. Plant Physiology 150: 658–669.

171. Shutov AD, Kakhovskaya IA, Braun H, Bäumlein H, Mäntz K. 1995. Legumin-like and vicilin-like seed storage proteins: Evidence for a common single-domain ancestral gene. Journal of Molecular Evolution 41.

172. Singh H, Johri BM. 1972. DEVELOPMENT OF GYMNOSPERM SEEDS. In: Seed Biology. Elsevier, 21–75.

173. Somers DE, Schultz TF, Milnamow M, Kay SA. 2000. ZEITLUPE Encodes a Novel Clock-Associated PAS Protein from Arabidopsis. Cell 101: 319–329.

174. Song J, Xie X, Chen C, Shu J, Thapa RK, Nguyen V, Bian S, Kohalmi SE, Marsolais F, Zou J, et al. 2021. LEAFY COTYLEDON1 expression in the endosperm enables embryo maturation in Arabidopsis. Nature Communications 12.

175. Souto LS, Oliveira DMT. 2014. Seed development in Malpighiaceae species with an emphasis on the relationships between nutritive tissues. Comptes Rendus Biologies 337: 62–70.

176. Spencer ART, Wang S-J, Dunn MT, Hilton J. 2013. Species of the medullosan ovule Stephanospermum from the Lopingian (late Permian) floras of China. Journal of Asian Earth Sciences 76: 59–69.

177. Sreenivasulu N, Borisjuk L, Junker BH, Mock H-P, Rolletschek H, Seiffert U, Weschke W, Wobus U. 2010. Barley Grain Development. In: International Review of Cell and Molecular Biology. Elsevier, 49–89.

178. Sreenivasulu N, Wobus U. 2013. Seed-Development Programs: A Systems Biology– Based Comparison Between Dicots and Monocots. Annual Review of Plant Biology 64: 189–217.

179. Stracke R, Werber M, Weisshaar B. 2001. The R2R3-MYB gene family in Arabidopsis thaliana. Current Opinion in Plant Biology 4: 447–456.

180. Swamy BGL, Parameswaran N. 1963. THE HELOBIAL ENDOSPERM. Biological Reviews 38: 1–50.

181. Szklarczyk D, Morris JH, Cook H, Kuhn M, Wyder S, Simonovic M, Santos A, Doncheva NT, Roth A, Bork P, et al. 2017. The STRING database in 2017: Quality-controlled protein-protein association networks, made broadly accessible. Nucleic Acids Research 45: D362–D368.

182. Takahashi S, Sakamoto AN, Tanaka A, Shimizu K. 2007. AtREV1, a Y-Family DNA Polymerase in Arabidopsis, Has Deoxynucleotidyl Transferase Activity in Vitro. Plant Physiology 145: 1052–1060.

183. The Angiosperm Phylogeny Group. 2016. An update of the Angiosperm Phylogeny Group classification for the orders and families of flowering plants: APG IV. Botanical Journal of the Linnean Society 181: 1–20.

184. Thines B, Katsir L, Melotto M, Niu Y, Mandaokar A, Liu G, Nomura K, He SY, Howe GA, Browse J. 2007. JAZ repressor proteins are targets of the SCFCOI1 complex during jasmonate signalling. Nature 448: 661–665.

185. Tiwari S, Spielman M, Schulz R, Oakey RJ, Kelsey G, Salazar A, Zhang K, Pennell R, Scott RJ. 2010. Transcriptional profiles underlying parent-of-origin effects in seeds of Arabidopsis thaliana. BMC Plant Biology 10: 72.

186. Tobe H, Jaffré T, Raven PH. 2000. Embryology of Amborella (Amborellaceae): Descriptions and Polarity of Character States. Journal of Plant Research 113: 271–280.

187. Tobimatsu Y, Chen F, Nakashima J, Escamilla-Trevino LL, Jackson L, Dixon RA, Ralph J. 2013. Coexistence but Independent Biosynthesis of Catechyl and Guaiacyl/Syringyl Lignin Polymers in Seed Coats. The Plant Cell 25: 2587–2600.

188. Tsuwamoto R, Fukuoka H, Takahata Y. 2008. *GASSHO1* and *GASSHO2* encoding a putative leucine-rich repeat transmembrane-type receptor kinase are essential for the normal development of the epidermal surface in Arabidopsis embryos. The Plant Journal 54: 30–42.

189. Ungru A, Nowack MK, Reymond M, Shirzadi R, Kumar M, Biewers S, Grini PE, Schnittger A. 2008. Natural variation in the degree of autonomous endosperm formation reveals independence and constraints of embryo growth during seed development in Arabidopsis thaliana. Genetics 179: 829–841.

190. Van Der Graaff E, Laux T, Rensing SA. 2009. The WUS homeobox-containing (WOX) protein family. Genome Biology 10: 248.

191. Vandelook F, Newton RJ, Bobon N, Bohley K, Kadereit G. 2021. Evolution and ecology of seed internal morphology in relation to germination characteristics in Amaranthaceae. Annals of Botany 127: 799–811.

192. Vanstraelen M, Inzé D, Geelen D. 2006. Mitosis-specific kinesins in Arabidopsis. Trends in Plant Science 11: 167–175.

193. Vazquez F, Gasciolli V, Crété P, Vaucheret H. 2004. The Nuclear dsRNA Binding Protein HYL1 Is Required for MicroRNA Accumulation and Plant Development, but Not Posttranscriptional Transgene Silencing. Current Biology 14: 346–351.

194. Vercruysse J, Van Bel M, Osuna-Cruz CM, Kulkarni SR, Storme V, Nelissen H, Gonzalez N, Inzé D, Vandepoele K. 2020. Comparative transcriptomics enables the identification of functional orthologous genes involved in early leaf growth. Plant Biotechnology Journal 18: 553–567.

195. Vergara Z, Gomez MS, Desvoyes B, Sequeira-Mendes J, Masoud K, Costas C, Noir S, Caro E, Mora-Gil V, Genschik P, et al. 2023. Distinct roles of Arabidopsis ORC1 proteins in DNA replication and heterochromatic H3K27me1 deposition. Nature Communications 14: 1270.

196. Vick BA, Zimmerman DC. 1983. The biosynthesis of jasmonic acid: A physiological role for plant lipoxygenase. Biochemical and Biophysical Research Communications 111: 470–477.

197. Vogt T. 2010. Phenylpropanoid Biosynthesis. Molecular Plant 3: 2–20.

198. Wagner U, Edwards R, Dixon DP, Mauch F. 2002. Probing the diversity of the Arabidopsis glutathione S-transferase gene family. Plant Molecular Biology 49: 515– 532.

199. Wasternack C, Feussner I. 2018. The Oxylipin Pathways: Biochemistry and Function. Annual Review of Plant Biology 69: 363–386.

200. Wasternack C, Strnad M. 2018. Jasmonates: News on Occurrence, Biosynthesis, Metabolism and Action of an Ancient Group of Signaling Compounds. International Journal of Molecular Sciences 19: 2539.

201. Weber H, Borisjuk L, Wobus U. 2005. MOLECULAR PHYSIOLOGY OF LEGUME SEED DEVELOPMENT. Annual Review of Plant Biology 56: 253–279.

202. Williams CG (Ed.). 2009. Syngamy, Embryo Development and Seed Dispersal. In: Conifer Reproductive Biology. Dordrecht: Springer Netherlands, 107–121.

203. Williams JH, Friedman WE. 2002. Identification of diploid endosperm in an early angiosperm lineage. Nature 415: 522–526.

204. Willis CG, Baskin CC, Baskin JM, Auld JR, Venable DL, Cavender-Bares J, Donohue K, Rubio De Casas R, The NESCent Germination Working Group. 2014. The evolution of seed dormancy: environmental cues, evolutionary hubs, and diversification of the seed plants. New Phytologist 203: 300–309.

205. Xin M, Yang R, Yao Y, Ma C, Peng H, Sun Q, Wang X, Ni Z. 2014. Dynamic parent-of-origin effects on small interfering RNA expression in the developing maize endosperm. BMC Plant Biology 14: 192.

206. Xing Q, Creff A, Waters A, Tanaka H, Goodrich J, Ingram GC. 2013. ZHOUPI controls embryonic cuticle formation via a signalling pathway involving the subtilisin protease ABNORMAL LEAF-SHAPE1 and the receptor kinases GASSHO1 and GASSHO2. *Development (Cambridge*, England) 140: 770–779.

207. Xu W, Fiume E, Coen O, Pechoux C, Lepiniec L, Magnani E. 2016. Endosperm and Nucellus Develop Antagonistically in Arabidopsis Seeds. The Plant Cell: tpc.00041.2016.

208. Yang T, Guo L, Ji C, Wang H, Wang J, Zheng X, Xiao Q, Wu Y. 2020. The B3 domain-containing transcription factor ZmABI19 coordinates expression of key factors required for maize seed development and grain filling. The Plant Cell 33: 104–128.

209. Yang S, Johnston N, Talideh E, Mitchell S, Jeffree C, Goodrich J, Ingram G. 2008. The endosperm-specific *ZHOUPI* gene of *Arabidopsis thaliana* regulates endosperm breakdown and embryonic epidermal development. Development 135: 3501–3509.

210. Yang H, Xiang D, Venglat SP, Cao Y, Wang E, Selvaraj G, Datla R. 2009. *PolA2* is required for embryo development in *Arabidopsis* This paper is one of a selection of papers published in a Special Issue from the National Research Council of Canada – Plant Biotechnology Institute. Botany 87: 626–634.

211. Yu Y, Hu H, Doust AN, Kellogg EA. 2020. Divergent gene expression networks underlie morphological diversity of abscission zones in grasses. The New Phytologist 225: 1799–1815.

212. Zahn LM, Kong H, Leebens-Mack JH, Kim S, Soltis PS, Landherr LL, Soltis DE, dePamphilis CW, Ma H. 2005. The Evolution of the SEPALLATA Subfamily of MADS-Box Genes. Genetics 169: 2209–2223.

213. Zhang K, Novak O, Wei Z, Gou M, Zhang X, Yu Y, Yang H, Cai Y, Strnad M, Liu C-J. 2014. Arabidopsis ABCG14 protein controls the acropetal translocation of root-synthesized cytokinins. Nature Communications 5: 3274.

214. Zhang Y-J, Wang W, Yang H-L, Li Y, Kang X-Y, Wang X-R, Yang Z-L. 2015. Molecular Properties and Functional Divergence of the Dehydroascorbate Reductase Gene Family in Lower and Higher Plants (J-S Zhang, Ed.). PLOS ONE 10: e0145038.

215. Zhao D, Ni W, Feng B, Han T, Petrasek MG, Ma H. 2003. Members of the *Arabidopsis-SKP1-like* Gene Family Exhibit a Variety of Expression Patterns and May Play Diverse Roles in Arabidopsis. Plant Physiology 133: 203–217.

216. Zheng Y, Wang Z. 2015. The cereal starch endosperm development and its relationship with other endosperm tissues and embryo. Protoplasma 252: 33–40.

217. Zhou M, Law JA. 2015. RNA Pol IV and V in gene silencing: Rebel polymerases evolving away from Pol II’s rules. Current Opinion in Plant Biology 27: 154–164.

218. Zhou A, Wang H, Walker JC, Li J. 2004. BRL1, a leucine-rich repeat receptor-like protein kinase, is functionally redundant with BRI1 in regulating *Arabidopsis* brassinosteroid signaling. The Plant Journal 40: 399–409.

219. Zimmer A, Lang D, Richardt S, Frank W, Reski R, Rensing SA. 2007. Dating the early evolution of plants: detection and molecular clock analyses of orthologs. Molecular Genetics and Genomics 278: 393–402.

220. Zumajo-Cardona C, Ambrose BA. 2021. Deciphering the evolution of the ovule genetic network through expression analyses in *Gnetum gnemon*. Annals of Botany 128: 217–230.

221. Zumajo-Cardona C, Little DP, Stevenson D, Ambrose BA. 2021. Expression analyses in Ginkgo biloba provide new insights into the evolution and development of the seed. Scientific Reports 11: 21995.

