## Supplementary methods and figures for "Comparative transcriptomics of seed nourishing tissues: uncovering conserved and divergent pathways in seed plants" (from another version of this preprint: version 1, 2023-12-01)

The following Supporting Information is available for this article:

SI Experimental Procedures
SI References
SI Tables S1 to S9
SI Figures S1 to S10

**Methods S1**

*Plant material and sample preparation*

Immature female cones from *Pinus pinaster* Ait. were collected between mid-June and end of July 2013 from open-pollinated trees in a clonal seed orchard at Mata Nacional do Escaroupim, Portugal (Longitude 8°44'W, latitude 39°4'N) upon permission by Alexandre Aguiar and Isabel Carrasquinho from INIAV (Oeiras, Portugal) complying with institutional and national guidelines. In these species pollination takes place one year before fertilization. Seeds were removed and opened to dissect megagametophytes which were categorized into stages MG0, MG3 and MG4 in accordance with the developmental stage of the respective embryos following quick evaluation under the stereomicroscope, as previously described (Rodrigues *et al.*, 2019). Megagametophytes were immediately frozen in liquid nitrogen and stored at -80ºC until further analysis. Total RNA was extracted using the MagMax Plant RNA Isolation Kit and cDNA was synthesized with the RevertAid First Strand cDNA synthesis kit using an oligo dT primer. Libraries were prepared using the Truseq RNA Illumina kit and sequenced using a Novaseq6000 machine. *A. trichopoda* seeds were collected from naturally pollinated flowers in the Bonn Botanical Garden in January of 2021 (pollination in *A. trichopoda* normally takes place around November) as the earliest time point of collection possible for this species (seed development in *A. trichopoda* takes around one year; Figure S1). *N. caerulea* seeds were collected from the flowers of the Potsdam botanical garden one week after manual pollination. The endosperms of both species were laser-captured as described by (Florez-Rueda *et al.*, 2016a). Briefly, the material was rapidly fixed, and embedded in paraffin. After sectioning and slide preparation, the tissues were laser microdissected (Figure S1). In *N. caerulea* endosperm and perisperm were dissected separately. Total RNA was extracted with the PicoPure RNA Isolation kit and cDNA was synthesized using the SMART-Seq v4 Ultra Low Input RNA Kit. The recovered tissue libraries were prepared using Nextera XT kits and sequenced using a Novaseq6000 machine. For the *N. caerulea* leaf transcriptome, RNA was extracted using the MagMax Plant RNA Isolation kit, and libraries were prepared and sequenced using a Novaseq6000 machine. Additional information for the generated datasets is in Table S1.

*Data acquisition*

For our analyses, we used publicly available data from the earliest available time points of seed development for each species. Data was gathered from the European Nucleotide Archive (ENA) as follows: for Arabidopsis endosperm, we used data from (Moreno-Romero *et al.*, 2019; Batista *et al.*, 2019) for wild-type Col-0 leaf transcriptomes, we used data from Cao *et al.*, (2020) and additional libraries from the PRJNA686693 repository. For *Solanum peruvianum*, we used the leaf and endosperm data from (Florez-Rueda *et al.*, 2016b). For *Mimulus guttatus*, we used endosperm data from (Kinser *et al.*, 2021) and leaf data from (Zuellig & Sweigart, 2018) . For rice, we used the endosperm data from (Wang *et al.*, 2018b) and (Chen *et al.*, 2018) and leaf data from the PRJNA552367 repository and (Gao *et al.*, 2021). For maize, the endosperm data were obtained from (Yang *et al.*, 2020; Wang *et al.*, 2022) and leaf data were gathered from (Wang *et al.*, 2018a) and the PRJEB48470 repository. Leaf data for *Pinus pinaster* were obtained from the

PRJNA262683 repository. *Amborella* leaf data were gathered from PRJEB38698 on ENA. All *Ginkgo* data was described in Additional information for the assembled datasets is in Table S1.

*Orthogroup inference*

To compare the transcriptional landscape of the seed-nourishing tissue across dissimilar distant seed conformations, we first identified OGs between our target species using OrthoFinder (Emms & Kelly, 2019). Reference protein genome sequences were retrieved as follows (file and web location): *Arabidopsis thaliana* (*Arabidopsis_thaliana.TAIR10.pep.all.fasta* in https://www.arabidopsis.org/download/index-auto.jsp?dir=%2Fdownload_files%2FGenes%2FAraport11_genome_release), *Solanum lycopersicon* (*S_lycopersicum_chromosomes.4.00.fa* in https://solgenomics.net/ftp/tomato_genome/annotation/ITAG4.1_release/), *Amborella trichopoda* (*Atrichopoda_291_v1.0.protein.fasta* in https://data.jgi.doe.gov/refine-download/phytozome?organism=Atrichopoda&expanded=291), *Nymphaea colorata* (*Ncolorata_566_v1.2.protein.fa* in https://data.jgi.doe.gov/refine-download/phytozome?organism=Ncolorata&expanded=566), *Pinus tadea* (*Pita.2_01.pep.fa* in https://treegenesdb.org/FTP/Genomes/Pita/v2.01/), *Mimulus guttatus* (*Mguttatus_256_v2.0.protein.fa* in https://data.jgi.doe.gov/refine-download/phytozome?organism=Mguttatus&expanded=256), *Zea mays (Zea_mays.Zm-B73-REFERENCE-NAM-5.0.pep.all.fa,* http://plants.ensembl.org/Zea_mays/Info/Index), *Oryza sativa* (*Oryza_sativa.IRGSP-1.0.pep.all.fa*, http://plants.ensembl.org/Oryza_sativa/Info/Index) and *Ginkgo biloba* (*Ginkgo_biloba.pep.all.v2.fa.gz* in *https://ginkgo.zju.edu.cn/ftp/Genome/version-2021/protein/*) for Orthogroup inference. In addition, by implementing Orthofinder, we inferred a species tree from the set of unrooted orthogroup gene trees. The species tree displays the number of gene duplications that occurred on each branch of the species tree as evidenced by multiple copies of the gene in at least one species (Emms & Kelly, 2019). Orthofinder was run with default settings.

*Transcriptomic analyses*

RNAseq datasets were downloaded or newly produced for endosperm and leaf tissue as described in Table S1. Briefly, raw reads were quality-filtered using Trimmomatic and then mapped to reference genomes corresponding to the previously referenced versions and sources using Hisat2 (Kim *et al.*, 2015). StringTie (Pertea *et al.*, 2015) was used to generate counts per gene using only high quality mapped reads. DESEQ (Love *et al.*, 2014) was used to normalize counts from different sources and allow for data integration.

In order to visualize the relationships, similarities and differences of the expression patterns of the different nourishing tissues, a data frame with expression information for 28723 OGs identified by OrthoFinder for all eight species was assembled. Due to the low complexity of the Ginkgo transcriptomes these were excluded from this analysis. A PCA was constructed using PCAtools (Blighe *et al.*, 2023). Clades were defined as Gymnosperms represented by *Pinus*, Eudicots with

*Arabidopsis*, *Mimulus* and *Solanum*, early divergent angiosperms with *Amborella* and *Nymphaea* and Monocots with *Oryza* and *Zea*. Eigenvalues were tested for correlation to species, tissue and clade correspondence.

Differential gene expression analyses were performed using DESEQ (Love *et al.*, 2014) separately on each of the species analyzed. It was our aim to identify differentially expressed genes (DEGs) between the somatic tissues, represented by leaves, and the nourishing tissues, represented by endosperm in most angiosperms. For pine and ginkgo, the nourishing tissue is the megagametophyte, whereas for *Nymphaea,* the transcriptomes of perisperms were included in the analysis. Additionally, for *Nymphaea* we performed DGE analyses between perisperms and endosperms. Differentially expressed genes were defined as those with > 2 absolute fold changes and <0,01 corrected p-values. Full list of DEGs per species and DE contrast are defined in Table S3. Due to the poor functional annotation of the *Nymphaea* genome and with the aim to maximize our inferences, enrichment analyses were performed implementing a hypergeometric test of PFAM terms using R (version 4.2.0) with the 'stats' package.

In order to identify functions that define the nourishing function we refrained our subsequent analyses to those genes that were activated, i.e. upregulated in those tissues. To identify conserved pathways across the different plant lineages studied, sets of DEGs per species were translated to OG assignments, resulting in differentially expressed orthogroups (DEOGs). A single OG may be composed of a set of related genes in each species, therefore the number of DEGs in nourishing tissues is always higher than the number of corresponding OGs (Table S1). Different set operations were implemented to uncover trends of expression in the different nourishing tissues. Intersections were used to detect overlaps, and side complements were used to detect DEOGs that were exclusive of specific clades. UpSetR (Conway *et al.*, 2017) and nVennR (Pérez-Silva *et al.*, 2018) were then used to carry out the set operations and recover the GO IDs. We tested the significance of these overlaps by using hypergeometric tests using R (version 4.2.0) with the 'stats' package. We used all the OGs found differentially expressed as the universe and the identity of the sets of DE OGs in the seven species studied as factors for the tests. To identify enriched functions in OGs that were identified in multiple taxa, we annotated them with the best BLAST hits in the *Arabidopsis* Araport11 database. These *Arabidopsis* annotations for OGs are an attempt to bring to the reader an understanding of the biological function of the OG, but these represent the set of genes from the species used in this study, which descended from a single gene in the last common ancestor of our species set (Emms & Kelly, 2019). Using *Arabidopsis* identifiers and with the aim of identifying sets of related DEGs, we drew an interaction network in STRING (Szklarczyk *et al.*, 2017), with only OGs for which there is evidence for functional and physical interactions. Using the same approach on STRING, enrichment analyses of the full set of OGs expressed in nourishing tissues were performed per target clade separately. To test for enrichment, a gene universe was delimited consisting of all the OGs that were found expressed on the nourishing tissues corresponding to the species sets used in the test. All enriched terms reported are significant to an FDR corrected p-value <0,05. To separate the sets of shared DGEs into genes with discrete functions, an MCL clustering algorithm with a 1.5 inflation parameter was implemented in

STRING. In order to describe the expression patterns of clusters of OGs of importance heatmaps were drawn with the complexHeatmap package in R (Gu, 2022) including details of taxa specific annotation of the OGs.

### References


**Batista RA, Moreno-Romero J, Qiu Y, van Boven J, Santos-González J, Figueiredo DD, Köhler C**. **2019**. The mads-box transcription factor pheres1 controls imprinting in the endosperm by binding to domesticated transposons. *eLife* **8**.

**Blighe K, Brown A-L, Carey V, Hooiveld G, Lun A**. **2023**. PCAtools: PCAtools: Everything Principal Components Analysis.

**Cao J-Y, Xu Y-P, Cai X-Z**. **2020**. Integrated miRNAome and Transcriptome Analysis Reveals Argonaute 2-Mediated Defense Responses Against the Devastating Phytopathogen Sclerotinia sclerotiorum. *Frontiers in Plant Science* **11**: 500.

**Chen C, Li T, Zhu S, Liu Z, Shi Z, Zheng X, Chen R, Huang J, Shen Y, Luo S, *et al.*2018**. Characterization of Imprinted Genes in Rice Reveals Conservation of Regulation and Imprinting with Other Plant Species. *Plant Physiology* **177**: 1754–1771.

**Conway JR, Lex A, Gehlenborg N**. **2017**. UpSetR: an R package for the visualization of intersecting sets and their properties. *Bioinformatics* **33**: 2938–2940.

**Emms DM, Kelly S**. **2019**. OrthoFinder: Phylogenetic orthology inference for comparative genomics. *Genome Biology* **20**.

**Florez-Rueda AM, Grossniklaus U, Schmidt A**. **2016a**. Laser-assisted Microdissection (LAM) as a tool for transcriptional profiling of individual cell types. *Journal of Visualized Experiments*: e53916.

**Florez-Rueda AM, Paris M, Schmidt A, Widmer A, Grossniklaus U, Städler T**. **2016b**. Genomic imprinting in the endosperm is systematically perturbed in abortive hybrid tomato seeds. *Molecular Biology and Evolution* **33**: 2935–2946.

**Gao M, He Y, Yin X, Zhong X, Yan B, Wu Y, Chen J, Li X, Zhai K, Huang Y, *et al.*2021**. Ca2+ sensor-mediated ROS scavenging suppresses rice immunity and is exploited by a fungal effector. *Cell* **184**: 5391-5404.e17.

**Gu Z**. **2022**. Complex heatmap visualization. *iMeta* **1**: e43.

**Kim D, Langmead B, Salzberg SL**. **2015**. HISAT: A fast spliced aligner with low memory requirements. *Nature Methods* **12**: 357–360.

**Kinser TJ, Smith RD, Lawrence AH, Cooley AM, Vallejo-Marín M, Conradi Smith GD, Puzey JR**. **2021**. Endosperm-based incompatibilities in hybrid monkeyflowers. *The Plant Cell* **33**: 2235–2257.

**Love MI, Huber W, Anders S**. **2014**. Moderated estimation of fold change and dispersion for RNA-seq data with DESeq2. *Genome Biology* **15**: 550.

**Moreno-Romero J, De León GDT, Yadav VK, Santos-González J, Köhler C**. **2019**. Epigenetic signatures associated with imprinted paternally-expressed genes in the Arabidopsis endosperm. *Genome Biology* **20**: 41.

**Pérez-Silva JG, Araujo-Voces M, Quesada V**. **2018**. nVenn: generalized, quasi-proportional Venn and Euler diagrams. *Bioinformatics* **34**: 2322–2324.

**Pertea M, Pertea GM, Antonescu CM, Chang T-C, Mendell JT, Salzberg SL**. **2015**. StringTie enables improved reconstruction of a transcriptome from RNA-seq reads. *Nature biotechnology* **33**: 290–295.

**Rodrigues AS, Chaves I, Costa BV, Lin YC, Lopes S, Milhinhos A, Van de Peer Y, Miguel CM**. **2019**. Small RNA profiling in Pinus pinaster reveals the transcriptome of developing seeds and highlights differences between zygotic and somatic embryos. *Scientific Reports* **9**: 1–14.

**Szklarczyk D, Morris JH, Cook H, Kuhn M, Wyder S, Simonovic M, Santos A, Doncheva NT, Roth A, Bork P, *et al.*2017**. The STRING database in 2017: Quality-controlled protein-protein association networks, made broadly accessible. *Nucleic Acids Research* **45**: D362–D368.

**Wang B, Regulski M, Tseng E, Olson A, Goodwin S, McCombie WR, Ware D**. **2018a**. A comparative transcriptional landscape of maize and sorghum obtained by single-molecule sequencing. *Genome Research* **28**: 921–932.

**Wang Q, Wang M, Chen J, Qi W, Lai J, Ma Z, Song R**. **2022**. ENB1 encodes a cellulose synthase 5 that directs synthesis of cell wall ingrowths in maize basal endosperm transfer cells. *The Plant Cell* **34**: 1054–1074.

**Wang L, Yuan J, Ma Y, Jiao W, Ye W, Yang DL, Yi C, Chen ZJ**. **2018b**. Rice Interploidy Crosses Disrupt Epigenetic Regulation, Gene Expression, and Seed Development. *Molecular Plant* **11**: 300–314.

**Yang T, Guo L, Ji C, Wang H, Wang J, Zheng X, Xiao Q, Wu Y**. **2020**. The B3 domain-containing transcription factor ZmABI19 coordinates expression of key factors required for maize seed development and grain filling. *The Plant Cell* **33**: 104–128.

**Zuellig MP, Sweigart AL**. **2018**. Gene duplicates cause hybrid lethality between sympatric species of Mimulus. *PLOS Genetics* **14**: e1007130.

**Table S1 Data provenance, tissue details, number of total mapped reads, total number of genes detected, DEGs and DEOGs.**

| *species* | *tissue* | *source* | *reference* | *mapped reads* | *genes assessed* | *total DGEs* | *up DGEs* | *DGEOs* |
|---|---|---|---|---|---|---|---|---|
| *Arabidopsis* | endosperm 6 days | SRR8353381 | Batista *et al.* 2019 | 135988095 | 38850/41671 | 8775 | 4697 | 2264 |
| | endosperm 6 days | SRR8353382 | Batista *et al.* 2019 | 67261111 | | | | |
| | endosperm 6 days | SRR8353383 | Moreno Romero *et al.* 2019 | 63098287 | | | | |
| | endosperm 4 days | SRR7825979 | Moreno Romero *et al* 2019 | 29386934 | | | | |
| | endosperm 4 days | SRR7825980 | Moreno Romero *et al* 2019 | 38654315 | | | | |
| | leave | SRR11128223 | Cao *et al.* 2020 | 37479628 | | | | |
| | leave | SRR11128224 | Cao *et al.* 2020 | 43280855 | | | | |
| | leave | SRR11128225 | Cao *et al.* 2020 | 40706150 | | | | |
| | rossette leaves | SRR13291162 | PRJNA686693 | 60865532 | | | | |
| | rossette leaves | SRR13291163 | PRJNA686693 | 60614678 | | | | |
| *Solanum* | young leaves | SRR3675045 | Florez Rueda *et al.* 2016 | 24188840 | 26269/34688 | 8557 | 3909 | 2172 |
| | young leaves | SRR3675016 | Florez Rueda *et al.* 2016 | 72264034 | | | | |
| | endosperm 14 days | SRR3674959 | Florez Rueda *et al.* 2016 | 23862014 | | | | |
| | endosperm 14 days | SRR3674974 | Florez Rueda *et al.* 2016 | 62184166 | | | | |
| | endosperm 14 days | SRR3674975 | Florez Rueda *et al.* 2016 | 68511566 | | | | |
| | endosperm 14 days | SRR3674976 | Florez Rueda *et al.* 2016 | 61538546 | | | | |
| *Mimulus gutatus* | leaves | SRR6391490 | Zuellig *et al.* 2018 | 31316180 | 22337/28140 | 2407 | 654 | 453 |
| | leaves | SRR6391491 | Zuellig *et al.* 2018 | 38538677 | | | | |
| | leaves | SRR6391492 | Zuellig *et al.* 2018 | 29238533 | | | | |
| | leaves | SRR7976169 | Zuellig *et al.* 2018 | 29416194 | | | | |
| | leaves | SRR7976181 | Zuellig *et al.* 2018 | 29022027 | | | | |
| | leaves | SRR7976189 | Zuellig *et al.* 2018 | 34481348 | | | | |
| | leaves | SRR7976210 | Zuellig *et al.* 2018 | 31238578 | | | | |
| | endosperm 11 days | SRR14078407 | Kinser *et al.* 2021 | 4015137 | | | | |
| | endosperm 11 days | SRR14078408 | Kinser *et al.* 2021 | 35854703 | | | | |
| | endosperm 11 days | SRR14078423 | Kinser *et al.* 2021 | 31780315 | | | | |
| | endosperm 11 days | SRR14078425 | Kinser *et al.* 2021 | 30765703 | | | | |

| *species* | *tissue* | *source* | *reference* | *mapped reads* | *genes assessed* | *total DGEs* | *up DGEs* | *DGEOs* |
|---|---|---|---|---|---|---|---|---|
| *Oryza sativa* | endosperm 6 days | SRR1743008 | Wang *et al.*, 2018 | 85806814 | 29469/38978 | 9027 | 3559 | 2245 |
| | endosperm 6 days | SRR1743009 | Wang *et al.*, 2018 | 109458658 | | | | |
| | endosperm 6 days | SRR7073535 | Chen *et al.* 2018 | 1192982 | | | | |
| | endosperm 6 days | SRR7073537 | Chen *et al.*, 2018 | 41814234 | | | | |
| | endosperm 6 days | SRR7073539 | Chen *et al.*, 2018 | 47975215 | | | | |
| | leaves | SRR9637049 | PRJNA552367 | 46074003 | | | | |
| | leaves | SRR9637050 | PRJNA552367 | 45299296 | | | | |
| | leaves | SRR9637051 | PRJNA552367 | 50077632 | | | | |
| | leaves | SRR13803768 | Gao *et al.*, 2021 | 46727794 | | | | |
| | leaves | SRR13803770 | Gao *et al.*, 2021 | 45276235 | | | | |
| | leaves | SRR13803771 | Gao *et al.*, 2021 | 3620432 | | | | |
| *Zea mays* | endosperm 20 days | ERR2889276 | Wang *et al.* , 2022 | 69966941 | 33404/44303 | 7184 | 2943 | 1860 |
| | endosperm 20 days | ERR2889277 | Wang *et al* . 2022 | 46768844 | | | | |
| | endosperm 15 days | SRR13171426 | Wang *et al.* 2022 | 62696368 | | | | |
| | endosperm 15 days | SRR13171427 | Wang *et al.* 2022 | 53648804 | | | | |
| | endosperm 16 days | SRR11696189 | Yang *et al.* 2021 | 69365894 | | | | |
| | endosperm 16 days | SRR11696190 | Yang *et al.* 2021 | 60674599 | | | | |
| | leaves | ERR2096662 | Wang *et al.* 2018 | 29865619 | | | | |
| | leaves | ERR2096664 | Wang *et al.* 2018 | 12912806 | | | | |
| | leaves | ERR2096666 | Wang *et al.* 2018 | 42582348 | | | | |
| | leaves | ERR7197486 | PRJEB48470 | 64718832 | | | | |
| | leaves | ERR7197495 | PRJEB48470 | 96894261 | | | | |
| | leaves | ERR7197500 | PRJEB48470 | 134919978 | | | | |
| | leaves | ERR7197509 | PRJEB48470 | 70003551 | | | | |
| *Amborella* | L_83 | ERR4193683 | | 7254364 | 17275/26846 | 7669 | 1317 | 927 |
| | L_84 | ERR4193684 | | 28296302 | | | | |
| | L_85 | ERR4193685 | | 25429495 | | | | |
| | L_86 | ERR4193686 | | 24479406 | | | | |
| | L_87 | ERR4193687 | | 30325147 | | | | |
| | L_88 | ERR4193688 | | 29231791 | | | | |
| | E_135_LCM 2 months | | our own | 17954116 | | | | |
| | E_2_LCM 2 months | | our own | 7371514 | | | | |

| *species* | *tissue* | *source* | *reference* | *mapped reads* | *genes assessed* | *total DGEs* | *up DGEs* | *DGEOs* |
|---|---|---|---|---|---|---|---|---|
| *Nymphaea* | endosperm 18_car1 | 1 week old seeds | our own | 19179797 | 26588 | 4281 | 856 | 634 |
| | endosperm 23_car2 | 1 week old seeds | our own | 47987419 | | | | |
| | endosperm 24_car1 | 1 week old seeds | our own | 41087596 | | | | |
| | endosperm 27_car2 | 1 week old seeds | our own | 30736712 | | | | |
| | endosperm 28_car1 | 1 week old seeds | our own | 18715160 | | | | |
| | endosperm_47 | 1 week old seeds | our own | 50059321 | | | | |
| | perisperm_105 | 1 week old seeds | our own | 46526033 | | | | |
| | perisperm_115 | 1 week old seeds | our own | 14042670 | | | | |
| | perisperm_47 | 1 week old seeds | our own | 108209778 | | | | |
| | Leaves_NL1 | young leaves | our own | 53075534 | | | | |
| | Leaves_Nl2 | young leaves | our own | 31585950 | | | | |
| *Pine* | Leiria leaves | SRR1593617 | Cañas *et al.,* 2015 | 144067197 | 32162/50557 | 10539 | 1948 | 1096 |
| | Leiria leaves | SRR1593618 | Cañas *et al.,* 2015 | 222864501 | | | | |
| | Tamrabta leaves | SRR1593619 | Cañas *et al.* , 2015 | 129548479 | | | | |
| | Tamrabta leaves | SRR1593620 | Cañas *et al.,* 2015 | 52400420 | | | | |
| | Tamrabta leaves | SRR1593621 | Cañas *et al.,* 2015 | 160568861 | | | | |
| | MG_0 1 year old | 1 | our own | 110612101 | | | | |
| | MG_4 1 year old | 2 | our own | 199637168 | | | | |
| | MG_3 1 year old | 3 | our own | 230803674 | | | | |
| | MG_3 1 year old | 4 | our own | 37655075 | | | | |
| | MG_4 1 year old | 6 | our own | 528791186 | | | | |
| | MG_0 1 year old | 7 | our own | 107677504 | | | | |
| *Ginkgo* | GbilobaMG1 | stage 9 | Zumajo-Cardona *et al.* , 2021 | 167257040 | 1549/41309 | 26 | 25 | 15 |
| | GbilobaMG2 | stage 9 | Zumajo-Cardona *et al.,* 2021 | 147481619 | | | | |
| | GbilobaMG | stage 9 | Zumajo-Cardona *et al.,* 2021 | 16677641 | | | | |
| | GinkgoLeaf1A | | Zumajo-Cardona *et al.* , 2021 | 93262307 | | | | |
| | GinkgoLeaf2B | | Zumajo-Cardona *et al.,* 2021 | 83088035 | | | | |
| | GinkgoLeafA | | Zumajo-Cardona *et al.,* 2021 | 113110054 | | | | |

**Table S2 Orthogroup description.** Species gene IDs, gene counts and best Arabidopsis blast hit for each of the 34040 Orthogroups identified. Excel Table provided as a separate file.

**Table S3 – Results of differential gene expression between nourishing tissues and leaves of all species studied.** Gene IDs, DEOG correspondence, normalized counts, DGE statistics and functional annotation. Excel Table provided as a separate file.

**Table S4. Conserved transcriptional network of the angiosperm seed nourishing tissue.** Full details of the 175 OGs conserved in all main angiosperm clades and description of the protein networks inferred. Excel Table provided as a separate file.

**Table S5 – The transcriptional network of the *Pinus pinaster* megagametophyte.** Full details of the 608 OGs conserved in the pine megagametophyte and comparison with the angiosperm conserved OG set. Description of the protein networks inferred. Excel Table provided as a separate file.

**Table S6 – The private transcriptional network of the *Amborella trichopoda* endosperm.** Details of the 437 OGs private of the *Amborella* endosperm and description of the protein network clusters. Excel Table provided as a separate file.

**Table S7 – The transcriptional network of the *Nymphaea caerulea* seed.** Differential gene expression analyses between endosperm and perisperm in the *Nymphaea* seed. Description of the 334 OGs that compose the private transcriptional network of the *Nymphaea* endosperm. Excel Table provided as a separate file.

**Table S8 – The monocot endosperm transcriptional network.** Full details of the 235 OGs conserved in the maize and rice endosperms and not shared with any other taxa studied. Description of the protein networks inferred. Core angiosperm analyses detailing OGs present in monocots and not in eudicots. Excel Table provided as a separate file.

**Table S9 – The transcriptional network of the eudicot endosperm.** Full details of the 187 OGs conserved in the core eudicot taxa studied and not in any other one. Description of the protein networks inferred. Core angiosperm analyses detailing OGs present in eudicots and not in monocots. Excel Table provided as a separate file.

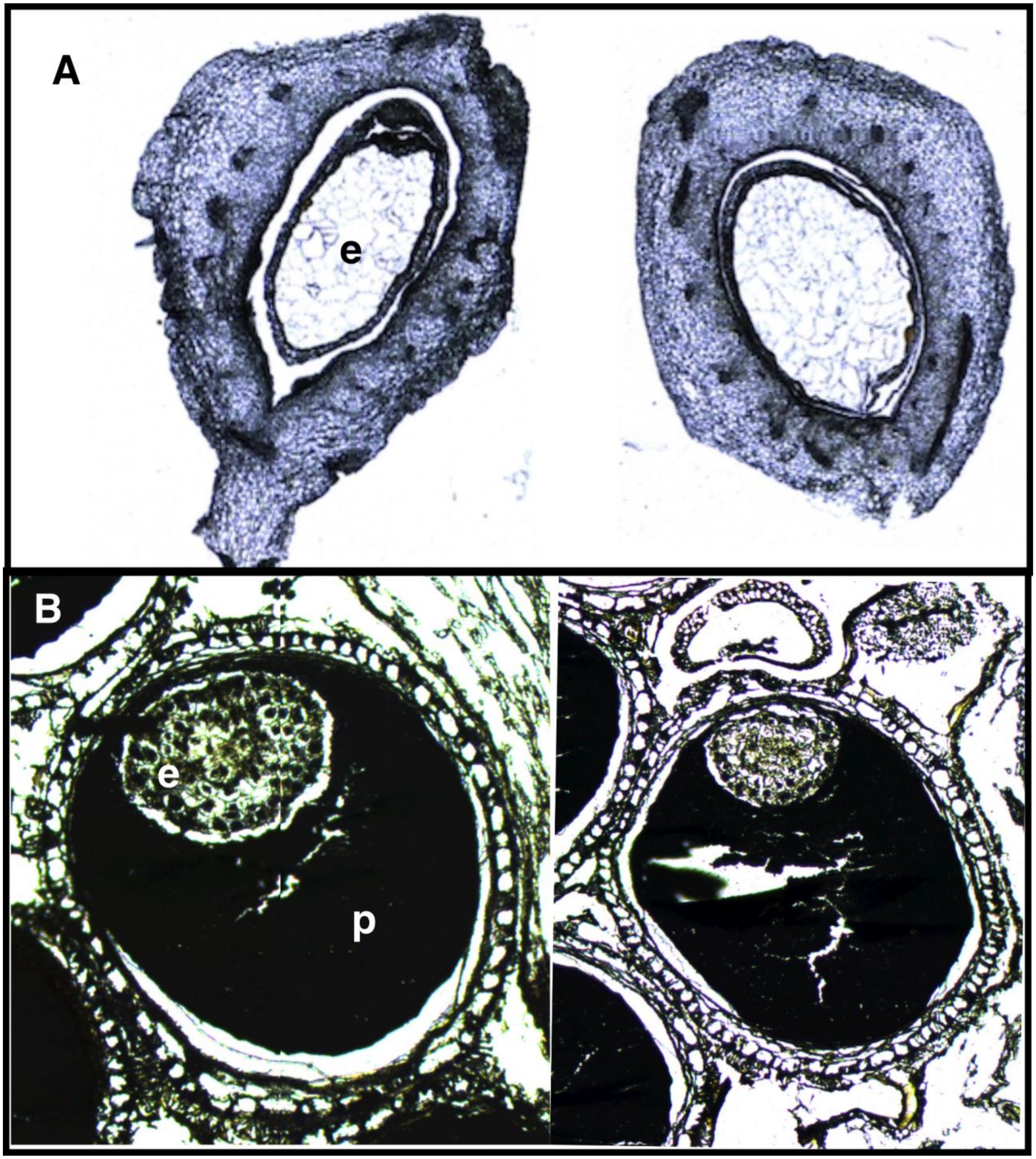


**Fig. S1 Examples of early developing seeds of *Amborella trichopoda* and *Nymphaea caerulea* used for microdissection and transcriptome generation.** A. Example of *Amborella trichopoda* sectioned seeds. No embryos were visible in any of the sectioned seeds. B. *Nymphaea caerulea* seeds upon dissection, endosperm and perisperm were collected. e: endosperm, p: perisperm.

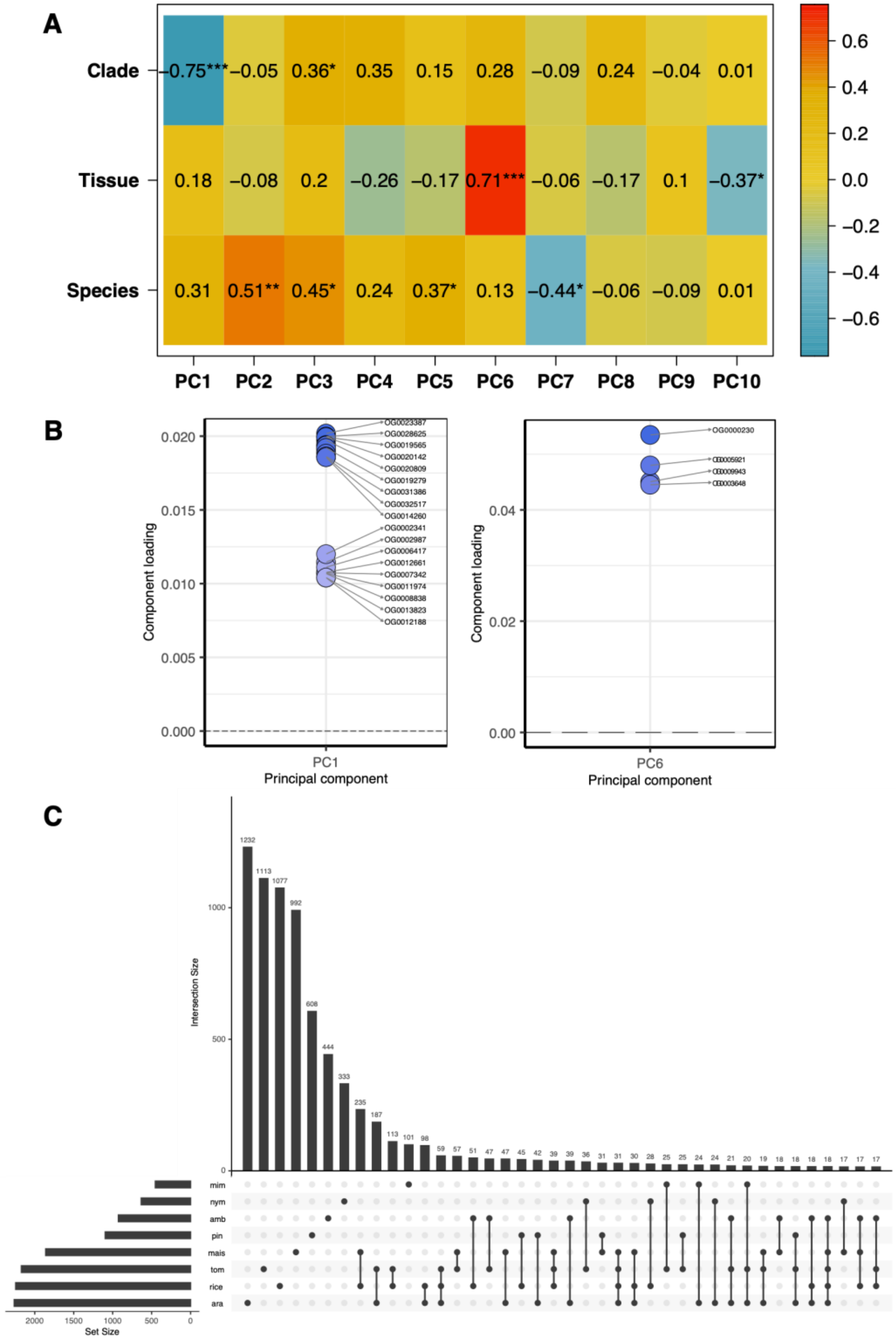


**Fig. S2 Details of PCA analyses and Upset plot of DEOGs showing intersections** A. Eigenvalue correlation plot with tissue clade and species variables. B. Top loadings representing OGs for the PC1 and PC6. C.Upset plot displaying the sets of DEOGs in nourishing tissues of target plant species, showing the largest unique and overlapping sets among different plant clades.

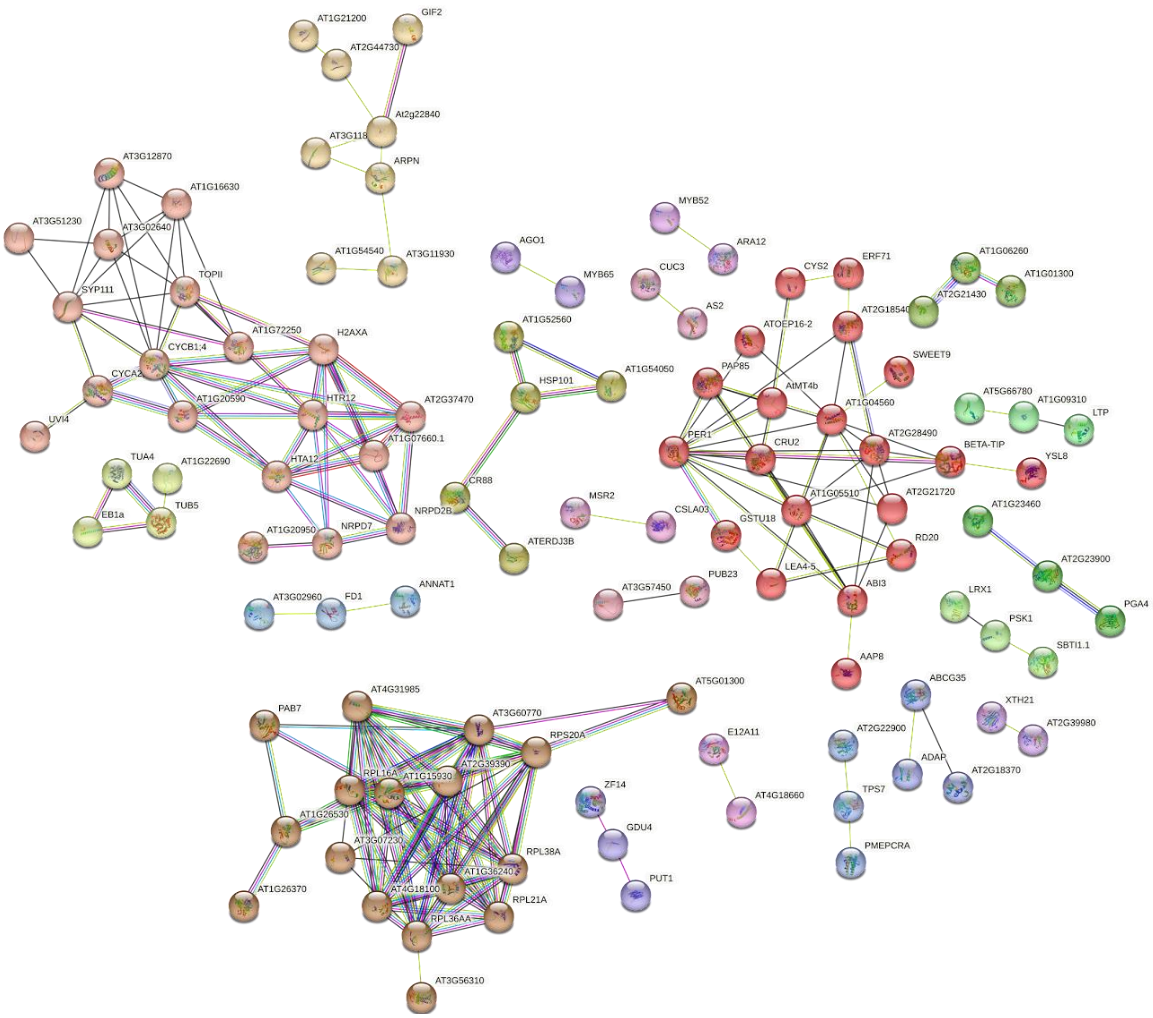


**Figure S3. The conserved transcriptional network of the angiosperm endosperm.** Complete details of orthogroup correspondence and putative protein annotation are found in Table S4.

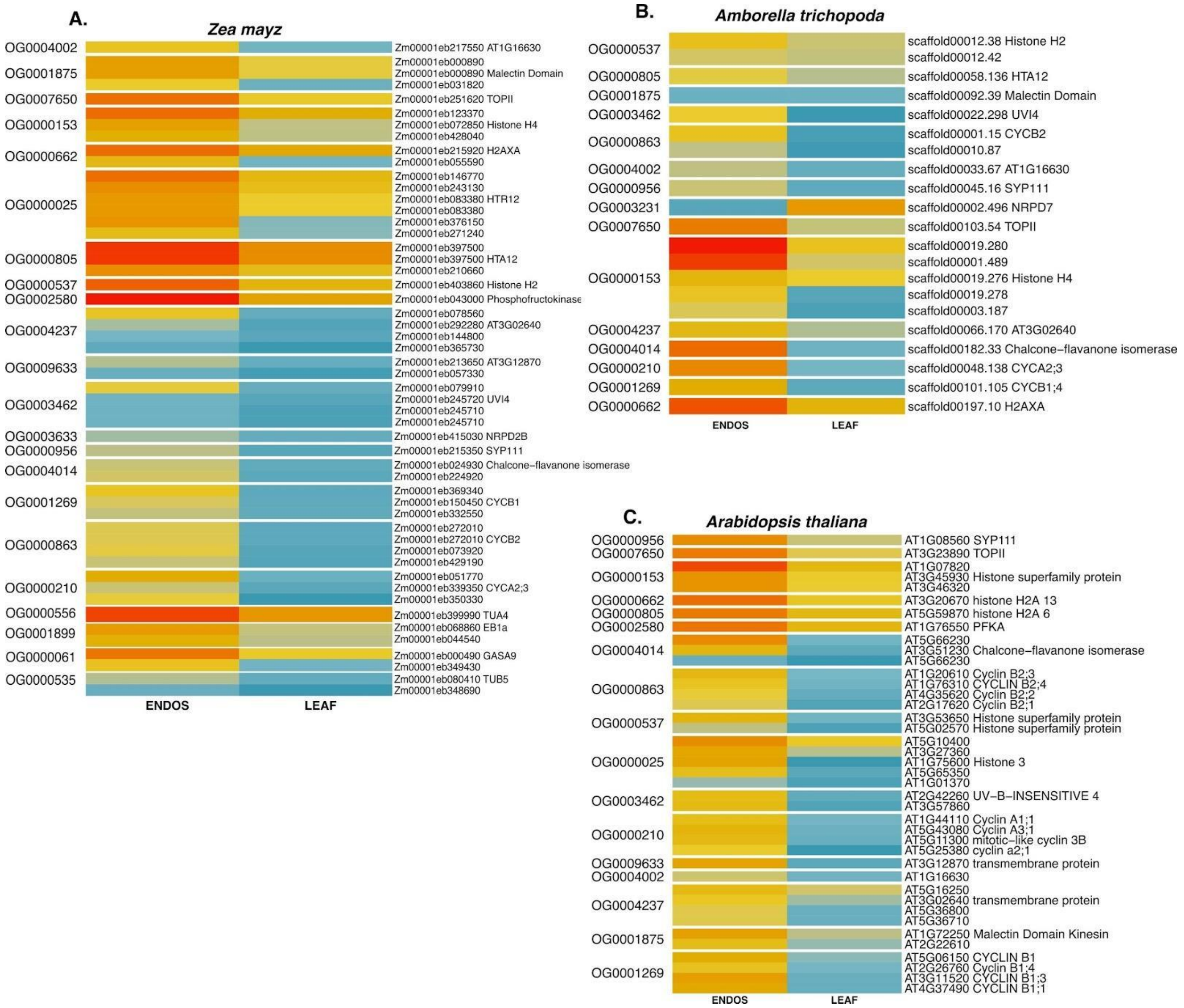


**Figure S4. The conserved transcriptional network of the angiosperm endosperm, the cell cycle and nucleosome architecture interacting network** Orthogroup to DEG correspondence and expression values in CPM for A. *Zea mays*. B. *Amborella trichopoda* and C. *Arabidopsis thaliana.*

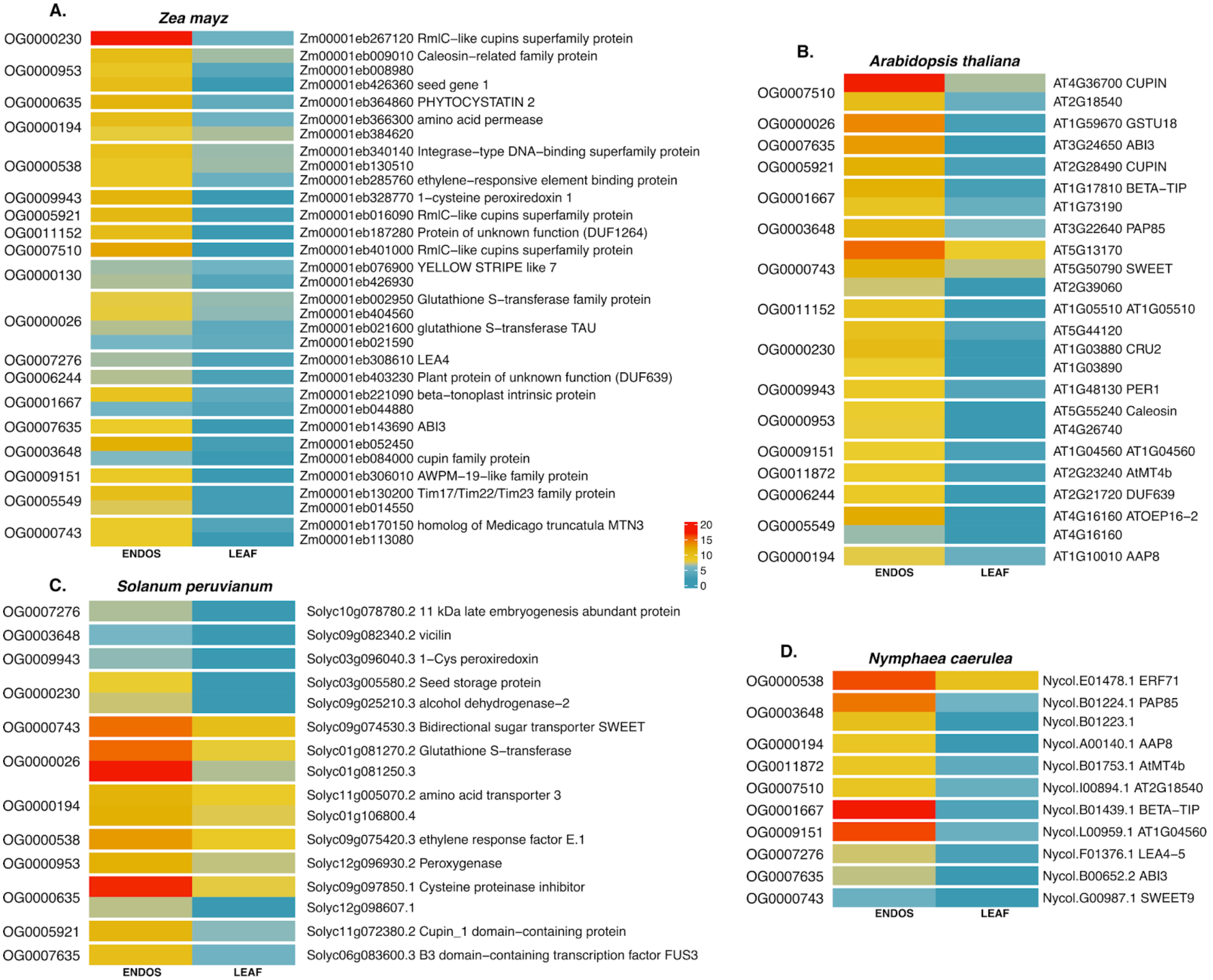


**Figure S5. Conserved transcriptional network of the angiosperm seed nourishing tissue, the Nourishing cluster protein interacting network** Orthogroup to DGE correspondence and expression values in CPM for A. *Zea mays.* B. *Arabidopsis thaliana*. C. *Solanum peruvianum* and D, *Nymphaea caerulea*.

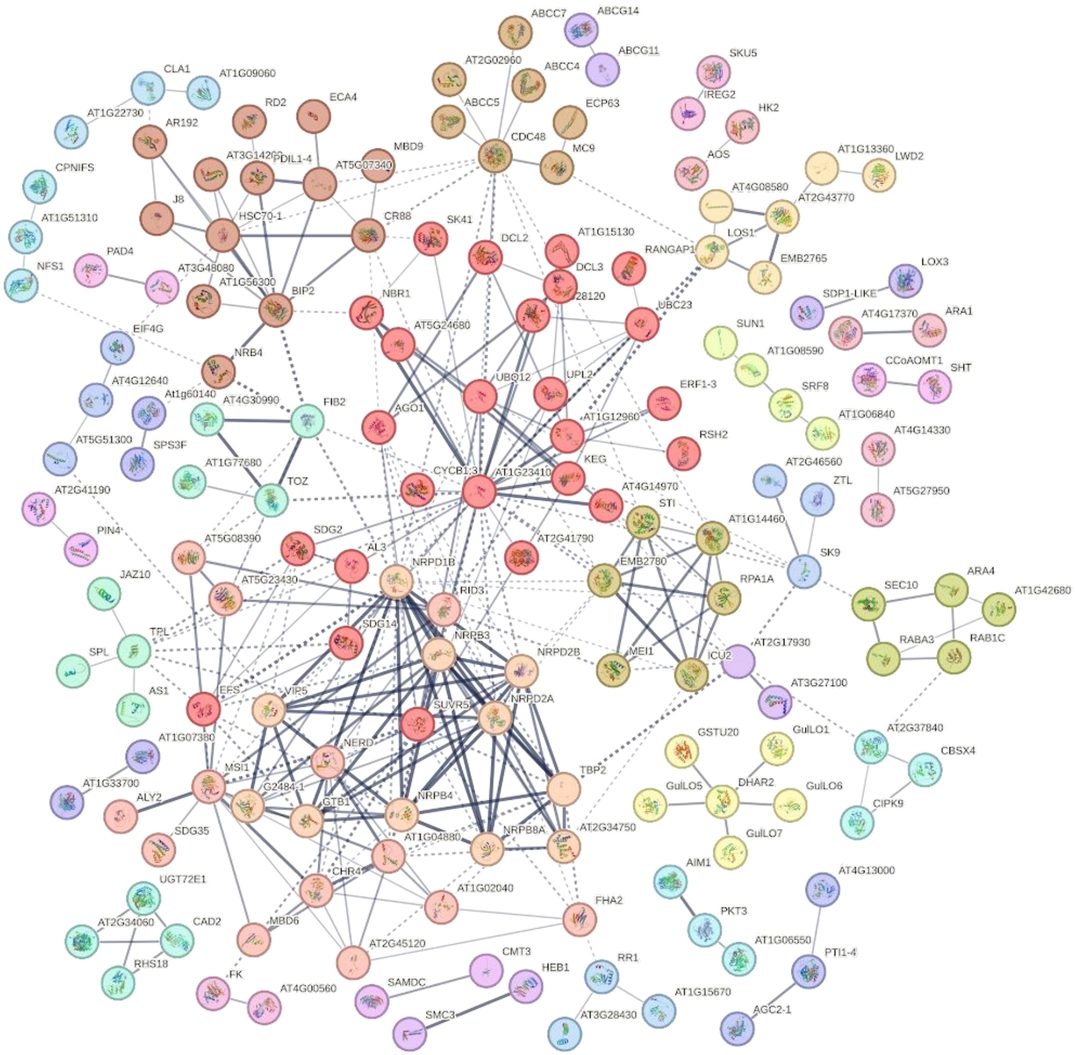


**Figure S6. The *Pinus pinaster* megagametophyte private protein network.** Complete details of orthogroup correspondence and putative protein annotation are found in Table S5.

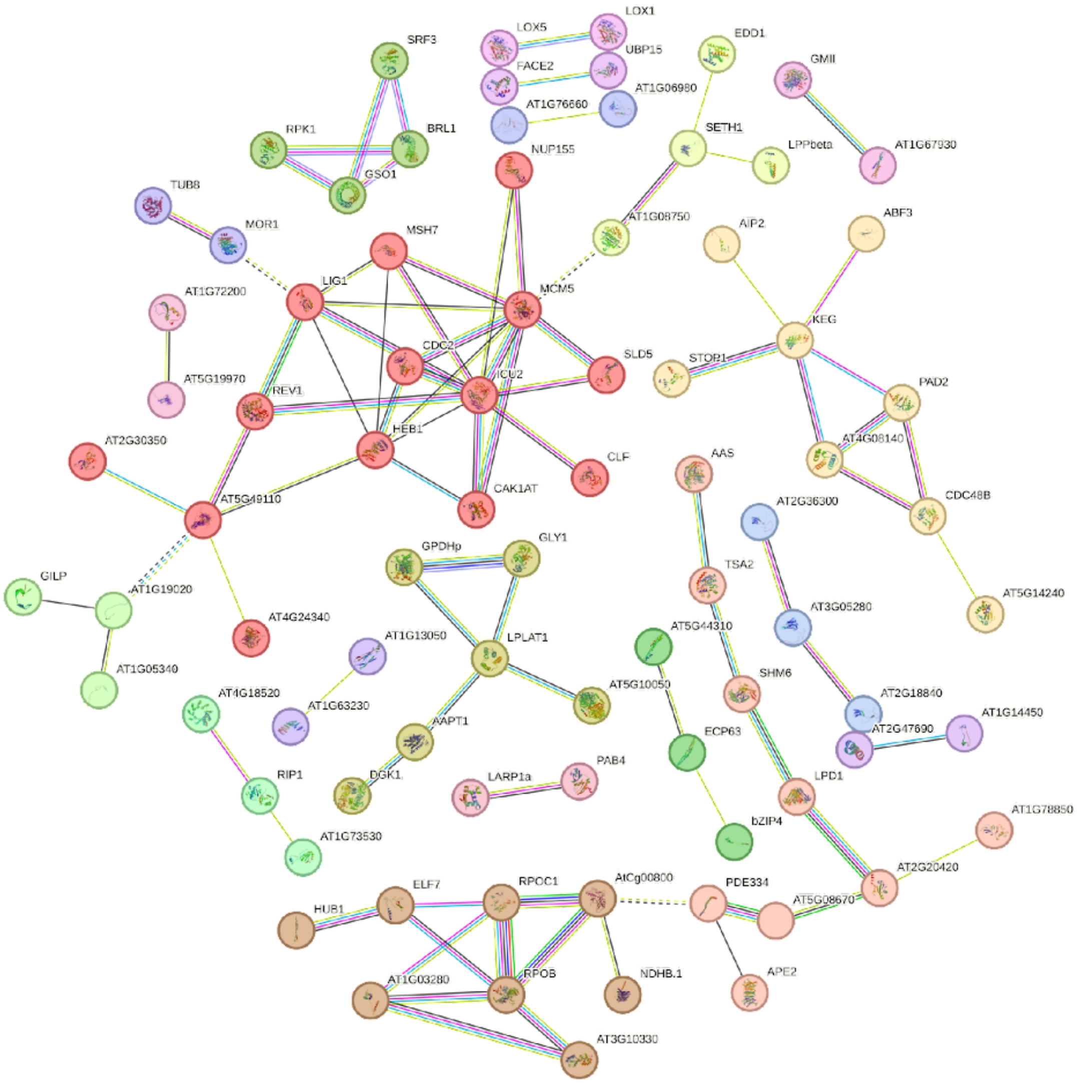


**Figure S7 The *Amborella* endosperm private protein network.** Complete details of orthogroup correspondence and putative protein annotation are found in Table S6.

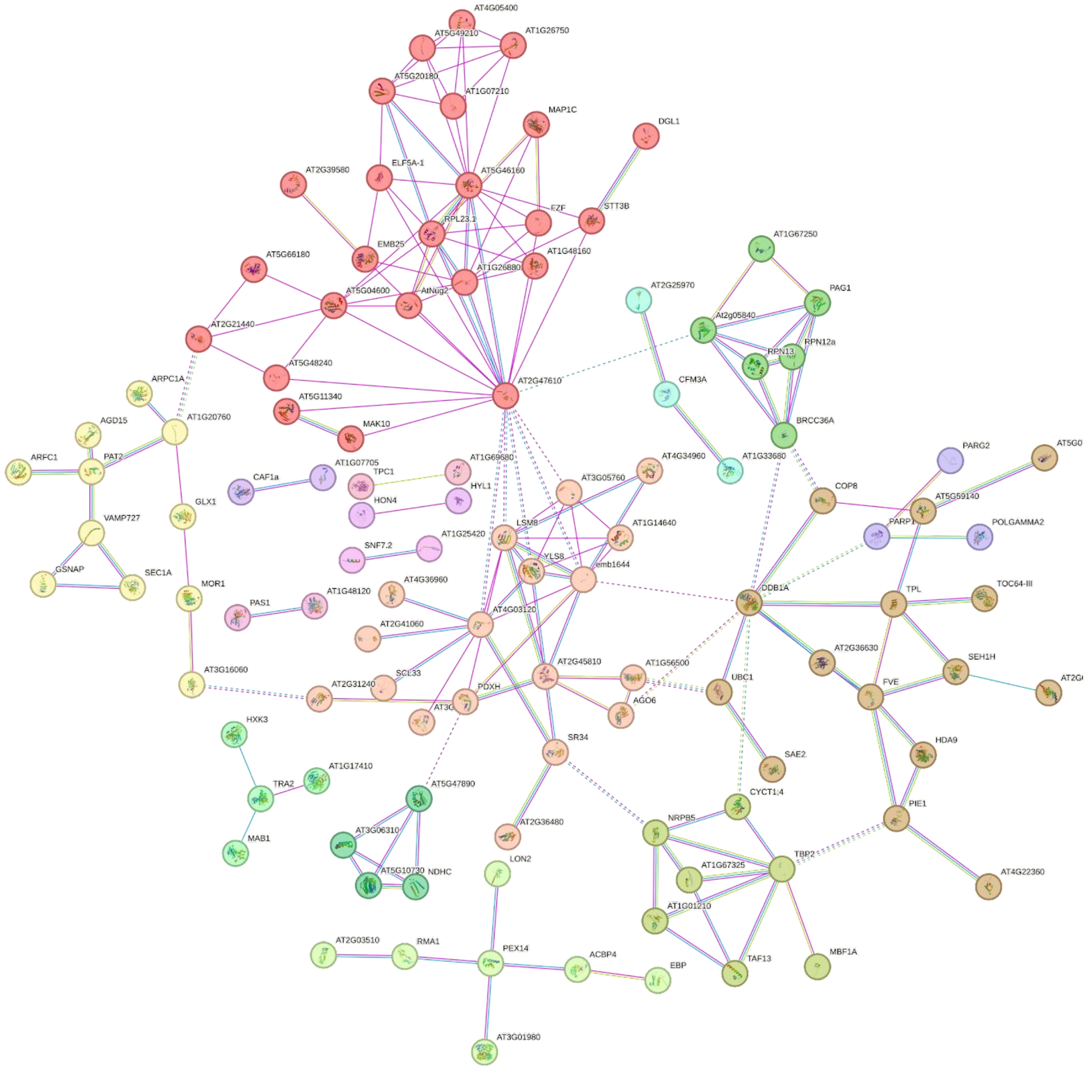


**Figure S8. The *Nymphaea caerulea* endosperm private protein network.** Complete details of orthogroup correspondence and putative protein annotation are found in Table S7.

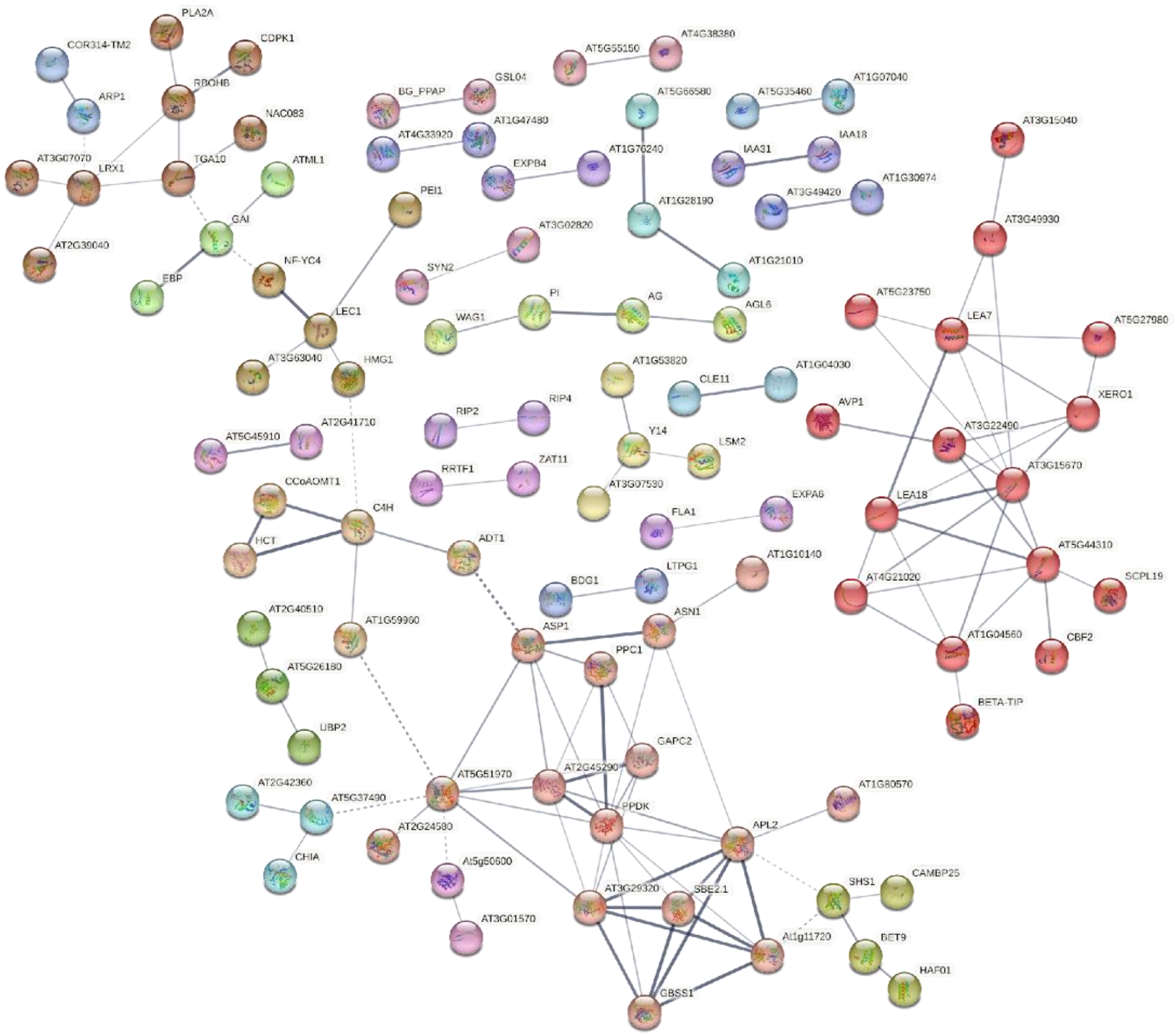


**Figure S9. The monocot endosperm private protein network.** Complete details of orthogroup correspondence and putative protein annotation are found in Table S8.

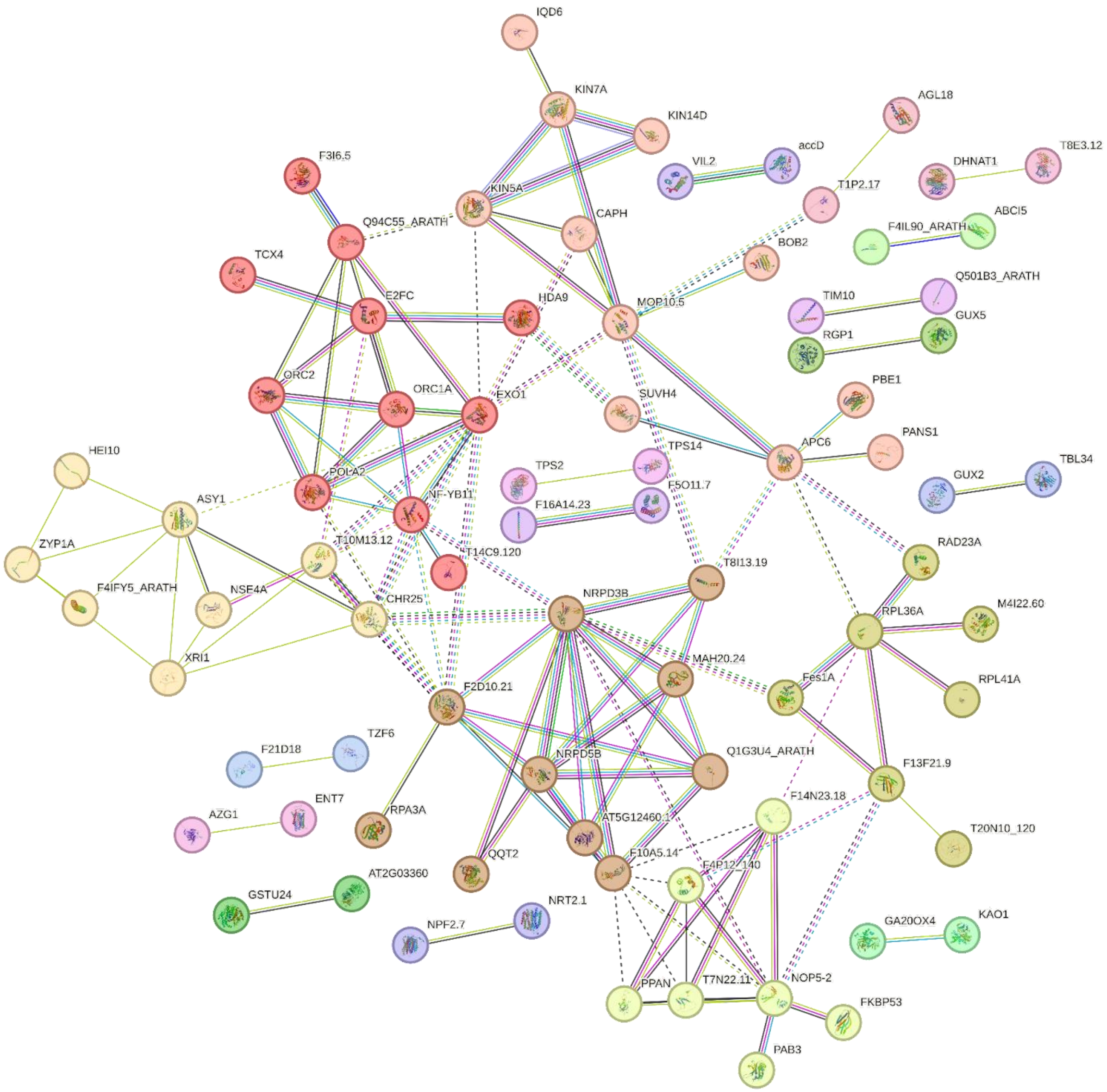


**Figure S10. The eudicot endosperm private protein network.** Complete details of orthogroup correspondence and putative protein annotation are found in Table S9.
